# Centripetal integration of past events by hippocampal astrocytes and its regulation by the locus coeruleus

**DOI:** 10.1101/2022.08.16.504030

**Authors:** Peter Rupprecht, Sian N Duss, Denise Becker, Christopher M Lewis, Johannes Bohacek, Fritjof Helmchen

**Affiliations:** Laboratory of Neural Circuit Dynamics, Brain Research Institute, University of Zurich, Switzerland; Neuroscience Center Zurich, University of Zurich and ETH Zurich, Switzerland; Laboratory of Molecular and Behavioral Neuroscience, Institute for Neuroscience, Department of Health Sciences and Technology, ETH Zurich, Switzerland; University Research Priority Program (URPP), Adaptive Brain Circuits in Development and Learning, University of Zurich, Switzerland

## Abstract

An essential feature of neurons is their ability to centrally integrate information from their dendrites. The activity of astrocytes, in contrast, has been described as mostly uncoordinated across cellular compartments without clear central integration. Here, we describe conditional centripetal integration as a principle of how astrocytes integrate calcium signals from their distal processes to induce somatic activation. We found in mouse hippocampus that global astrocytic activity, as recorded with population calcium imaging, is well explained as a leaky integration of past neuronal and behavioral events on a timescale of seconds. Salient past events, indicated by pupil dilations, facilitated propagation of calcium signals from distal processes to the soma on this slow timescale. Centripetal propagation was reproduced by optogenetic activation of the locus coeruleus, a key regulator of arousal, and reduced by pharmacological inhibition of α1-adrenergic receptors. Together, our results establish astrocytes as computational units of the brain that slowly and conditionally integrate calcium signals to activate their somata upon behaviorally relevant events.

## Introduction

Astrocytes have long been thought to play a supportive rather than computational role in the brain. This picture started to change when the close interaction of astrocytic processes and neuronal synapses was discovered, leading to the concept of the tripartite synapse^1,2^. More recent studies have begun to establish a role for astrocytes, as for neurons, in diverse computational processes of the brain^3,4^. For example, calcium signals in astrocytes have been suggested to represent sensory-related or internally generated information^5–8^. In contrast to neurons, however, information processing in single astrocytes is thought to rely on distributed and mostly uncoordinated activity patterns across the compartments of a single astrocyte, without central integration^9–11^. It is therefore not clear whether and how astrocytes integrate signals that are sensed by their distributed compartments.

The study of signal integration in astrocytes is particularly challenging since they express a large set of receptors that enables them to sense the direct and indirect effects of both neuronal activity and neuromodulatory signals^9,12,13^, all of which interact with behavior. Recent work from systems neuroscience has demonstrated that spontaneous behaviors and motor activity drive a large fraction of the variability of neuronal activity in the mouse cortex^14,15^. These studies highlight the importance of a systematic investigation of possibly interdependent observables in the brain to understand potential confounds between task-related behaviors, spontaneous behaviors, cellular activity patterns, and brain states. It is therefore essential to study astrocytic activity together with factors that have been shown to correlate with astrocytic activity, *e*.*g*., neuromodulatory signals^7,16,17^, pupil diameter^18^, locomotion^17,19–21^ and neuronal activity^6,10,11,22,23^. Moreover, it is crucial to carefully analyze how astrocytic activity reflects these factors, both temporally, in a population of astrocytes, and spatially, within individual astrocytes.

Here, we perform a systematic exploration of astrocytic activity in the hippocampal subregion CA1 in behaving mice using two-photon calcium imaging of astrocytic populations. Using an unbiased approach, we find that global population activity can be described as a temporal integration of past behavioral events such as unexpected air puffs or self-generated body motion. On the single-astrocyte level, the global activity pattern proceeded as spatial propagation of calcium signals from distal to somatic compartments. Spontaneous behavior analysis as well as perturbations based on optogenetics and pharmacology suggest that this centripetal propagation is gated by noradrenaline-release from the locus coeruleus (LC), a brainstem nucleus that is key for arousal effects. Together, our observations reveal a novel principle of spatiotemporal integration of salient past events within astrocytes in the awake, behaving animal.

## Results

### A global astrocytic activity mode during behavior

To record astrocytic activity in the hippocampus across a wide range of behaviors, we virally induced expression of GCaMP6s (n = 6 mice; AAV9-GFAP-GCaMP6s; Fig. S1) and performed two-photon calcium imaging in head-fixed mice that were free to run on a treadmill and received water rewards at a defined location of the treadmill, resulting in variable behavior including periods of active running and quiet wakefulness (Fig. 1a). First, we analyzed calcium transients in the astrocyte population using a single imaging plane in *stratum oriens* of CA1 (Fig. 1b-e; Movie 1). Across active regions of interest (ROIs, Fig. S2), astrocytic calcium signals were highly correlated (Fig. 1f-i). Therefore, the global astrocytic activity, which we define as the average fluorescence trace across the population of astrocytes in the entire field of view (FOV), explained a large fraction of the variance of single astrocyte activity (Fig. 1h,i; correlation: 0.72 ± 0.20, median ± s.d. across 204,686 pairs of astrocytes from 41 imaging session from n = 6 mice). The correlation of activity only decayed slightly with the distance between astrocyte pairs (Fig. 1j), indicating a global rather than local synchronization within hippocampal CA1. In addition, we observed local events in single astrocytes independent of global activity (white arrows in Fig. 1f) and local modulations of global activity in single astrocytic ROIs (black arrows in Fig. 1g; Movie 1). However, the overall spontaneous activity during behavior was dominated by a global mode across astrocytes.

**Figure 1.**
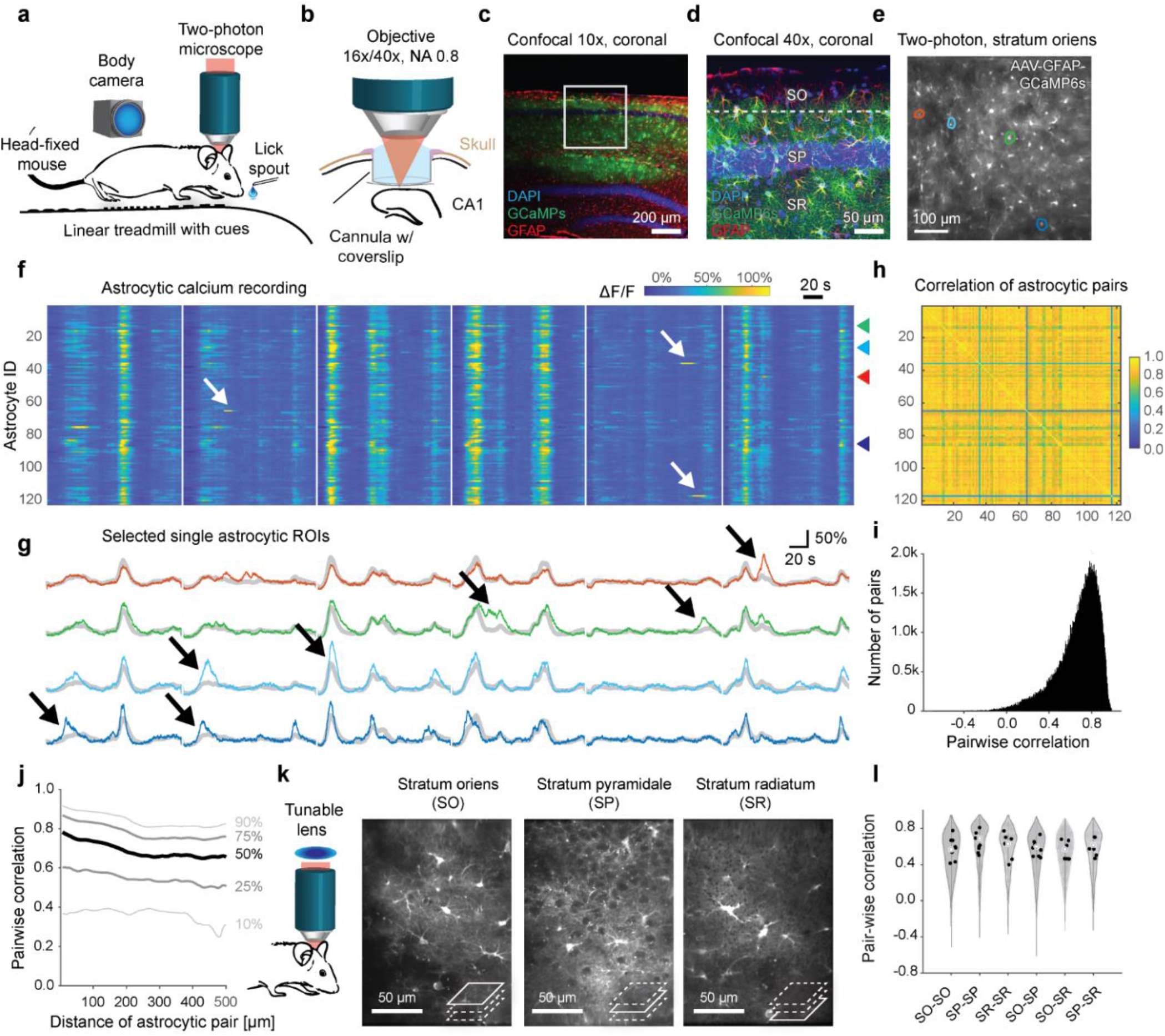
Hippocampal astrocytes in CA1 exhibit global and local events during behavior. **a**, Scheme of the in vivo recording setup. **b**, Hippocampal two-photon imaging through an implanted cannula. **c-d**, Histology of virus-induced GcaMP6s expression in hippocampal astrocytes (green) together with GFAP-antibody staining (red) and nuclear stain (blue). See also Figs. S1 and S2. **e**, Average fluorescence two-photon image of astrocytes in the *stratum oriens* expressing GcaMP6s. **f**, Temporal calcium dynamics of active astrocyte ROIs from the FOV shown in panel (e). White arrows indicate isolated local calcium events. Recording segments (140 s) are indicated through vertical white spacers. **g**, Example of four astrocytic ROIs (highlighted with matching colors in (i) and in (f) with arrow heads), indicating local modulation of global events for astrocytic ROIs (black arrows). Global mean across the FOV is overlaid as grey traces. **h**, Activity correlation between the astrocytic pairs from panel (f), same ordering of ROIs. **I**, Distribution of activity correlations between astrocytic active region pairs across the entire population (204,686 astrocyte ROI pairs from 41 experimental sessions and 6 animals; 0.72 ± 0.20, median ± s.d.). **j**, Distance-dependence of pairwise correlations, with the median (50%) and other percentile lines of the distribution shown. **k**, Using a tunable lens to quasi-simultaneously image multiple CA1 layers. **l**, Pairwise correlation across simultaneously imaged astrocyte pairs associated with specific CA1 layers (distributions in the violin plots), and medians across astrocyte pairs for each session (black dots). No significant differences (p > 0.2 for all comparisons) across conditions for session-based testing (n = 8 sessions from 2 animals). ROI, region of interest; FOV, field of view; SO, *stratum oriens*; SP, *stratum pyramidale*; SR, *stratum radiatum*.

### Global astrocytic activity across hippocampal layers

To understand how astrocytic activity was correlated across different depths of hippocampal CA1, we performed triple-layer calcium imaging using fast z-scanning with a tunable lens^24^. With this approach, we were able to image quasi-simultaneously from astrocytes in the *stratum oriens*, the *stratum pyramidale* and the *stratum radiatum* (Fig. 1k; Movie 2). Astrocytic activity was highly correlated across layers (Fig. 1l) and we did not find evidence that astrocytic pairs were less correlated across layers than within layers (p > 0.2 for all comparisons, Wilcoxon’s rank sum test, n = 8 sessions from 2 animals, with 4,738 to 10,432 astrocytic ROI pairs for each comparison). Due to the sparse occurrence of astrocytic somata in the *stratum pyramidale*^25^ and the better imaging access to *stratum oriens*, we focus all the remaining analyses and experiments, unless otherwise stated, on the *stratum oriens* layer of hippocampal CA1.

### Astrocytic activity recorded together with neuronal activity, motor behaviors, and pupil diameter

To probe how astrocytic activity relates to the animals’ behavior and hippocampal neuronal activity, we performed simultaneous calcium imaging of astrocytes in *stratum oriens* (SO) of CA1 (viral expression of GCaMP6s as before) and of neurons (transgenic Thy1-GCaMP6f background, ref. ^26^) in the *stratum pyramidale* (SP), 60-90 μm below the SO-imaging plane (Fig. 2a-c; n = 4 mice, 22 imaging sessions). GCaMP signals from astrocytes and neurons were clearly distinct from each other due to their different location (SO vs. SP) and dynamics (slow vs. fast transients). Within SO, neuronal and glial signals were unmixed (Fig. S3 and Methods for details and validation of this approach). To quantify average neuronal population activity, we extracted ΔF/F traces from ROIs of neuronal somata, denoised these traces using a supervised deconvolution algorithm based on deep networks^27^, and averaged across the denoised spike rates, resulting in an overall hippocampal spike rate (SR) estimate. In addition to calcium imaging, we recorded the animals’ run speed as well as the spatial location on the treadmill. Moreover, we used a video camera to record body movements (Fig. 2d). We estimated the movement of the mouth and the paws using the across-frames correlation in corresponding subareas of the videos to calculate movement strengths, and we quantified licking by detecting the tongue from the behavior videos. Finally, we used a dim focused UV light to constrict the pupil in the otherwise dark environment, which allowed us to record pupil diameter as a proxy for neuromodulatory tone and arousal^28–30^. These multiple perspectives (Fig. 2e; Movie 3) enable a comprehensive analysis of astrocytic activity in the context of diverse, possibly related processes and events.

**Figure 2.**
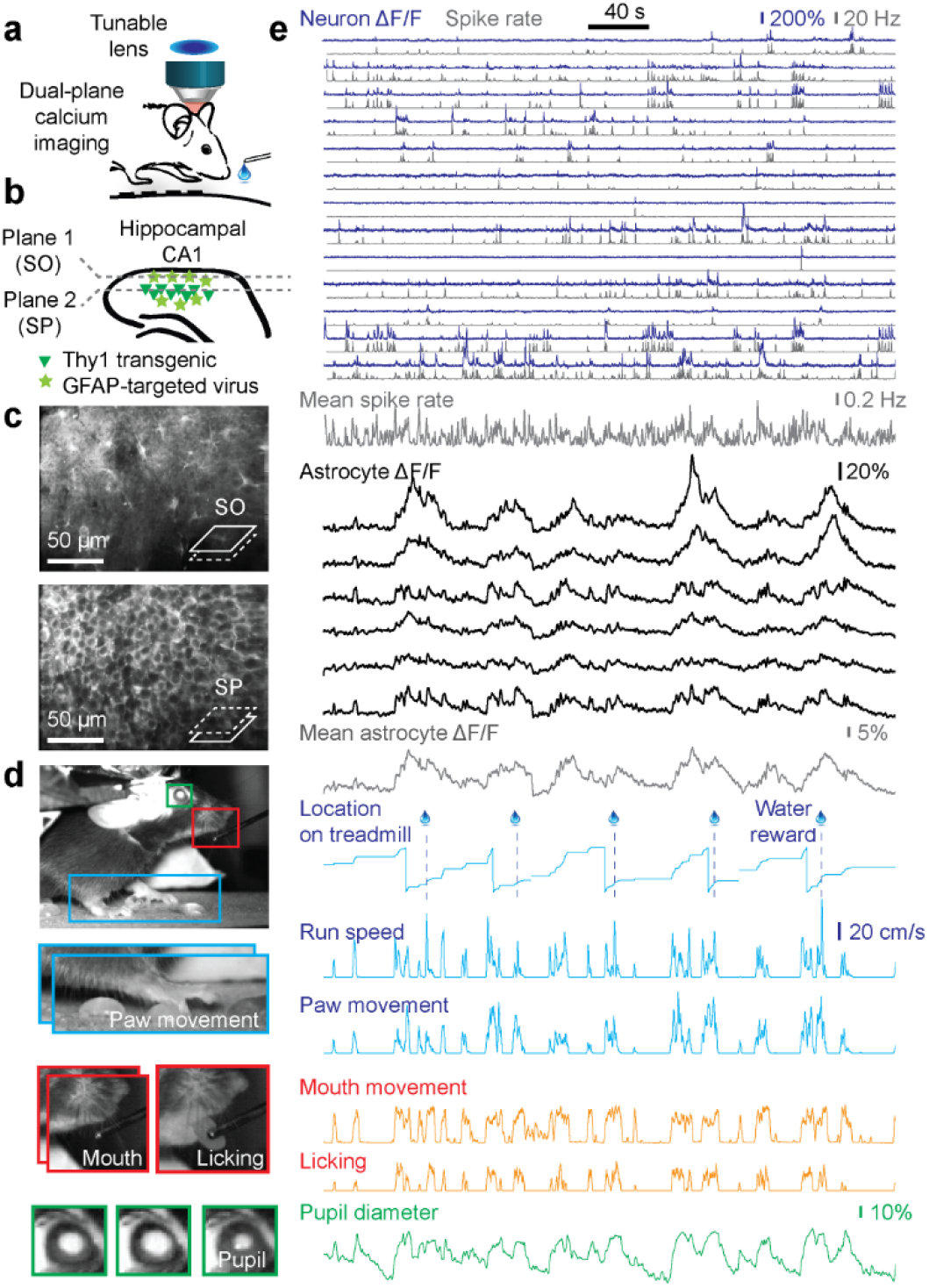
Simultaneous monitoring of astrocytic and neuronal population activity, pupil diameter and behavior. **a**, Dual-plane calcium imaging using a tunable lens. **b**, Simultaneously imaging of spatially separated astrocytes (*stratum oriens* (SO) layer, virally induced GcaMP6s; star symbols) and transgenically expressed neurons (stratum pyramidale (SP) layer, Thy1-GcaMP6f; triangle symbols). **c**, Mean fluorescence image of simultaneously imaged SO layer (with astrocytes) and SP layer (with neurons). **d**, Example behavioral camera image. Blue: Paw movement extracted from subsequent video frames. Red: Mouth movement extracted from subsequent video frames, licking extracted by tongue detection. Green: Pupil diameter visible due to laser light passing from the brain through the eye. **e**, Example of simultaneous recordings, from top to bottom: Subset of neuronal ΔF/F traces (blue, 13 from a total of 107 neurons) extracted from FOV in (c), together with deconvolved spike rates (black) and mean spike rate across all 107 neurons (bottom). Extracted astrocytic ΔF/F traces (6 out of a total of 34 active astrocytic ROIs), together with the mean astrocytic trace across the FOV in (c). Blue: Tracking of the position along the treadmill, together with time points of water rewards; run speed is determined from a rotary encoder, paw movement is extracted from video analysis (blue panel in (d)). Red: Mouth movement and licking are extracted from video analysis (red panels in (d)). Green: Pupil diameter, relative change with respect to median, extracted from video analysis (green panel in (d)).

### Global astrocytic activity can be explained by pupil diameter, body movement or neuronal activity

Due to the slow changes of global astrocytic activity over time (Fig. 1f,g; Fig. 2e), a typical 15-35 min recording only sparsely samples the state space of astrocytic activity. To understand and explain astrocytic activity, we therefore aimed, in a first step, to explain the global mode of astrocytic activity by either run speed, body movements, pupil diameter or mean neuronal spike rate using simple linear models. In a first approximation, we estimated the shared information between global astrocytic activity and these factors using instantaneous correlation (Fig. 3a). Interestingly, we found that pupil diameter displayed the highest correlation with the global astrocytic signal, whereas run speed, body movements and neuronal activity were less correlated (Fig. 3c).

**Figure 3.**
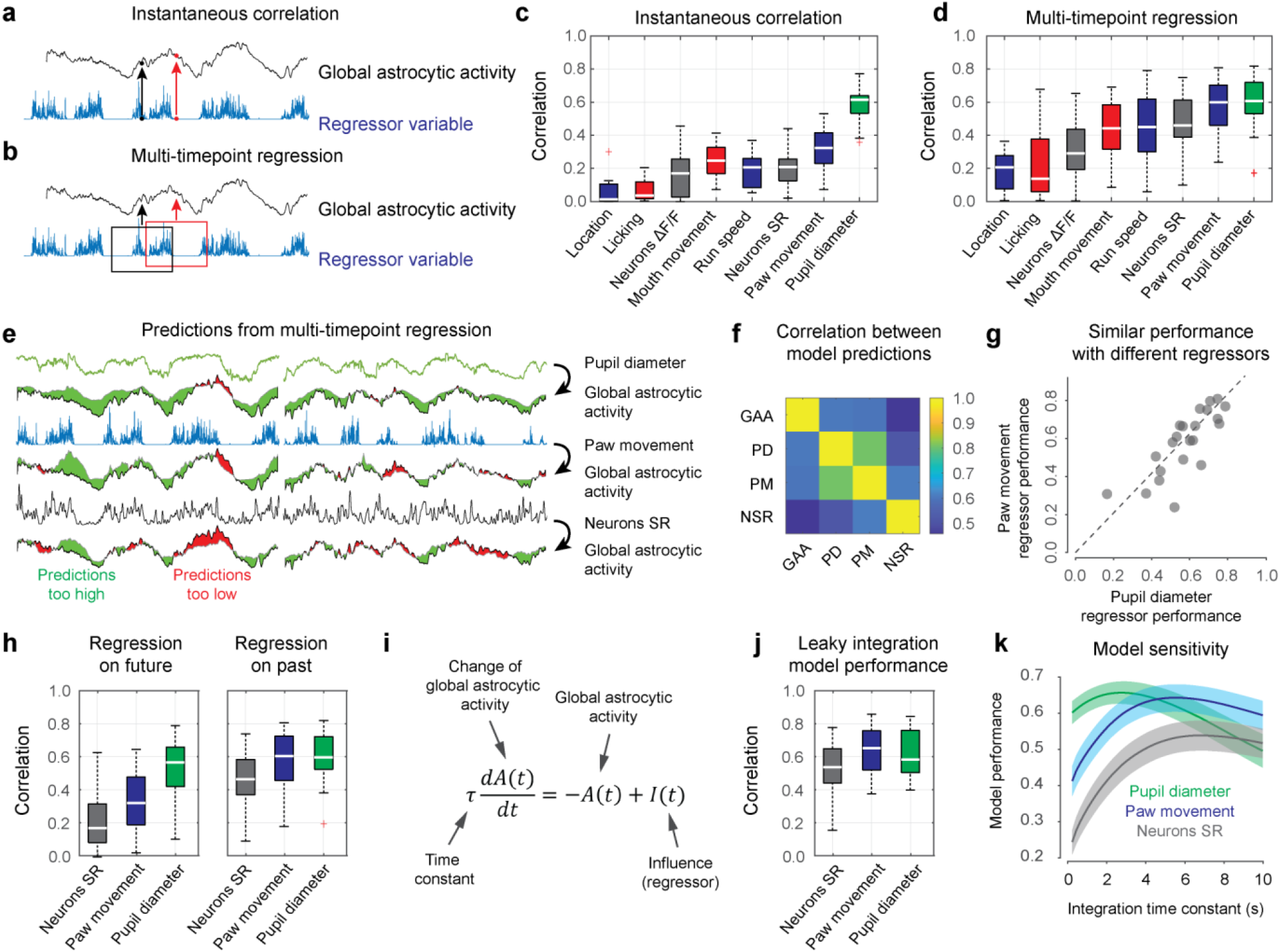
Global astrocytic activity can be well explained by past behavior, mean neuronal spike rate or pupil diameter. **a**, Schematic illustration of instantaneous correlation. A single time point is used to predict a simultaneous astrocytic time point. **b**, Schematic illustration of multi-timepoint regression. A range of data points in a window is used to predict a single time point of astrocytic activity. **c-d**, Performance of various regressors when using instantaneous correlation (c) or multi-timepoint analysis (d) to predict astrocytic activity (n = 22 imaging sessions from 4 animals). The mean neuronal spike rate is abbreviated as ‘Neurons SR’. Box plot properties are defined as described in Methods. **e**, Example cross-validated predictions from the three best regressors (pupil diameter, paw movement, mean neuronal spike rate). Green areas indicate when predicted activity is too high, red areas when it is too low. **f**, Average correlation between global astrocytic activity and the activity predicted by the three regressors (GAA, global astrocytic activity; PD, pupil diameter; PM, paw movement; NSR, neurons spike rate). **g**, Example of performance for two different regressors (pupil diameter, paw movement) across sessions (correlation *r* = 0.79). See also Fig. S6. **h**, Multi-timepoint regression, but using only either future or past time points. **i**, Leaky integration differential equation to model global astrocytic activity A(t) as a function of regressor input I(t), dependent on the integration time constant τ. **j**, Model performance using the leaky integration differential equation. **k**, Model sensitivity with respect to the integration time constant τ (mean ± bootstrapped 90% confidence interval of the mean, determined from 22 imaging sessions).

Next, to account for possible delayed effects that cannot be captured by correlation or other instantaneous measures, we used past and future time points of neuronal and behavioral variables to explain the current value of global astrocytic activity using multi-timepoint, “dilated” linear regression (Fig. 3b; window extent -17…+17 s; see Methods for details). Interestingly, with this analysis the time courses predicted by paw movement or neuronal spike rate were now highly correlated with global astrocytic activity, to a level where they explained global astrocytic activity almost equally well as pupil diameter (Fig. 3d). Notably, mean neuronal spike rate was a much better predictor than the average ΔF/F trace before deconvolution, due to the denoising property of deconvolution^27^. Paw movement was a better predictor than mouth movement, and when only licking was used as predictor, predictions degraded even more (Fig. 2d-e). These findings are consistent with direct observations from behavioral monitoring that mouth-only movements did not reliably evoke astrocytic responses (Movie 4). Analysis of additional experiments that used single-plane imaging to record from astrocytes (SO layer) without neuronal imaging confirmed these results (Fig. S4). Based on these findings, we focus in the following on pupil diameter, paw movement, and mean neuronal spike rate as the best predictors of global astrocytic activity. Overall, predictions were not significantly improved when using multiple or higher-dimensional regressors for dilated regression (Fig. S5), suggesting mostly redundant regressors. In agreement with the idea of redundant processes, predictions of global astrocytic activity based on the different regressors were highly correlated (Fig. 3e). For example, predictions based on paw movement and pupil were even more highly correlated among each other than with the global astrocytic activity itself (Fig. 3f, 0.76 ± 0.14 vs. 0.59 ± 0.15, mean ± s.d.; p = 0.00026, Wilcoxon signed-rank test). In addition, two regressors typically performed similarly for the same imaging session but co-varied across sessions (Fig. 3g; Fig. S6). Furthermore, regressors were not correlated but often were able to mutually explain each other when considering multi-timepoint dilated regression (Fig. S7). Together, these analyses highlight that seemingly unrelated behavioral and neuronal variables can explain global astrocytic activity equally well when non-instantaneous dependencies are considered.

### Global astrocytic activity can be explained as a leaky integration of past events

Next, to better understand why multi-timepoint models performed much better than models based on instantaneous correlation, we systematically studied the relative relationship of the observables in time. First, we repeated the regression analysis but used only past or future regressor time points to predict global astrocytic activity. These analyses showed that past but not future time points of paw movement or neuronal spike rate could be used to predict astrocytic calcium transients (Fig. 3h). For pupil diameter, we found a less striking difference between predictors based on past vs. future time points (Fig. 3h) and the multi-timepoint model based on pupil diameter did not exhibit a significant performance increase with respect to instantaneous correlation (cf. Fig. 3c,d; p = 0.53, hierarchical bootstrapping test with 22 sessions from 4 mice). To directly describe the temporal relationship, we attempted to model global astrocytic activity as a temporal integration of mean neuronal spike rate, paw movement, or pupil diameter changes. Instead of using a more complex linear regression with multiple weights for past and future time points as before, we fitted a linear differential equation that simulates global astrocytic activity as a leaky integration of a single variable, with a single free parameter, the time constant τ (Fig. 3i). Interestingly, this simple model showed a trend towards explaining an even higher amount of variance than the cross-validated regression for the paw movement and neuronal activity regressors (Fig. 3j vs. Fig. 3d; p = 0.061 and 0.041 for neuronal spike rate and paw movement, p = 0.75 for pupil; hierarchical bootstrapping test). The integration time constants that were obtained as fit parameters were relatively short for pupil diameter (τ = 2.8 ± 0.5 s; mean ± 90% bootstrapped confidence intervals across sessions) but substantially longer for paw movement (5.6 ± 0.5 s) and neuronal spike rate (6.8 ± 0.5 s). To determine the parameter-sensitivity of the model, we evaluated each model’s performance for a range of time constants. This analysis confirmed the longer time constants for paw movement and neuronal spike rate regressors, and, in addition, highlighted a relatively flat peak of the parameter dependency (Fig. 3k). We conclude that global astrocytic activity can be well described as a nearly instantaneous readout of pupil diameter but, alternatively, also as a leaky integration of past neuronal population spike rate or body movement.

### Global astrocytic activity can be better explained by past body movement than by navigation

While our analyses do not reveal the processes that causally trigger astrocytic activity, they show that mean neuronal spike rate, paw movement, and pupil diameter are almost equally reliable readouts of this process and are therefore largely redundant as explanatory variables for global astrocytic activity. Closer examination of the dataset together with additional experiments enabled us to refine our description. Because of the central role of the hippocampus for navigation, and given the results shown in Fig. 3, it seemed an interesting hypothesis that hippocampal astrocytes would be activated mainly during phases of reward expectation during spatial navigation, as proposed recently^8^. We inspected a set of externally generated and spontaneous behavioral events to investigate this hypothesis. We found that global astrocytic activity indeed ramped towards the location of an expected reward (Fig. S8a). However, this ramping occurred simultaneously with body movements, locomotion, neuronal activity, and pupil diameter changes (Fig. S8a). A similar, albeit smaller, increase of global astrocytic activity could be seen upon spontaneous rewards that were delivered at random time points (Fig. S8b). Given our results of the regression analysis, which showed that spatial location is not a good predictor of global astrocytic activity (Fig. 3d), the more parsimonious explanation of astrocytic activity therefore seems an integration of past events (*e*.*g*., movement or internal arousal), rather than an encoding of future reward. In addition, we found that global astrocytic activity was better described by regressing paw movement than by regressing simple locomotion-associated run speed (Fig. 3c), suggesting that past paw movements rather than locomotion are “encoded” in global astrocytic activity. In line with that, we found that a purely non-navigational behavior, when the mouse occasionally used its hands to reach for the lick spout, most likely to retrieve sugar water residues, consistently elicited an increase of global astrocytic activation despite lack of locomotion (Fig. S9; Movie 5). Moreover, we also observed that behavioral events without significant body movement could elicit astrocytic activation: In a subset of sessions, we applied a salient, rare and unpredictable stimulus (air puff to the left face) when the animal was sitting still. In some cases, this resulted in movement and running of the animal, together with increased pupil diameter and global astrocytic activation (Fig. S8c), consistent with previous cortical calcium imaging experiments of astrocytes during startle responses^16,17,31,32^. However, in a subset of stimulus applications, the mouse remained immobile or frozen despite the stimulus, while the pupil diameter increased together with astrocytic activity (Fig. S8c, top rows of the sorted traces).

Together, our analyses support the idea that hippocampal global astrocytic activity does not primarily reflect behaviors relevant for navigation, such as locomotion and expectation of rewards. While spontaneous past body movement improved our model to explain global astrocytic activity compared to locomotion, it failed to explain global astrocytic activity during immobility. Therefore, our observations support the existing evidence from other brain areas that increased global astrocytic activity is triggered by arousal and mediated by noradrenergic neuromodulation^13^; accordingly, global astrocytic activity is reflected by an increase of the pupil diameter.

### Temporal sequence of events preceding global astrocytic activity

We next aimed to quantify the temporal relationship of the observables functionally associated with astrocytic activity. To assess temporal relationships directly from the experimental data without imposing a model, we computed the correlation functions between observables, which enabled us to estimate the delay of any recorded observable with respect to the global astrocytic calcium signal (Fig. 4a-g). Strikingly, we found consistent delays as quantified by the peak of the correlation function. Whereas the deconvolved neuronal spike rate peaked first (4.2 ± 0.7 s prior to astrocytic calcium signal; Fig. 4a), paw movements followed with a small delay (3.7 ± 0.2 s prior; Fig. 4b; see also Fig. S10), followed by pupil diameter (1.5 ± 0.2 s prior; Fig. 4f). An increase of the positive delay side of the correlation functions would suggest that global astrocytic activity may result in downstream effects on the investigated variables. However, we did not observe such increases at positive lags (Fig. 4a-f) for the observed set of variables. Together, we found, on average, a consistent and stable sequence of events, from neuronal spike rate changes and various body movements, to pupil diameter changes, to astrocytic activation.

**Figure 4.**
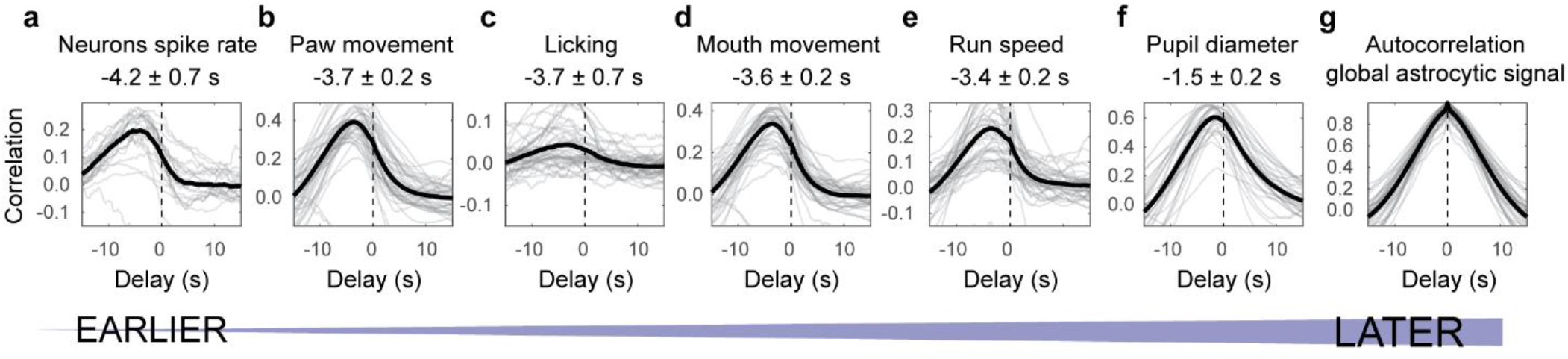
Temporal sequence of neuronal spike rate, motor behaviors, pupil diameter, and global astrocytic activity. Correlation functions were computed between the global astrocytic signal and the variable of interest: neuronal activity (**a**), paw movement (**b**), licking (**c**), mouth movement (**d**), run speed (**e**), pupil diameter (**f**) and astrocytic signal (**g**, autocorrelation). A peak of the correlation function with negative lag indicates that the inspected variable peaked on average earlier than global astrocytic activity. Grey traces are correlation functions extracted from single sessions, black traces are averages across sessions. The delays indicated are median values ± standard error across sessions (n = 22 sessions across 6 animals, except for pupil with 33 sessions across 6 animals and neuronal spike rate with 22 sessions across 4 animals).

### Propagation of astrocytic activity from distal to somatic compartments

Correlation functions average across time points and therefore can reveal temporal relationships that are obscured by variability and noisy single events. To better understand astrocytic activity in single cells, we applied our analysis based on correlation functions not only to the global astrocytic signal but also to the activity of single astrocytic ROIs (either somatic or gliapil regions) by computing the correlation function of each astrocytic ROI with the global astrocytic activation. Surprisingly, we observed that the delay of a given astrocytic ROI with respect to the global mode was variable across ROIs but consistent for a given ROI, and could be relatively long (a few seconds) (Fig. 5a,b). To understand the spatial organization of such delays, we projected the measured delays of ROIs onto the spatial map determined by the mean fluorescence image. We found that ROIs with positive delays (activation after the global mode) tended to map onto regions comprising astrocytic somata, while ROIs with negative delays (activation prior to the global mode) mapped onto regions that did not contain somata or large processes (Fig. 5c,d).

**Figure 5.**
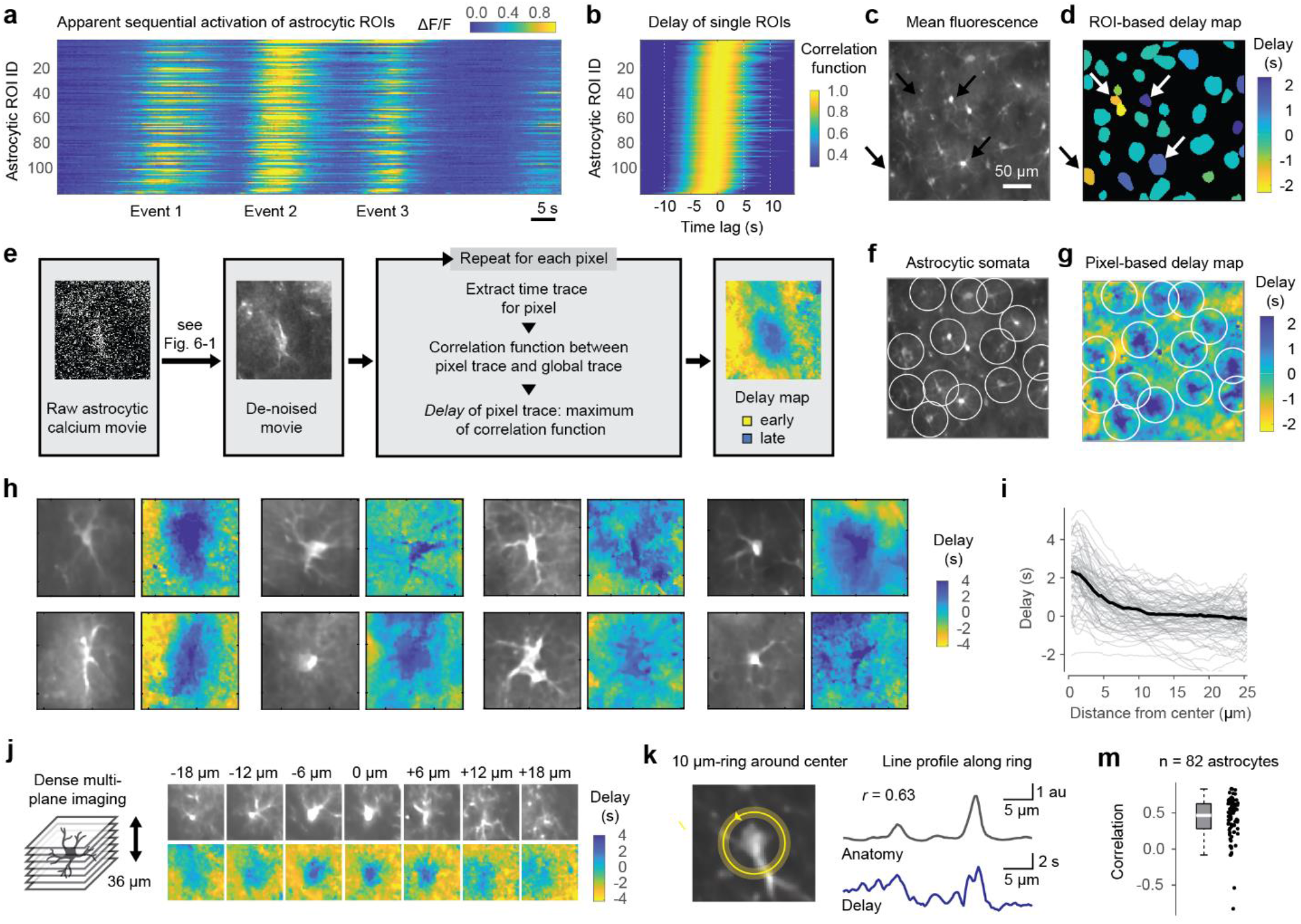
Propagation of astrocytic activity from distal to somatic compartments. **a**, Excerpt of calcium imaging across a population of hippocampal CA1 astrocytes during behavior. Astrocytic ROIs are sorted by the delay of the correlation function with respect to the global mean. An apparent sequential activation of astrocytic ROIs is visible for each of the three events. **b**, Correlation functions of the single astrocytic ROIs, same sorting as in (a). The time window in (a) is only a small part of the activity pattern used to compute the correlation functions. **c**, Excerpt from the mean fluorescence of the astrocytic FOV recording from (a,b). **d**, Delays extracted as maxima in (b), mapped onto the astrocytic ROIs. Right-ward pointing arrows highlight ROIs with strong negative delay (undefined anatomical structure), left-ward pointing arrows highlight example ROIs with strong positive delay (cell bodies). **e**. Processing pipeline for pixel-wise delay maps, consisting of de-noising, pixel-wise computation of the correlation function, extraction of the delay and spatial mapping of each pixel’s delay (see Fig. S11). **f**, Same map as in (c), with white circles highlighting anatomically defined astrocytic cell bodies. **g**, Pixel-based delay map corresponding to the anatomy overview in (f). Somatic regions (center of white circles) are activated with a positive delay (blue), gliapil with a negative delay (yellow) with respect to the global astrocytic activity. **h**, Zoom-in to selected delay maps around anatomically identified cell bodies of astrocytes. The side length of each FOV excerpt is approximately 55 μm. See Fig. S12 for more examples and Fig. S13 for entire delay maps. **i**, Delay vs. distance from astrocytic soma center. Each line is the radial distribution computed from a single astrocyte. The thick black line is the average across astrocytes (82 astrocytes from 11 imaging sessions in four mice). **j**, Example of dense multi-plane calcium imaging, FOV excerpt focused on a single astrocyte. The 3D delay map (computed from denoised data) exhibits the longest positive delay in the imaging plane with the center of the soma. The side length of a single tile is approx. 55 μm. See Fig. S14 for more examples. **k**, A 2.5 μm-thick ring with diameter 10 μm defines a circular line plot that covers both large (bright anatomy) as well as small (dark) processes. Both delay and fluorescence (anatomy) are extract along the circular line plot and are visibly correlated. **m**, Distribution of Pearson correlation values between the fluorescence line profile and the delay map line profile as illustrated in panel (k) across 82 astrocytes (0.46 ± 0.29, median ± s.d.).

Next, to analyze the spatio-temporal astrocytic patterns without the bias of manually selected ROIs, we attempted to use the time trace of each pixel in the calcium movies to determine a fine-grained delay map on a single-pixel level. To enable such a precise spatio-temporal analysis that is normally prevented by shot noise, we trained and used self-supervised deep networks^33^. This de-noising algorithm takes advantage of the temporally and spatially adjacent pixel values to make an improved estimate of the intensity value of the pixel of interest. This method enabled us to de-noise the raw data and generate meaningful time courses for each single pixel (Fig. 5e; Fig. S11a-h; Movie 6). The correlation functions of these single-pixel traces with the global astrocytic signal revealed a smooth map of delays across the FOVs (Fig. 5g), with features that were obscured in the ROI-based delay map (Fig. 5d) and were also less clearly visible in delay maps based on raw data without denoising (Fig. S11i-l). As a striking and surprising feature of these pixel-wise delay maps, delays tended to be negative for gliapil regions devoid of somata and large astrocytic processes, whereas they increased and became positive when approaching the astrocytic somata (Fig. 5f-i; Figs. S12 and S13). This feature of the delay maps, which we refer to as ‘centripetal integration’, indicates that astrocytic activity, on average, propagates from distal, fine processes to the soma, on a timescale of several seconds. To validate this finding, we additionally performed volumetric calcium imaging of multiple closely spaced imaging planes around selected astrocytic somata and computed 3D delay maps. Consistent with our 2D analysis, 3D delay maps consistently exhibited negative delays for fine processes and positive delays in somata in three dimensions (Fig. 5j; Fig. S14).

All these analyses were powered by correlation functions, which averaged across all events of an imaging session and therefore enabled the extraction of spatiotemporal event structures that were mostly invisible to the eye. However, in some cases, we could see calcium events that propagated from distal to somatic compartments also by eye, and we were able to use manually drawn subcellular ROIs or semiautomatically detected event structures (using the AQuA toolbox, ref. ^34^). Using these complementary approaches, we found results that were consistent with our delay maps based on correlation functions (Fig. S15).

To better understand the structural underpinnings of this centripetal propagation, we aimed to analyze whether propagation occurred diffusely, for example through extracellular space, or along visible processes. To this end, we selected the center of clearly identifiable astrocytic somata and then defined a 10 μm-diameter ring around this point. Along this ring, regions that contain large processes can be identified based on their increased fluorescence (Fig. 5k). We found that fluorescence along this ring as an indicator of large processes was positively correlated with the delay for almost all astrocytes (Fig. 5k,m; 82 astrocytes from 11 imaging sessions in 4 mice), suggesting that centripetal propagation indeed proceeds along astrocytic processes. Beyond our primary finding of centripetal propagation, we observed that a few astrocytic somata did not exhibit such a positive delay, suggesting some heterogeneity of the effect (green arrow heads in Fig. S13a,g). Furthermore, visual inspection of raw movies showed substantial local ongoing activity that cannot be described as a propagation from distal to somatic compartments (*e*.*g*., Movies 1, 2 and 6). Therefore, the propagation of activity from distal processes to the central soma dominated the average delay maps but occurred in the presence of other processes that were averaged out in our analysis.

### Centripetal propagation is conditional on the animal’s arousal state

To better understand the features of this centripetal integration in astrocytes towards the soma, we used the delay maps to extract the average time course of all FOV pixels within a specific range of delays (binned in 1-s bins; Fig. 6a,b). Following the results in Fig. 5, the time course of FOV pixels with negative delays thus reflected activity of distal processes, while the time course of FOV pixels with positive delays reflected activity of somatic regions (Fig. 6c). Importantly, this analysis enabled us to compare the time course of the same calcium event in putative distal processes vs. somatic regions. Previously, we had shown that the mean astrocytic activity is delayed compared to neuronal activity by approx. 4 s (Fig. 4a). Now, we found that calcium events in most distal processes occurred ∼2.5 s later than the average firing of simultaneously recorded pyramidal neurons and ∼9 s later in the most central parts (Fig. S16). It seems possible that targeted expression in distal compartments would reveal even shorter delays between neuronal and distal astrocytic activity^10^.

**Figure 6.**
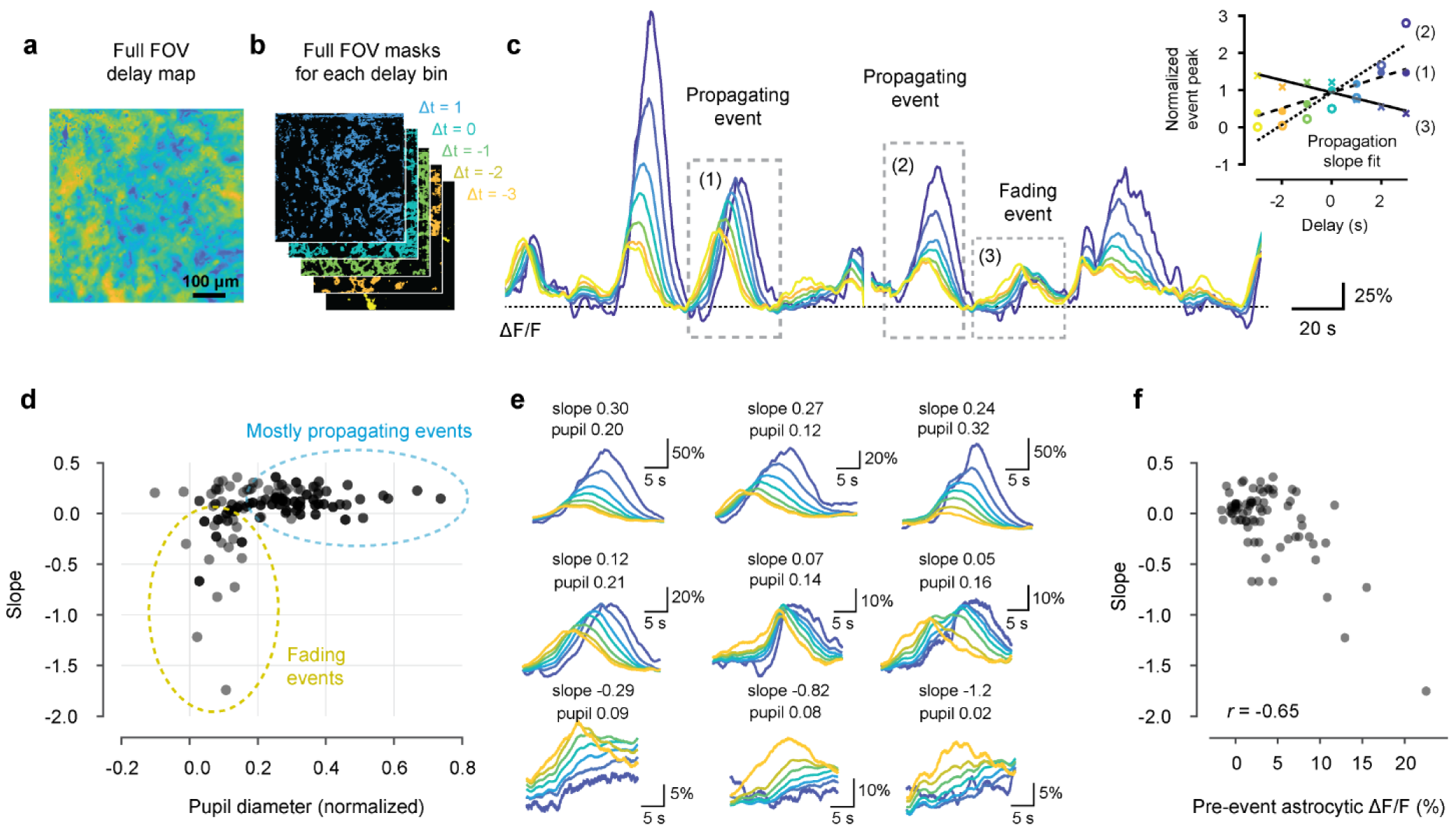
Centripetal propagation of activity in astrocytes is conditional on arousal state and cell-intrinsic calcium signaling history. **a-b**, A smoothed delay map of the entire FOV is binned according to the delays of each pixel (Δt = 1 corresponds to the 0.5…1.5 s interval). **c**, The delay bin masks (b) are used to extract global astrocytic traces from pixels with a specific delay from the de-noised imaging data (color-coding). Due to centripetal propagation, yellow traces represent gliapil and blue traces astrocytic somata. Some calcium events propagate to somata (events (1) and (2)), while others do not (event (3)), quantified by a positive or negative propagation slope, respectively (inset). Therefore, centripetally propagating and non-propagating events can be determined by the slope of peak activity with respect to delay. **d**, Propagation slope of events plotted against the normalized pupil diameter as a proxy for arousal state. Each data point corresponds to a single event. **e**, Example events are plotted in the order of sorted slope values. See Fig. S18 for additional example events. **f**, Pre-event astrocytic ΔF/F, averaged across the 20 s prior to the event, is negatively correlated with the slope value. Only data points with normalized pupil diameter < 0.2 from (d) were included for this panel.

Next, we semi-automatically detected individual events (typically 15-30 s long; see Methods) and found that many detected events reliably propagated activity from distal processes towards the somatic centers of astrocytes. However, some events started in distal processes but then decayed and failed to activate the cell body (Fig. 6c), indicating a thresholding of centripetal propagation. To quantify whether a given event propagates to the center of the cell or not, we computed the “slope” of propagation for each event, determined by a linear fit of the activity peak of the ΔF/F trace vs. the respective delay, such that a negative slope indicates a non-propagating, fading event whereas a neutral or positive slope indicates a centripetally propagating event that activates the soma (Fig. 6c, inset). The majority of events exhibited a slope slightly above or around zero, indicating stable propagation towards the center, whereas some events exhibited a negative slope, reflecting a failure to activate the soma (Fig. 6c-e). These non-propagating events only occurred when the associated pupil diameter (pupil diameter z-scored within each session) was rather low (Fig. 6d) as well as the astrocytic calcium event itself (Fig. S17a). An increased or decreased pupil diameter could reflect either the relatively long-lasting pupil diameter changes during an alert or quiescent state, or phasic pupil diameter changes upon salient sensory or self-generated events. We isolated the phasic component by computing the rectified derivative of pupil diameter as a ‘saliency score’. As for the pupil diameter (Fig. 6d), propagation slopes were negative only for low values of the saliency score (Fig. S17b). Therefore, we find that centripetal propagation in astrocytes failed to occur during low arousal. For very high arousal, on the other hand, we observed events with particularly prominent centripetal propagation. In these cases, centripetal propagation resulted in a non-linear and persistent activation of the soma that outlasted gliapil activation by 10s of seconds (Fig. 6e; Fig. S18e,f). Together, these findings suggest that centripetal integration of astrocytic activity is a non-linear process that is conditional on the animal’s state and facilitated by high levels of arousal.

### Centripetal propagation is conditional on cell-intrinsic calcium signaling history

We noticed that some events with similar normalized pupil diameter or saliency score were propagated to the soma (slope >0) but others not (slope <0; Fig. 6d, Fig. S17b). Previous studies have shown that the strength of astrocytic activation depends on past activity, described as a “refractory period” or cell-intrinsic “memory trace” dependent on intracellular Ca^2+^ store dynamics^21,35,20,36^. Indeed, we found that the slope of events associated with low arousal (normalized pupil diameter <0.2; Fig. 6d) was negatively correlated with the mean astrocytic ΔF/F during the 20 s prior to the investigated event (Fig. 6f). This analysis shows that centripetal propagation is conditional not only on the animal’s state as measured with the pupil diameter, but also on the prior history of astrocytic activation.

### The strength of centripetal propagation can be variable across astrocytes

Next, to investigate the variability of centripetal propagation across cells and to better understand possible cell-intrinsic effects, we extended our analysis to single astrocytic domains. To this end, we used seeded watershed segmentation to approximate distinct astrocytic domains (see Methods) and to quantify centripetal dynamics within each of the individual domains (Fig. S19a-d). The majority of astrocytic domains followed a correlated pattern that consisted of either propagating or non-propagating dynamics for a given event (Fig. S19e). This correlation among putative domains only weakly depended on the inter-domain distance (Fig. S19f). Interestingly, the propagation slope in some astrocytic domains was much higher than for other domains during the same event (Fig. S19d,g), suggesting that the strength of somatic activation upon centripetal propagation is not binary (propagation vs. non-propagating) but follows a continuous distribution across astrocytes for a given global event. As another interesting and related observation, we noticed astrocytes that did not follow the global pattern, *e*.*g*., exhibiting a centripetally propagating event while most other domains exhibited non-propagative behavior (Fig. S19d,h). These results suggest that conditional centripetal propagation of activity, while mostly synchronized across the population of astrocytes, is a process that has the potential to occur separately in each astrocytic domain and could therefore be determined independently for each astrocyte.

### Optogenetic activation of locus coeruleus reproduces centripetal propagation in astrocytes

Given that the observed centripetal propagation was conditional on arousal, we investigated how arousal is communicated to hippocampal astrocytes. One of the key brain regions which communicates arousal throughout the brain and which has been shown to activate astrocytes in the neocortex by noradrenaline release^13,17^ is the locus coeruleus (LC) in the brain stem.

To understand the role of LC, we used an optogenetic activation approach. First, we expressed the optogenetic activator ChR2 specifically in LC neurons using DBH-iCre mice. In the awake mouse, we then stimulated LC with an optical fiber while simultaneously recording virally induced GCaMP6s bulk fluorescence from hippocampal astrocytes in the ipsilateral CA1 using fiber photometry with a second optical fiber (Fig. 7a; Methods). Upon LC stimulation, we observed a striking increase of bulk fluorescence, which was comparable to tail-lift evoked^37^ astrocytic signals (Fig. 7b,c), confirming a role of LC in activating hippocampal astrocytes.

**Figure 7.**
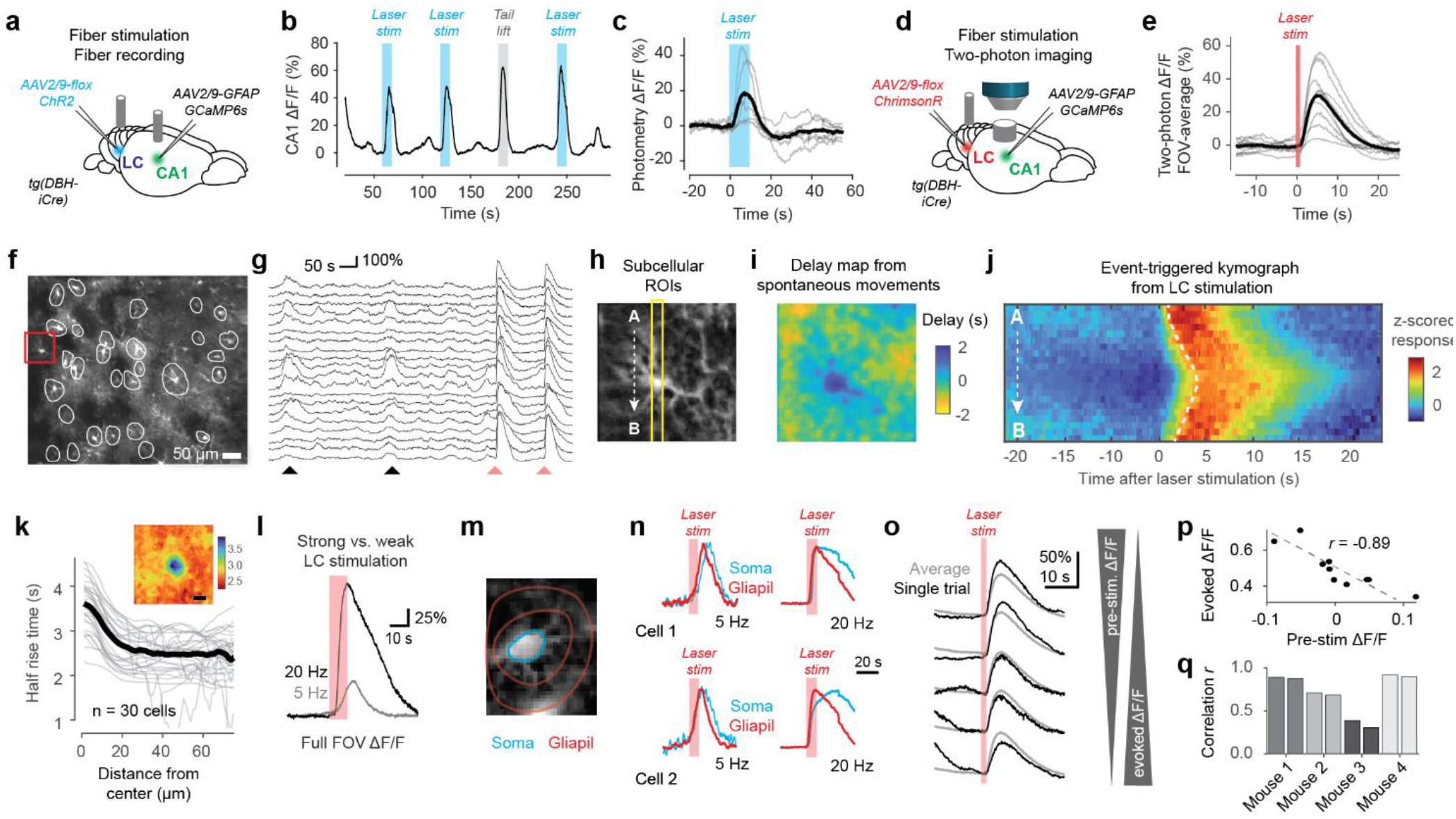
Optogenetic activation of locus coeruleus triggers centripetal propagation of calcium signals in astrocytes. **a**, Setup for fiber-optic recording of calcium signals in hippocampal astrocytes (CA1) together with optogenetic stimulation of locus coeruleus (LC). **b**, Typical example of fiber photometry during optogenetic LC stimulation (blue shading) or during external stimulation as positive control (tail lift; grey shading). **c**, Stimulus-triggered average ∆F/F from fiber photometry in CA1 (grey traces for individual mice) . **d**, Setup for two-photon imaging of calcium signals in hippocampal astrocytes together with optogenetic LC stimulation. **e**, Stimulus-triggered average of FOV-averaged ∆F/F across 4 mice (grey traces for 9 individual imaging sessions). **f**, Example FOV in the *stratum oriens* with manually labeled ROIs. **g**, ∆F/F traces from the selected subset of ROIs surrounding astrocytes from (f) with spontaneous astrocytic population activity related to movement (black arrow heads) and opto-evoked astrocytic activity (blue arrow heads). **h**, Zoom-in to a single astrocyte from the FOV (red rectangle in (f); enhanced contrast). **i**, Delay map for the astrocyte shown in (h), computed as in Fig. 5 using the spontaneous activity and excluding optogenetic stimulation periods from denoised imaging data. **j**, Optogenetic stimulation-triggered average kymograph along the A→B line in panel (h), showing centripetal propagation in a single astrocyte. Average across 10 stimulations across 18 min, computed from raw non-denoised data. Dashed line indicates half-rise time. **k**, Radial plot of the half rise-time of LC-evoked responses as a function of distance from the astrocytic soma center. Inset shows the underlying 2D map of half rise times (scale bar 20 μm). Average across 30 astrocytes with clearly identifiable soma in the imaging plane from 4 imaging sessions in 4 mice. **l**, Example of opto-evoked average responses for weak (5 Hz, grey) and strong (20 Hz, black) stimulation from a single imaging session. **m**, Schematic of ROIs manually defined to separately analyze somatic and gliapil calcium signals. **n**, Examples of persistent activation of soma (blue) but not gliapil (red) for strong but not weaker stimulation in individual astrocytes. See Fig. S20 for more examples. **o**, Strength of optogenetically evoked responses depends on pre-stimulus calcium levels. Example of full-FOV responses for a single imaging session. Responses are decreased for higher pre-stimulus ∆F/F levels. **p**, Pre-stimulus ∆F/F and optogenetically evoked responses are negatively correlated (quantified from the data partially shown in (o)). **q**, The absolute value of the correlation shown in (p) plotted across mice (two imaging sessions per mouse; statistically higher than for shuffled data with p < 0.001 for mice 1, 2 and 4, and p = 0.047 for mouse 3).

Next, we wanted to investigate the cellular and subcellular dynamics of the LC-driven activation of hippocampal astrocytes. LC stimulation could result in simultaneous activation of all subcellular compartments of astrocytes, or could, alternatively, trigger the process of centripetal propagation directly. To dissect these possible effects of LC stimulation, we used a fiber for optogenetic stimulation of LC in awake animals using the red-shifted opsin ChrimsonR (virally induced in DBH-iCre transgenic mice). At the same time, we performed two-photon calcium imaging of hippocampal astrocytes with subcellular resolution (Fig. 7d-f; Methods). As expected from fiber photometry recordings, LC stimulation triggered strong calcium responses when averaged across the field of view (Fig. 7e; Movie 7). Astrocytic calcium signals across the population of astrocytes were increased both during periods of spontaneous movement and arousal and during LC stimulation (Fig. 7g). From periods of spontaneous behavior, we could extract delay maps around single astrocytes to replicate our finding of centripetal propagation (Fig. 7h-i). Interestingly, we observed LC-evoked calcium dynamics that mirrored the spontaneously generated centripetal propagation (Movie 8), as shown by a stimulation-triggered kymograph for an example neuron (Fig. 7j). We quantified the half-rise time of LC-evoked calcium transients around anatomically identifiable astrocytic somata (n = 30 astrocytes across 4 mice) and found increased half-rise times closer to the soma (Fig. 7k). These results show that optogenetic LC stimulation is capable of reproducing not only global astrocytic activation but also centripetal propagation in single astrocytes.

The locus coeruleus exhibits a range of firing patterns depending on external and internal inputs^38,39^. We found that, as to be expected, astrocytic activation was stronger and faster when we increased the stimulation pulse frequency (20 Hz vs. 5 Hz) while leaving the single pulse duration constant (Fig. 7l). However, we also made a surprising observation. For 5 Hz stimulation, somatic activation seemed mostly a delayed version of the calcium signal in the gliapil processes, suggesting an approximately linear propagation of calcium signals. For 20 Hz stimulation, however, gliapil calcium signals decayed after stimulation offset, while somatic calcium signals persisted much longer (Fig. 7m-n, Fig. S20). Interestingly, this non-linear persistent somatic activation induced by artificial LC stimulation mirrors our previous observation of persistent somatic but not distal calcium signals in a subset of centripetal propagation events during very high arousal (Fig. S18e,f).It also reflects our previous observation that some astrocytes exhibited higher propagation slopes than others upon the same behavioral event (Fig. S19). These results suggest that the strength of LC activation determines the strength and duration of somatic calcium signals upon centripetal propagation. It is tempting to speculate that the arrival of the astrocytic calcium signal in the soma might also trigger transcriptional changes, given the recently described transcriptional changes in hippocampal astrocytes upon LC stimulation^40^.

Additionally, we probed our previously established observation that pre-event calcium concentration levels affect centripetal propagation (Fig. 6f). To this end, we analyzed how LC-evoked ∆F/F increases were affected by pre-stimulation ∆F/F levels. We consistently found across 4 mice that higher pre-stimulus levels reduced evoked responses, while lower pre-stimulus levels increased evoked responses (Fig. 7o-q). This finding is consistent with our previous result based on calcium signals during spontaneous behavior (Fig. 6f) and supports the idea that calcium signaling during centripetal propagation depends on internal calcium stores that are partially depleted after prominent calcium events.

### Inactivation of noradrenergic signaling

Finally, we attempted to inactivate astrocytic calcium events and centripetal propagation. To this end, we used several complementary experiments that were based on the hypothesis that centripetal propagation controlled by the LCis mediated by noradrenaline release.

First, we investigated astrocytic and neuronal calcium signals in hippocampus in animals during isoflurane anesthesia, a state where noradrenaline release is reduced^41,42^ (Fig. 8a). We found that astrocytic activity was strikingly reduced. The global astrocytic activity observed during behavior was completely absent whereas some astrocytes still exhibited local events (Fig. 8b; Fig. S21). At the same time, and consistent with recent results in CA1^43^, neuronal spike rates were slightly but significantly reduced (Fig. 8c; 0.14 ± 0.03 Hz vs. 0.09 ± 0.01 Hz; p < 10^-4^, hierarchical bootstrapping test). Moreover, the mean neuronal spike rate failed to explain global astrocytic activity beyond chance level during anesthesia (Fig. 8d; p = 0.88, Wilcoxon signed-rank test). These findings are consistent with previous studies that reported strongly reduced astrocytic calcium levels in cortex during anesthesia^44,11^ or during sleep^45^. Global astrocytic activity re-occurred, however, when the animal woke up and started to move (Fig. S21). These results suggest that awake behavior and the associated neuromodulator release is necessary to trigger global astrocytic activity.

**Figure 8.**
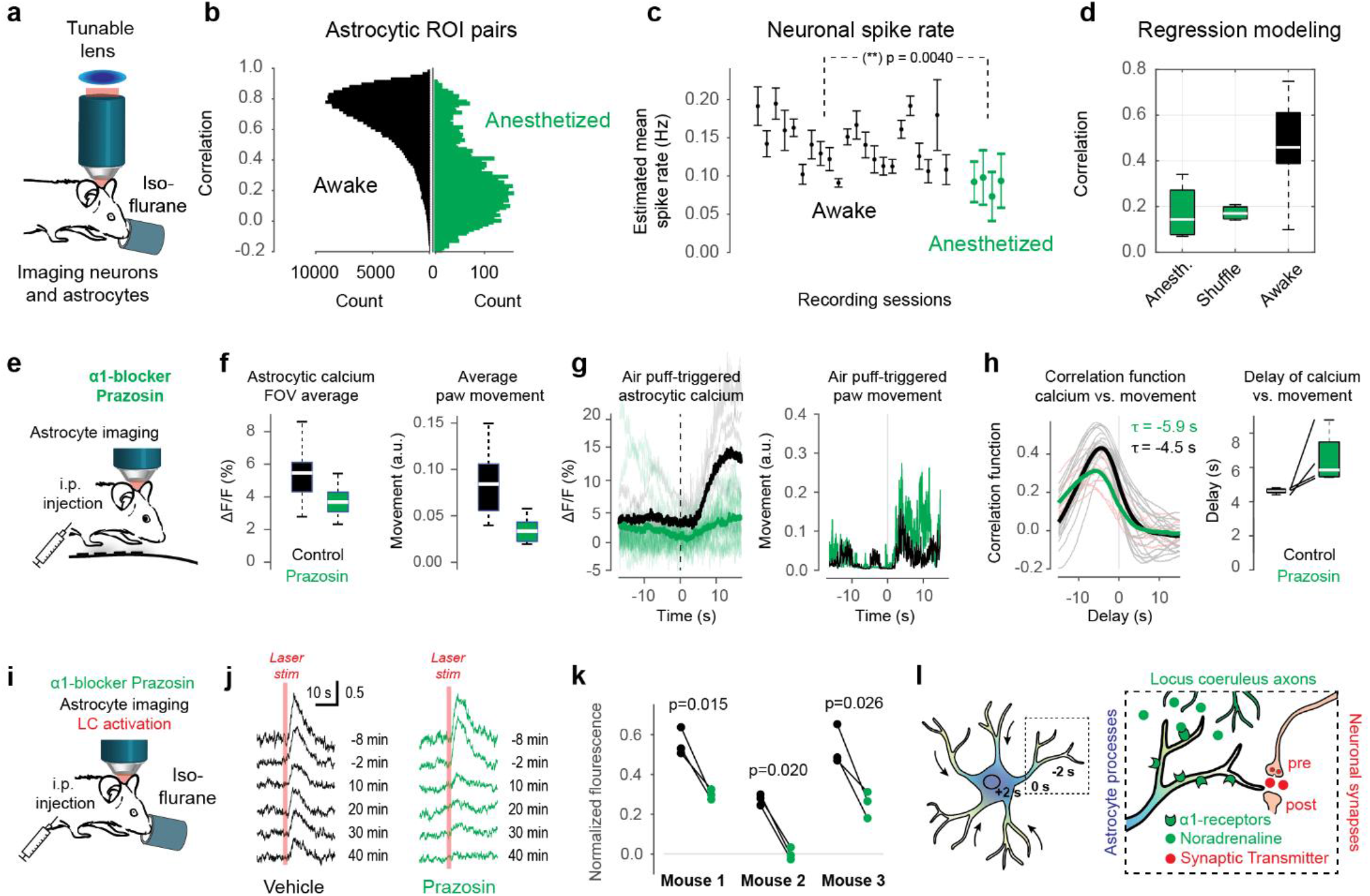
Perturbation of LC-driven noradrenergic signaling impedes global calcium signals and centripetal propagation. **a**, Simultaneous imaging of astrocytes and neurons during isoflurane-induced anesthesia (1.5% in O_2_), a state of reduced noradrenergic signals. **b**, Pairwise correlation between astrocytic ROIs during awake and anesthetized conditions. The awake distribution corresponds to Fig. 1i. **c**, Estimated neuronal spike rate for each awake and anesthetized session (mean ± s.d. across neurons), showing a slight reduction of spike rates during anesthesia (p < 10^-4^, hierarchical bootstrapping test). **d**, Global astrocytic activity is well predicted by neuronal activity during wakefulness but not during anesthesia. **e**, Two-photon imaging of astrocytic calcium signals during behavior after i.p.-injection of the α_1_-blocker prazosin. **f**, Prazosin injection reduced average global astrocytic calcium levels (left; computed with median as F_0_ baseline) but also spontaneous body movement (right). Data from 19 vs. 8 imaging sessions for control vs. prazosin condition in 4 mice. **g**, Reduced stimulus-evoked calcium signals after prazosin injection (left) despite increased stimulus-triggered movement (right) (17 air puffs for prazosin, 6 air puffs for control condition). **h**, Correlation functions showing that astrocytic activity is more delayed with respect to movement after prazosin injection than after control, indicating slowed centripetal integration in the prazosin condition (left). Delay in paired condition (animal on first day with saline injection, second day with prazosin condition) with increased delay seen for each pair (right; n = 4 animals). See Fig. S22 for the per-animal correlation functions. **i**, I.p.-injection of the α_1_-blocker prazosin using a catheter with simultaneous astrocytic two-photon imaging and optogenetic LC stimulation during anesthesia. **j**, Example (mouse 1) for reduction of opto-evoked calcium signals after injection. The reduction is visibly stronger for the prazosin condition (right) than for the saline condition (left). **k**, Replication of the reduction of opto-evoked calcium response after prazosin application (panel j) across three mice. ∆F/F responses are normalized to pre-injection levels (“1.0”). Data point triples are taken from time points 20, 30 and 40 min after injection and statistical tests (t-test) are applied for these pairs of triples. **l**, Schematic working model of conditional centripetal propagation in astrocytes, gated by noradrenergic arousal signals of the LC.

Next, we attempted to directly block noradrenaline signaling pathways using the selective α1-adrenergic receptor antagonist prazosin. After i.p. injection of prazosin, calcium signals in the behaving animal were strongly reduced (Fig. 8e,f). However, prazosin exhibited obvious side-effects on mouse behavior, in particular a reduction of overall movement (Fig. 8f). Since calcium signals in awake mice are highly correlated with behavioral engagement and movement (Fig. 3,4), it remains unclear whether the reduction of calcium signals was due to astrocytic α1-receptor blockade or due to a general reduction in movement. To define a more controlled experiment, we applied air puffs to induce astrocytic calcium signals and found that responses appeared indeed weaker after prazosin-injection compared to control conditions (∆F/F increase during a 15-s window upon stimulation: 3.2 ± 1.9% for prazosin and 9.1 ± 4.6% for control; p = 0.0087, Mann-Whitney test; Fig. 8g). However, the number of recorded air puffs was low and unequally applied across imaging sessions (total of 17 air puffs in 5 imaging sessions from 4 mice for prazosin; 6 air puffs in 2 imaging sessions from 2 animals for control) and therefore precludes any strong conclusion.

Thus, to further assess the contribution of noradrenaline and α1-receptor activation to centripetal propagation, we reasoned that centripetal propagation would be slowed down when the action of noradrenaline was inhibited, similar to our finding that centripetal propagation was slower for weaker LC stimulation (Fig. 7l). Such a slowing down of spontaneous activity would be apparent in the delay of astrocytic signals with respect to a temporal reference signal like movement (cf. Fig. 4). We therefore analyzed the correlation functions of global astrocytic activity and paw movement. We observed that the correlation functions were indeed shifted to longer delays (5.9 s for prazosin vs. 4.5 s for control, average across animals and sessions), and we saw this increase when comparing control and prazosin sessions for each of four animals (Fig. 8h, Fig. S22). These results show that astrocytic calcium dynamics during global events were not completely abolished but reduced and slowed down when α1-receptors were inhibited by prazosin.

Finally, we used our previously established optogenetic LC stimulation paradigm to provide more clearly defined stimulation events. To establish highly controlled conditions, we performed these experiments under anesthesia. Short calcium imaging segments with LC stimulations were temporally spaced before and after an i.p. injection of prazosin. The injection was performed using a catheter that enabled us to leave the mouse in place and continuously image the same FOV. LC-evoked responses slightly decreased during the course of anesthesia also for a vehicle injection (Fig. 8j, left; black dots in Fig. 8k; response amplitude normalized to pre-injection levels). However, injection of prazosin further decreased evoked responses significantly and consistently across mice (Fig. 8j, right; green dots in Fig. 8k; comparison across three stimulus repetitions 20, 30 and 40 minutes after injection with prazosin vs. vehicle; p-values for each mouse, t-test: p = 0.026 p = 0.015 and p = 0.020). Interestingly, as for the above experiments based on prazosin (Fig. 8e-h), astrocytic calcium responses were reduced but not completely abolished.

In conclusion, our inactivation experiments show that awake behavior is necessary for global astrocytic activity and centripetal propagation, likely due to the absence of strong neuromodulatory signals during anesthesia. Furthermore, blocking α1-receptors reduced global astrocytic signals and slowed down the global astrocytic dynamics that govern centripetal propagation.

## Discussion

In this study, we provide evidence that hippocampal astrocytes can be interpreted as slow temporal integrators of salient past events. We demonstrate that this integration proceeds centripetally – from distal processes to the soma. We further show that centripetal propagation is conditional both on the state of the astrocyte (past calcium transients) and the state of the animal (level of arousal), and we found that it can be replicated by stimulation of the locus coeruleus (LC) and impeded by blocking of noradrenergic signaling. Together, these findings suggest that conditional centripetal propagation represents a new principle for spatiotemporal calcium dynamics in astrocytes, establishing astrocytes as computational units of the brain that act on a time scale that is much slower than for neurons.

### The global astrocytic mode dominates in hippocampal CA1 *in vivo*

Recent evidence in cortical and cerebellar brain areas has shown that astrocytic activity *in vivo* exhibits a global mode of activation most likely mediated by noradrenergic neuromodulation^16,17,7,46,20,47^. Here we show that such a global pattern also describes the astrocytic population across hippocampal CA1 layers in awake (Fig. 1) but not anesthetized mice (Fig. 8a-d). In our experiments, calcium signals emerged quasi-simultaneously across the entire field of view (Fig. 5g; Fig. S13), consistent with the idea of a common, global influence converging on astrocytes, and arguing against a dominant role for wave-like propagation of activity within the syncytium of astrocytes through gap junctions (reviewed for example by ref. ^9^). Consistent with this finding, noradrenergic axonal terminals in hippocampal CA1 were recently shown to be highly correlated in their activity, suggesting a global neuromodulatory effect^48^. In addition to the global activity, we also observed prominent local events in subsets of astrocytes or individual astrocytes that were independent of global events (Fig. 1f,g; Fig. S21) and rich ongoing local activity in distal processes that could only be partially resolved using two-photon imaging (Movies 1, 2 and 6), consistent with converging evidence that astrocytic calcium events *in vivo* are prominent in distal processes^9–11,49,50^.

### Astrocytic activity reflects neuromodulation, neuronal activity, and movement

Astrocytic calcium signals in rodents have been shown to exhibit remarkable features that seem to be partially conserved across brain regions. First, astrocytes become globally active as a consequence of salient stimuli, most likely mediated by noradrenergic neuromodulation. While there is to our knowledge no direct evidence that neuromodulation influences specific behaviors in mammals via astrocytic activation, such evidence has been found in zebrafish larvae and drosophila^51,52^. Second, astrocytes in neocortex and cerebellum are activated upon movement or locomotion of the animal^19,21,17,20^. Connecting the aspects of arousal and movement, the temporal relationship of body movements and locomotion have been shown to be closely related to pupil diameter changes^15,53,54^. Pupil diameter, notably, is considered a proxy of noradrenergic (and also cholinergic) modulation, and was found to be delayed by approximately 1 s with respect to the activation of noradrenergic axon terminals in cortex^28^. Third, a causal effect of neuronal activity on astrocytic activity has been proposed, albeit with mixed evidence in favor or against^6,49,46,7^, including the interesting idea that the combined effect of neuronal and noradrenergic activity can result in astrocytic activation^17,55^. Here, we have studied these three processes – local neuronal activity, body movement, and pupil diameter – systematically and simultaneously, together with the activity of hippocampal astrocytes (Fig. 2). We find that all these factors can explain a high degree of variability of the astrocytic signal but only if non-instantaneous metrics are employed, i.e., when taking into account the influence of past events on the present astrocytic signal (Fig. 3). The strong redundancy of these three explanatory variables might be surprising but is possibly related to a common origin of movements, movement-related neuronal activity, and neuromodulatory signals in brainstem circuits^56^. Hence, our systematic investigation reveals candidates for coupling of behavioral and neuronal variables to astrocytic activation that might not be necessarily causal, *i*.*e*., sufficient for activation, but still physiologically relevant. Our quantitative description of the temporal sequence of the observed variables and events (Figs. 3,4; Fig. S8) provides a systematic scaffold to understand and further probe such functional coupling.

Going beyond these observations, our perturbation experiments using optogenetics and pharmacology (Figs. 7,8) highlight that prominent noradrenergic signaling through the LC is sufficient to induce global astrocytic activity. Under physiological conditions, additional factors might contribute to astrocytic activation, including neuronal activity or other neuromodulatory signals, e.g., from cholinergic neurons^57,58^. However, our results strongly support the idea of a central role of noradrenergic afferents from the LC rather than neuronal activity or movement to gate the emergence of global events.

### A caveat for studying the role of astrocytes in behaving animals

Our systematic description also highlights a potential caveat for future functional studies of astrocytic activity *in vivo*. Systems neuroscience has only recently become more broadly aware that neuronal signals across the entire brain might be explained by global or specific motor signals^14,15^. As a reflection of this insight, typical systems neuroscience studies now regularly consider task-irrelevant movement^59^, internal states^56^ or mouse-specific strategies^60,61^ as confounding variables or as features of neuronal representations. We hope that our systematic study, together with other recent astrocyte-focused work^20^, will raise awareness in this regard also in the field of astroglia physiology. The disentanglement of these processes, which has turned out to be challenging for neurons^56^, is not easier for astrocytes. For example, we show that paw movement is a better predictor of astrocytic activity compared to simple locomotion or running speed; therefore, video monitoring of behaviors should be considered more reliable for the observation of confounding motor movements than the measurement of pure displacement (Fig. 3c). In addition, we find that instantaneous measures of coupling fail to capture the delayed relationships of astrocytic signals with movement and neuronal activity (Fig. 3c; Fig. S4).

As another caveat from a more technical perspective, we find it important to point out that imaging conditions and labeling strategies might have strong and undesired effects on results obtained with astrocytic calcium imaging. For example, astrocytic somata that are slightly out of focus may be more strongly contaminated by the surrounding gliapil, resulting in temporally shifted calcium transients (Fig. 5a-d). Similar concerns apply to imaging conditions where resolution is degraded due to imaging depth or imaging modality and therefore results in mixed somatic and gliapil signals.

In the light of these findings and analyses, previous studies claiming specific computational roles of hippocampal astrocytes, *e*.*g*., for reward or place encoding^5,8^, would probably benefit from careful controls for confounding variables, in particular those that are shifted in time by several seconds. For example, the activity of astrocytes in a stereotypical behavioral running task could in theory be described to code for the current location but, given our systematic description (e.g., Fig. 4), is much more likely to code for a past location instead. In addition, our study provides strong evidence that astrocytic activity in hippocampus is more parsimoniously explained as a response to past arousing events rather than reflecting an expectation of future spatial location as recently suggested^8^.

### Unbiased analysis using global astrocytic activity as a reference

The complex and often seemingly independent activity patterns in the fine processes of individual astrocytes make it challenging to use ROI-based methods for the analysis of astrocytic calcium signals. Recent years have seen a surge of excellent tools specifically designed for the analysis of astrocytic calcium imaging. However, these tools are based either on discrete spatial ROIs^32,62,63^ or on discrete events^34,64^, in both cases requiring definition of an event via an arbitrary criterion. Here, we computed pixel-wise delay maps with respect to a global reference (the global astrocytic activity), a method without such requirement. Our delay map analysis takes advantage of self-consistent denoising methods^33^, uses the averaging power of correlation functions and projects the extracted properties of the correlation function onto the anatomical map, pixel by pixel (Fig. 5). Moreover, the resulting map can be used for the extraction of fluorescence traces based on the extracted properties assigned to each pixel (Fig. 6). This workflow was the enabling factor for our analysis of centripetal propagation, and it should be generally applicable for the unbiased extraction of average spatio-temporal activity patterns from raw or denoised calcium movies. We provide a well-documented demo code on Github in both MATLAB and Python to facilitate adoption of our method by other researchers (https://github.com/HelmchenLabSoftware/Centripetal_propagation_astrocytes).

### Conditional centripetal integration in astrocytes

The most striking feature of astrocytes is their star-shaped morphology. Here we provide evidence that the soma as the center of this star-shape might act as an integration hub that is activated upon salient events. First, we show that astrocytic activation can be well described by a simple leaky integration differential equation with the integration time constant as a single parameter (Fig. 3k-j). We systematically measure the delay of astrocytic activity with respect to other observed variables and show that neuronal activity, movement, and pupil changes precede astrocytic activity in a sequence with consistent delays (Fig. 4). These results are consistent with previous findings that showed delayed activation of astrocytic somata with respect to external stimuli or self-generated movement^21,20,7,55,16,17^. Second, we show that the global astrocytic activity, which reflects this temporal integration, can be described not only as a temporal but also as a spatial pattern that consists of activity propagating from distal astrocytic processes to the respective soma on a timescale of several seconds (Fig. 5). Consistent with the idea of earlier distal activations, a few recent studies have found that low-latency activation of astrocytes can occur in fine distal processes^11,10,65^. Interestingly, it has been shown in hippocampal slices that calcium events are often restricted to fine processes but expand for stronger stimuli^66^, and sometimes invade the somatic region^11^. It has been hypothesized that larger events result from the integration of smaller local calcium signals and that such integration is thresholded by the stimulus intensity^67^. In our experiments, we systematically collected evidence in behaving animals that calcium events in distal astrocytic processes can either propagate or not propagate to the more proximal and somatic compartments. Moreover, we found that somatic calcium signals were enhanced for higher arousal or stronger LC stimulation (Fig. S18, Fig. 7n). Interestingly, very recent results have replicated our finding of centripetal propagation in cortical brain regions and make it likely that this phenomenon is not restricted to hippocampus^68^.

Our analyses and optogenetic experiments indicate that centripetal propagation is conditional on at least two factors: first, it is facilitated by high LC activity (Fig. 7) that reflects higher levels of arousal induced by recent salient self-generated or externally driven events (Fig. 6); in addition, it is impeded for already elevated calcium concentrations in the respective astrocyte (Fig. 6f; Fig. 7o-q). Previous *in vivo* experiments showed that calcium events in late-but not fast-responding astrocytic regions depend on both IP_3_ ^10,32^ and noradrenergic signaling^10^. Together with our results, this puts forward the hypothesis that calcium signals in distal astrocytic processes are driven by local synaptic activity (largely independent of IP_3_ and noradrenaline) and then propagate to the soma in an IP_3_-dependent manner only when saliency is communicated through noradrenergic signals. Our results therefore suggest that astrocytes slowly propagate calcium signals from their processes into their soma in a conditional manner. The astrocytic soma therefore operates as a computational unit and acts as an integrator of past saliency.

### A role for astrocytes to slowly process past events

Typical neurons not only integrate information from their dendritic processes in the soma but also convey the output of this process, the action potential, via their axon to connected neurons. Our finding of central integration of information in astrocytes therefore raises the question about the potential output generated upon centripetal activation of the astrocytic soma, especially in view of the absence of an ‘astrocytic axon’. Close inspection of the properties of centripetal integration might help to identify the role of such an output. As a first property, global astrocytic activity can be described as a temporal integration of past salient events (events such as a startle response or self-generated movement) (Fig. 3,4). Second, centripetal propagation of astrocytic activity is closely associated to arousal, since it is correlated with pupil dilation (Fig. 3), it can be triggered by LC stimulation (Fig. 7), and it is impeded by noradrenergic blockers (Fig. 8). Third, centripetal propagation acts on a time scale of seconds rather than milliseconds (Fig. 4,5), arguing against a specific role in low-latency neuronal computations and for a role in processes that act on longer time scales. Modulation of neuronal plasticity is a plausible candidate for such a process, since it acts on behavioral timescales^69^ and takes place upon salient events^70^. Such modulation has been studied for hippocampal and other astrocytes both *in vitro*^57,71,72^ and more recently also *in vivo*^58,73–76^.

Our results highlight that centripetal propagation is a non-linear process that can occur in individual astrocytes but not in others (Fig. S19), sometimes also with long-lasting somatic activation in highly arousing situations (Fig. 7m-n, Fig. S20). It is not clear what output signals may be triggered by somatic activation of astrocytes, potentially instantiating feedback modulation of the surrounding neuronal processes. A possible astrocytic output signal that could fulfil this role and warrants further investigation in the context of centripetal integration is gliotransmission, the astrocytic release of glutamate^77–80^, D-serine^57,71,81^ or lactate^82,83^, in particular since this process appears prominently involved in neuronal plasticity^67,84^. It remains, however, to be discovered how somatic astrocytic activity could translate into potentiation or depression of specific synapses. In a possible scenario, synaptic activity preceding centripetal propagation could tag specific synapses by a signaling molecule with a half-life time of several seconds. The potential effect of somatically activated astrocytes would arrive back at neuronal synapses only several seconds later and specifically affect these tagged synapses, for example to induce synaptic depression. Further work and methods development will be required to dissect the mechanisms of such potential astrocytic outputs. However, centripetal integration of past events in astrocytes defines a plausible candidate mechanism how such an output could be orchestrated to conditionally mediate or modulate neuronal plasticity on a behavioral time scale.

## Acknowledgements

This work was supported by grants from the Swiss National Science Foundation (project grant 310030B_170269 and Sinergia grant CRSII5_180316 to F.H.; Ambizione grant PZ00P3_209114 to P.R), the European Research Council (ERC Advanced Grant BRAINCOMPATH, project 670757 to F.H.), the NIH Brain Initiative (grant U01NS115585 to F.H.) and by grants from the University of Zurich (Forschungskredit grants K-41220-04 to C.M.L. and K-41220-06-01 to P.R.; “Filling the Gap” grant to D.B.). The lab of J.B. was supported by the ETH Project Grant ETH-20 19-1, SNSF Grants 310030_172889 and 310030_204372, the Botnar Research Centre for Child Health, the Swiss 3R Competence Center, Roche, and the Hochschulmedizin Zürich Flagship project STRESS. We thank the group of Anna-Sophia Wahl for sharing their experience with the behavioral setup, and Stefan Giger, Martin Wieckhorst and Hansjörg Kasper for help and assistance with its construction. We thank Antoine Adamantidis and Bruno Weber for feedback on the manuscript, and all members of the Helmchen Lab for critical input.

## Contributions

P.R. conceived the study, established the methodology, contributed to all experiments except for fiber photometry, performed all analyses, and wrote the paper. S.N.D. performed fiber photometry experiments and contributed to optogenetics experiments. D.B. contributed to two-photon and pharmacological perturbation experiments. C.M.L. contributed to two-photon imaging experiments and to the conceptualization of results. J.B. supervised the optogenetics and fiber photometry experiments. F.H. conceived the study, supervised all experiments, contributed to analyses, and wrote the paper.

## Data availability

Example raw data of astrocytic calcium imaging together with MATLAB and Python programs to compute delay maps as shown in Fig. 5 have been made available on Github (https://github.com/HelmchenLabSoftware/Centripetal_propagation_astrocytes). All other processed data used in the manuscript will be available from the corresponding author upon request.

## Code availability

Demo programs to compute delay maps from raw astrocytic calcium imaging data as shown in Fig. 5 are provided via Github together with example data under https://github.com/HelmchenLabSoftware/Centripetal_propagation_astrocytes. Example code is provided in both MATLAB and Python. All other custom code used for analyses described in the manuscript will be available from the corresponding author upon request.

## Competing interests

The authors declare no competing interests.

## Methods

### Animals and surgery

All experimental procedures were carried out in accordance with the guidelines of the Federal Veterinary Office of Switzerland and were approved by the Cantonal Veterinary Office in Zurich. We used adult male and female 4-6 month old C57BL/6-Thy1-GCaMP6f (GP5.17, ref. ^26^), which express the calcium indicator GCaMP6f in a subset of pyramidal neuron in hippocampal CA1, C57BL/6-Tg(Dbh-iCre)1Gsc (DBH-iCre, ref. ^85^) mice, which express Cre in noradrenergic neurons, and wild-type C57BL/6 mice. Mice were provided with analgesia (Metacam 5 mg/kg bodyweight and Buprenorphine 0.1 mg/kg, s.c.) prior to surgery. Anesthesia was induced using isoflurane (5% in O_2_ for induction, 1-2% for maintenance during surgery), and the body temperature was maintained at 35-37°C using a heating pad. For surgeries, shaving cream was applied to the dorsal head above the brain, and an incision was made into the skin after local application of lidocaine. To induce expression of GCaMP6s in CA1 hippocampal astrocytes, an injection of AAV virus based on the human GFAP promoter fragment gfaABC1D (ca. 200 nL of ssAAV9/2-hGFAP-hHBbI/E-GCaMP6s-bGHp(A), titer 1.0x10^13^ vg/mL; Viral Vector Facility, University of Zurich) was made in hippocampal CA1 (coordinates: AP -2.0 mm, ML -1.5 mm from Bregma, DV -1.3 from the surface of the dura). The injection pipette was left in place after injection for at least 5 min to prevent reflux. A suture was made to close the skin above the skull, and reopened after two weeks to implant the hippocampal window as described previously by others^86^ and ourselves^87,88^. Briefly, to expose the brain, a 3-mm diameter ring was drilled into the skull, centered at the previous injection site but avoiding interference with the midline head bone structures. To provide a basis for attachment, two thin layers of light-curing adhesive (iBond Total Etch, Kulzer) were applied to the skull, followed by a ring of dental cement (Charisma, Kulzer) to prevent overgrowth of the preparation with skin. A 3-mm diameter biopsy punch (BP-30F, KAI) was inserted into the exposed brain until it reached the corpus callosum and left in place for at least 5 min to stop bleeding. Damage to deeper structures was avoided by a rubber stop added to the punch. Then, a flatly cut off injection cannula (Sterican 27G, B. Braun) connected to a vacuum pump was used to carefully remove the cortex in the exposed region until the white-red stripes of the corpus callosum became visible. The corpus callosum, different from previous studies targeting deeper regions^87^, was left intact. Bleedings, if they occurred, were stopped with absorbant swabs (Sugi, Kettenbach) and hemostatic sponges (Spongostan, Ethicon) before further surgery steps were performed. Then, a cylindrical metal cannula (diameter 3 mm, height 1.2-1.3 mm) attached with dental cement to a 0.17-mm thick coverslip (diameter 3 mm) was carefully inserted into the cavity and fine-positioned with a small-diameter glass capillary attached to the stereotaxic frame. When no further bleeding occurred, the hippocampal window was fixed in place using UV-curable dental cement (Tetric EvoFlow, Ivoclar) that was inserted into the gaps between the skull and the cannula. Finally, tissue glue (Vetbond, 3M) was used to connect the animal’s skin with the ring of dental cement (Charisma). A double wing-shaped head bar was attached to the Charisma ring using dental cement (Tetric EvoFlow) directly after the surgery or in a separate surgery session. After surgery, animals were monitored for 3 days with application of antibiotics (2.5% Baytril in drinking water, Vetpharm), and analgesics (Metacam, 5 mg/kg, s.c.) administered when necessary. Behavioral training started two weeks after surgery. Calcium imaging was performed 2-3 weeks after the start of behavioral training.

For optogenetic experiments in DBH-iCre mice, we followed procedures as previously described^39^. Briefly, a small hole was drilled at AP -5.4 mm and ML -0.9 mm relative to bregma. Mice were then injected unilaterally (Coordinates: AP -5.4 mm, ML - 0.9 mm, DV -3.8 mm) with 1 μL of an AAV construct carrying the optogenetic actuator ChR2 or ChrimsonR (ssAAV-5/2-hEF1α-dloxhChR2(H134R)_ EYFP(rev)-dlox-WPRE-hGHp(A) or ssAAV-5/2-hEF1α/hTLV1-dloxChrimsonR_tdTomato(rev)-dlox-WPRE-bGHp(A); Viral Vector Facility, University of Zurich) using a pneumatic injector (Narishige, IM-11-2) and calibrated microcapillaries (Sigma-Aldrich, P0549). During the same surgery, 400 nL of astrocyte-specific GCaMP6s-inducing AAV was injected to hippocampal CA1 as described above. For fiber photometry experiments (Fig. 7a-c), optical fibers were implanted 200 μm above the injection coordinates of locus coeruleus and hippocampus (diameter 200 μm, NA=0.37; Neurophotometrics, USA). For LC stimulation combined with two-photon imaging (Fig. 7d-q), an angled optical fiber was implanted 200 μm above the injection coordinates (low profile, angle 90°, diameter 200 μm, NA=0.66; Doric Lenses, Canada). Optical fibres were glued to the skull using a bonding agent (Etch glue, Heraeus Kulzer GmbH) and a UV-curable dental composite (Permaplast, LH Flow; M+W Dental, Germany). After 3 weeks, pupillometry was performed as described before to validate functional expression of the actuator^30^. For animals that showed pupil responses upon LC stimulation, hippocampal windows were implanted as described above. The head bar metal was cut at the posterior side of the head to spare an opening for the angled stimulation fiber implant.

### Two-photon microscopy

A custom-built two-photon microscope was used to monitor calcium signals in astrocytes and neurons in either a single or multiple layers of CA1, while the animal was spontaneously running on a linear treadmill. A femtosecond-pulsed laser (MaiTai, Spectra physics; center wavelength tuned to 911 nm; power below the objective 20-40 mW) was sent through a scan engine consisting of a 8-kHz resonant scanner (Cambridge Technology), a 2x magnifying relay lens system, and a slow galvo scanner (6215H, Cambridge Technology), to a dedicated scan lens (S4LFT0089/98, Sill Optics) and a tube lens (Ploessl lens system consisting of two 400-mm focal length achromatic doublet lenses; AC508-400-AB, Thorlabs) before entering the objective’s back aperture. Either a 16x (CFI75 LWD 16X W, NA 0.8, WD 3.0 mm; Nikon) or a 40x objective (CFI Apo NIR 40X W, NA 0.8, WD 3.5 mm; Nikon) were used for calcium imaging. The 16x objective provided a larger field of view (600 μm side length) but the back aperture was slightly underfilled, resulting in an axial resolution of 4-5 μm (FWHM). The 40x objective allowed us to overfill the back aperture, resulting in an improved axial resolution of 2-3 μm (FWHM) at the cost of a reduced FOV (200 μm side length). The 40x objective configuration also enabled the use of a small-aperture tunable lens (EL-10-30-C, Optotune) together with an offset lens (f = -100 mm) just before the back focal plane to enable fast z-scanning over a z-range of up to 300 μm as described previously^24^. For experiments with simultaneous photostimulation of locus coeruleus and two-photon imaging of hippocampal astrocytes, a 10x long working distance water immersion objective was used (Olympus XLPLN10XSVMP, NA 0.6, WD 8 mm) to avoid spatial constraints with the stimulation fiber below the objective, resulting in a larger FOV (600-1000 μm). Single-plane imaging was performed at a rate of 30.88 Hz (512 x 622 pixels). Volumetric rates were reduced accordingly for dual-plane (15.44 Hz), triple-plane imaging (10.29 Hz) and imaging across 7 planes (4.41 Hz). Scanning and data acquisition was controlled with custom-written software programmed in C++ (http://rkscope.sourceforge.net/, ref. ^89^).

### Fiber photometry

GCaMP6s signals were recorded using a commercially available photometry system (Neurophotometrics, Model FP3002) controlled via the open-source software Bonsai (2.6.2 version). The implanted fiber was attached to a pre-bleached recording patch cord (diameter 200 μm, NA=0.39; Doric Lenses). Two LEDs were used to deliver interleaved excitation light: a 470 nm LED for recording GCaMP-dependent fluorescence signal (F^470^) and a 415 nm LED for GCaMP-independent control fluorescence signals (F^415^). The recording rate was set to 120 Hz for both LEDs allowing 60 Hz for each channel. Excitation power at the fiber tip was set to 25-35 μW. Analysis of raw photometry data was performed using a custom-written MATLAB script as described previously^39^. First, to filter high frequency noise (above 1 Hz), a lowpass filter function was applied to both recorded signals (F^470^ and F^415^). Next, to correct for photobleaching, the baseline fluorescence was calculated as a linear fit of the filtered F^415^ signal to the level of the F^470^ signal during the 5-s baseline window preceding each LC stimulation. This rescaled F^415^ signal is termed F^415^_baseline-fit_. Finally, GCaMP signals were expressed as the ΔF/F value:

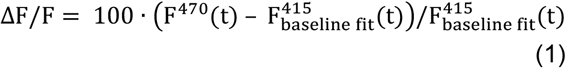

where F^470^(t) signifies the filtered fluorescence value at each time point *t* across the recording and F^415^_baseline-fit_(t) denotes the value of the fitted 415 nm signal at the time point *t*. The final ΔF/F signal was smoothed with a 100-points moving mean filter.

### Optogenetic stimulation

Optogenetic stimulation of locus coeruleus was provided via ChR2 (fiber photometry, Fig. 7a-c) or ChrimsonR (two-photon imaging, Fig. 7d-q) using a 473-nm or 635-nm laser with a fiber output power of approx. 10 mW when illuminating continuously. Stimulation protocols followed previous work to provide stimulation similar to natural locus coeruleus activity during arousal^39^. We used distinct stimulation protocols to address different questions. Long stimulations (10-s duration) were used to provide consistency with previous work^39^. Short stimulations (≤1 s) were used to provide clearly defined phasic stimuli, which more easily enabled stimulus-triggered analysis of centripetal propagation. For all stimulation protocols, 10-ms pulses of light were applied, but at different repetition frequencies. For experiments described in Fig. 7f-m, Fig. 7q-s and Fig. 8i-k, we used a 1-s stimulation with a pulse repetition frequency of 15 Hz (average power 1.5 mW). For experiments described in Fig. 7a-c and Fig. 7n-p, we used a 10-s stimulation with a pulse repetition frequency of 5 Hz or 20 Hz (average power, 0.5 and 2.0 mW).

### Behavioral setup

The treadmill consisted of two custom-designed light-weight wheels, one of which was attached to a rotary encoder (4-mm shaft optical rotary encoder, Phidgets, USA) to measure locomotion of the animal. A 130 cm long and 5 cm wide velvet belt (88015K1, McMaster-Carr) was equipped with sensory landmarks consisting of self-sticking elements, velcro strips and hot glue. A metal tape attached to a single location of the back side of the belt was used as a reflector for an IR sensor to provide a spatial reference signal in order to track the location of the animal on the belt. IR light (LIU850A, Thorlabs) together with a camera (DMK23UP1300, The Imaging Source, recording at 30 Hz; 16-mm EFL objective MVL16M23, Thorlabs) was used to monitor the animal’s behavior during the experiment. In a subset of experiment sessions (33 out of 42), a small and partially shielded UV LED (LED370E, Thorlabs) was directed towards the right eye of the animal, resulting in a less dilated pupil and allowing for pupil segmentation from the behavioral video. Rewards in form of sweetened water (30% sugar) were provided through a metal lick spout to the animal at a specific location of the belt that remained unchanged across sessions. Reward delivery was controlled by a solenoid valve (VDW22JA, SMC) that was gated by a relay circuit (Sertronics Relais Module, Digitech). Mice were free to consume the reward whenever they wanted, immediately after reward delivery or later. If mice did not retrieve a reward for 50 s through running, a spontaneous reward was delivered. Reward delivery time points were automatically recorded in the software, and reward consumption (first contact of the mouse’s tongue with the water drop) was manually detected for each reward from video monitoring. The time of reward consumption was used for the analyses in Fig. 4e,f. In a subset of experiments, brief air puffs to the left side of the animal’s face as additional sensory cues were provided randomly and rarely (maximally once per minute) without predictive cue. To enable the disentanglement of arousal generated by air puffs and movement, air puffs were only applied when the animal was not running for ≥10 s.

The behavioral setup was controlled using custom-written Python code, which controlled valves, camera triggers, and microscope acquisition start, and recorded the position of the rotary encoder, the IR sensor for absolute position and behavioral events like air puffs or water rewards. The temporal offset and relative recording speed of the camera with respect to two-photon scanning were calibrated in a separate experiment where both the behavioral camera and the two-photon software recorded the same signal of a flashing light.

### Behavior training and imaging experiments

One week before experiments, drinking water of mice was supplemented with citric acid (2% of volume) to motivate the mice during the running task^90^. Mice were handled by the experimenter for 15-20 min per day for 3 days without exposure to the behavioral setup. Afterwards, they were accustomed to the behavioral setup and the lick spout. For the next 3-7 workdays, depending on the animal’s ability to participate in the task without being stressed, mice were trained to sit or run on the treadmill while being head-fixed. When animals readily ran on the treadmill and consumed sugar water rewards for 15-20 min, imaging experiments were performed.

A behavioral session lasted in total for 15-35 min and consisted of 140 s-long recording segments, spaced by short breaks of 5-20 seconds due to limitations of the two-photon microscope software. The imaging plane was chosen to lay in the central part of *stratum oriens* of CA1 and to clearly contain visible astrocytic somata if possible. For dual-plane imaging of neurons and astrocytes, the first imaging plane was centered at the pyramidal cell layer of CA1, and the second imaging plane 60-90 μm more dorsally. For triple-layer imaging of astrocytes, imaging planes were spaced approximately 70-80 μm from each other, with the central plane residing in the pyramidal cell layer. For triple-plane imaging during anesthesia, two imaging planes spaced by 10-15 μm were positioned in the pyramidal layer of CA1 such that not the same neurons were sampled in both planes and the third plane 60-90 μm more dorsally. For volumetric imaging of astrocytes, the seven imaging planes around an astrocyte were spaced by 6 μm between adjacent imaging planes. Different sessions were performed on different days, and the FOV was changed between days.

### Pharmacology

Prazosin (Tocris, Cat No. 0623) was dissolved in distilled water (10 mg in 100 mL millipore) and mixed. For experiments described in Fig. 8e-h, mice were lightly anesthetized in isoflurane, i.p.-injected with 10 μL/g (solution/bodyweight) and then were allowed to wake up and recover from anesthesia without delay. Anesthesia during injection was applied to facilitate procedures and to reduce stress by injection and handling of the animals. Experiments started 20-25 min after injection and lasted 15-25 min. For experiments described in Fig. 8i-k, animals were anesthetized (5% isofluorane) and then quickly transferred to the head-fixation setup, where anesthesia was continued (2% and 1.5% isoflurane for male and female mice, respectively). A catheter consisting of the tip of an insuline needle (Braun, Omnican 50 LDS) was inserted to provide i.p. injection via an injection syringe. After baseline recording, i.p.-injection was performed, and optogenetic stimuli were applied together with calcium imaging every 10 min after the timepoint of injection. For control, the same procedure was repeated with distilled water instead of prazosin solution.

### Histology

After completion of calcium imaging experiments (approx. 2 months after start of behavioral experiments), animals were administered a lethal dose of pentobarbital (Ekonarcon, Streuli) and transcardially perfused with 0.1 M phosphate standard solution (PO4) followed by 4% paraformaldehyde (PFA, in 0.1 PO4). For histology experiments to check co-localization of GCaMP and GFAP (Fig. S1e-g), 60-μm thick coronal brain sections were stained with DAPI and anti-GFAP (primary AB rabbit-anti-GFAP 1:1000, secondary AB anti-rabbit with Cy3 1:250) and imaged with a confocal laser-scanning microscope (Olympus FV1000). Three separate channels recorded the nuclear stain DAPI, the intrinsic GCaMP6s of virally induced GFAP-based expression, and the astrocytic antibody staining of astrocytes with Cy3. For experiments to compare GFAP levels in injected vs. non-injected hemispheres of the same animal (Fig. S1a-d), the same histology procedure was applied. Care was taken to apply the same imaging power, zoom settings and imaging depth below the surface for ipsilateral and contralateral imaging sites. For the quantification of mean fluorescence and area fill fraction, we followed existing standard protocols, with the same threshold used to compute the fill fraction for both hemispheres for each slice^91^.

### Calcium imaging post-processing

Raw calcium imaging movies were spatially resampled to remove the distortion induced by non-linear sinusoidal scanning in the x-axis. Then, rigid movement correction in the xy-plane was applied^92^. Analyses in Fig. 1 were based on manually defined ROIs; analyses in Figs. 3, 4 and 5 were based on the global astrocytic activity, which we define as the average ΔF/F signal across the population of astrocytes within the FOV; and analyses in Figs. 6 and 7 were based on pixel-based activity traces (described below). Active ROIs were extracted manually with a previously described toolbox (https://github.com/PTRRupprecht/Drawing-ROIs-without-GUI, ref. ^93^), based on anatomical features of the mean fluorescence, as well as on the map of local correlations which highlights correlated spatially close pixels^94^.

For neuronal imaging data, both active and inactive neuronal somata were included to provide an approximately unbiased estimate of neuronal spike rates. For experiments with expression in both neurons (GCaMP6f) and astrocytes (GCaMP6s), bleedthrough of the gliapil signal in the pyramidal layer resulted in a slow contaminating signal superimposed onto the neuronal signals. We used the mean fluorescence in a 15-pixel wide surround region of the respective neuronal ROI to linearly unmix the astrocytic contamination (Fig. S3). Pixels that were part of other neuronal ROIs were not included in the surround ROI. Next, deconvolution of the neuronal traces with a deep network trained on a large ground truth dataset (CASCADE; ref. ^27^) was performed in Python, suppressing shot noise unrelated to spike-evoked calcium signals as described before^27^, but also discarding non-neuronal signal components like residual slow components stemming from astrocytes (Fig. S3).

For astrocytic imaging data, all clearly visible astrocytic cell bodies were selected as ROIs, but also active gliapil regions or putative astrocyte endfeet that showed a spatially coherent response as seen by the map of local correlations (Fig. S2). Processes that were putatively part of a single astrocyte due to correlated activity as seen by the map of local correlations were included in a single ROI. Contamination of the GCaMP6s signals through neuronal signals (GCaMP6f) for dual-layer imaging was negligible, primarily due the spatial separation of pyramidal neurons from the astrocytic imaging layer in the *stratum oriens*. The absence of contamination is evidenced by the correlation function between neuronal and astrocytic activity (Fig. 4a), which does not exhibit a peak at t = 0 s. Due to the known leakiness of the GFAP promotor, also a very sparse set of interneurons was labeled with GCaMP6s. These rare cells (0.7 ± 0.6 cells per FOV%; mean ± s.d. across 19 FOVs from 3 animals) were identified based on their distinct morphology in the mean fluorescence image (see Fig. S2a for an example) and based on their quickly fluctuating signals which were clearly distinct from astrocytic signal time courses (Fig. 1b-c). To illustrate the negligeable effect of these interneurons on our analyses based on mean ΔF/F across the FOV, we computed the fraction of pixels in a FOV covered by interneurons and found it to be much smaller than 1% (0.13 ± 0.19 %; mean ± s.d. across 19 FOVs from 3 animals).

Residual movement artifacts that involved detectable movement along the z-direction were visually detected from the extracted temporal traces as events of both correlated and anti-correlated changes of fluorescence across a majority of the FOV. These events were additionally inspected in the raw movies and blanked for further analyses. Mice for which such strong motion artifacts occurred regularly were not further used for experiments and analyses.

### Behavioral monitoring post-processing

The behavioral video was used to extract several behavioral features. The correlation between subsequent frames was used to compute the movement from two sub-regions that were manually drawn around the mouth and the front paws. The measured metric (1 - correlation) was scaled by the maximum within each session. To extract higher-dimensional face movements, we employed singular value component analysis of face movements^15^ and used a combination of these face movement components to explain astrocytic signals (Fig. S5).

Pupil diameter could be quantified since the infrared laser light of the two-photon system exited the pupil, which in turn appeared bright in the behavioral camera video. The equivalent diameter was computed from the area of the pupil, which in turn was obtained from a simple segmentation of the pupil with standard image processing methods. Briefly, the brightest round object in the ROI covering the eye was extracted from the binarized image using the *regionprops()* function in MATLAB (MathWorks). Dark spots inside the segmented pupil due to reflections were filled and connections of the segmented pupil to the bright upper eyelid were removed by repeated binary erosion and dilation of the segmented pupil. Occasional winking events were manually detected from the extracted pupil diameter traces, confirmed by video inspection, and automatically replaced by values obtained via linear interpolation. The difference between illumination periodicity (from the imaging system, 30.88 Hz) and the video recording rate (30 Hz) resulted in a beating pattern of 0.88 Hz that was also visible in the extracted pupil signal. A template filter based on the 0.88 Hz periodicity was used to remove this illumination-induced component from the extracted pupil diameter. For the analyses in Figs. 3 and 4, where the absolute values of the observed pupil diameter was not important for the analysis, the absolute pupil diameter was used. For analyses associated with Fig. 6, to allow for pooling of pupil diameter values across sessions, pupil diameters were normalized (z-score) for each behavioral session to account for variable lighting conditions.

Similarly, licking was quantified by classical segmentation methods that detected the presence of a bright and large object close to the lick spout (the tongue). The final output was binarized (licking vs. non-licking) based on a threshold of detected object size. The threshold for lick detection was adjusted manually for each session and for each mouse while inspecting the resulting segmentation together with the raw behavioral movies.

The absolute position of the mouse on the treadmill belt was computed using the run speed recorded with the rotary encoder as a relative position signal and the analog IR diode output, which increased when a stripe of reflective tape at the backside of the belt went past the diode, as an absolute position signal.

### Modeling global astrocytic activity with dilated linear regression

To model global astrocytic activity as a function of other variables (*e*.*g*., paw movement, pupil diameter or location), we used a dilated variant of linear temporal regression. Specifically, we averaged time points of the regressor around the to-be-regressed time point into bins that exponentially increased their width with the temporal distance from the current time point. Therefore, the first bin was 1 frame in width, the second 2 frames, the third 4 frames, the fourth 8 frames, and so on, resulting in an effective time window of ±17 s (±512 time points sampled at 30 Hz). This “dilated linear regression” is inspired by “dilated convolutions” used for signal processing and deep neuronal networks^95,96^ and serves to reduce the number of regressors while providing a multi-scale representation of the regressor variable. Such a reduction of the number of regressors is necessary to avoid overfitting. Overfitting is a problem when regressing global astrocytic signals, which provide – due to their slowly changing nature – only relatively few statistically independent data points even for longer recordings.

The vector of such dilated regressors was used to linearly regress the observed global astrocytic activity with the *glmfit()* function in MATLAB. Performance was evaluated on 5-fold cross-validated subsets within each recorded session to exclude overfitting. Performance was measured using the correlation between predicted and true signals across the 5 cross-validated segments of the entire session.

### Modeling global astrocytic activity with a linear differential equation

A standard leaky-integrator differential equation of the form

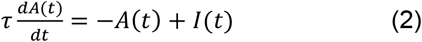

was implemented, with the integration time constant □, the astrocytic activity *A(t)* and the regressor input *I(t)*. Fitting was performed by grid search of the parameter □ and by evaluating the correlation of the recorded signal with the signal obtained by simulation of the differential equation. Since the correlation is not affected by the scaling of the two compared signals, a scaling factor of the input regressor *I(t)* was not necessary. For the evaluation, the first time points until *t* = 2□ were excluded to avoid the influence of the initial conditions during the simulation of the differential equation.

### Self-supervised denoising of calcium movies

Two-photon calcium imaging movies are often dominated by shot noise. Due to the sensitivity of astrocytes to laser-induced heating^97^, we limited the laser power in our experiments to moderate values, at the cost of a decreased signal-to-noise ratio. To enable complex analyses with single-pixel precision despite noisy pixel traces, we used recently developed algorithms for self-supervised denoising of imaging data based on deep networks^98,99^, implemented in the Python scripts of DeepInterpolation^33^. This implementation uses the pixels of the 30 frames prior and after the current frame to denoise the pixels of the current frame. More precisely, the algorithm infers for each pixel the value that is most likely, based on its spatio-temporal surrounding pixels and based on the priors of the networks. To adapt the network priors from previously established neuronal imaging data^33^ to our astrocytic imaging data, we retrained the deep network from scratch for each analyzed session with 10,000 frames of the recording and then ran the trained network on all imaging frames of the respective session. To enable the highest precision and to avoid (anti-)correlated movement artifacts through brain motion, we applied an algorithm for piecewise rigid motion correction on the denoised data^100^. While an entirely rigid motion correction algorithm corrects for most of the movement artifacts, the piecewise rigid motion correction also was able to correct small or nonlinear distortions that happened on faster timescales than the frame rate and required a denoised input.

### Delay maps of astrocytic activity

We used the global astrocytic activity (average across the entire FOV) as a global reference to compute the average delay of each pixel’s signal with respect to this global reference. To extract the average delay, we computed the correlation function between the reference signal and the pixel’s time trace, which was extracted from the denoised movie (described above). The correlation function was normalized such that the zero-lag component was identical with Pearson’s correlation coefficient. Next, the correlation function was smoothed with a 0.5-s window filter and the delay determined by taking the maximum of the correlation function in a -10 … +10 s window around the zero-lag time point. This procedure was repeated for each pixel, resulting in a map of delays. Code in both Matlab and Python to compute delay maps from raw or denoised data is provided, together with sample data, and documented at https://github.com/HelmchenLabSoftware/Centripetal_propagation_astrocytes. Delay maps were computed for each 140-s segment in a session and per-segment maps were combined to a session-map using weighted averaging, with the mean astrocytic activation (ΔF/F) of the entire FOV within the respective segment as the weight used for averaging.This procedure, comprising retraining and application of the self-supervised denoising network and the computation of delay maps, was applied to a total of 12 selected sessions that fulfilled the following criteria: (1) A sufficient amount of global astrocytic activity. For example, sessions with mice that barely moved during the sessions did not exhibit sufficient global astrocytic dynamics for a meaningful analysis. (2) Clearly visible astrocytic somata. In some sessions, especially when the imaging FOV was constrained by simultaneous neuronal imaging in the pyramidal layer but also in other recordings with suboptimal imaging conditions, only gliapil but no clearly detectable cell bodies of astrocytes could be identified, making those recordings less useful for analyses comparing calcium signals in cell bodies vs. distal processes. To validate our approach using delay maps and to show that denoising based on deep networks did not introduce artifacts, we performed a dedicated test imaging session of longer duration (33 min) and with higher laser power to increase SNR. We split the recording in two equal halves and computed reference delay maps for raw and denoised data based on the first half. We then used the second half of the recording to quantify how delay maps based on raw vs. denoised data converged to the reference. We found that delay maps based on denoised data converged faster, whether the reference was computed from raw or denoised data (Fig. S11i-l).

### Identification of centripetally propagating vs. non-propagating calcium events

To extract FOV-wide temporal delay components, the delay map was smoothed with a shape-preserving 2D median filter (filter size 6 μm; Fig. 6a) and then binned by rounding the delay to integer values (…, -2 s, -1 s, 0 s, +1 s, +2 s, …). Pixels with the respective delays across the entire FOV (Fig. 6b) were used to extract the mean time trace from the denoised movie, and this time trace was used to compute ΔF/F (Fig. 6c). A 25 pixel-wide boundary of the FOV was discarded to prevent the contamination by lateral movement artifacts that cannot be corrected at image boundaries. In addition, pixels that contained identified interneurons through ectopic GFAP-driven expression of GCaMP6s were excluded for each FOV using a manually drawn blanking mask.

The temporal delay components (one time trace each for -2 s, -1 s, 0 s, +1 s, etc. pixels) were extracted as raw fluorescence traces and then normalized as ΔF/F values (F_0_ defined as 20% quantile across the session for the respective delay component) to enable a comparison between normalized traces. The average across these traces was used to identify candidate events using the *findpeaks()* function in MATLAB. Since an average across delay components was used as a reference to detect events, it is likely that some gliapil-only events that did not propagate into the soma were not identified as events. The distribution of centripetally propagating and non-propagating events in Fig. 6d is therefore biased towards events that are also visible in processes closer to the soma. All candidate events were visually inspected and corrected. Typical events are shown in Fig. 6e, and other more complex events, which were still considered to be a single event for our analyses, are shown in Fig. S18. The beginning and end of events was estimated as the trough preceding or following an event peak, again proofed by visual inspection. For each event, the mean value of each delay component was extracted, and a linear fit (y = a · x + b) computed, with the extracted mean as y-values and the delay in seconds as x-values (Fig. 6c, inset). The fit parameter a was normalized by b, the fitted value of y at zero-lag x = 0 s, resulting in the “propagation slope” as used throughout the Results section. This normalization has the side effect that errors due to division by small values of b for very small events can result in potentially erroneous large positive or negative slopes; however, at the same time it ensures that the propagation of small-amplitude events is equally considered compared to large-amplitude events. The fit was improved by using the number of FOV pixels that contributed to each delayed trace as fit weights. To define putative astrocytic domains for single-cell analysis of delayed traces (Fig. S19), we manually seeded cell centers based on cell bodies. This analysis could only be performed for sessions in which astrocytic cell bodies could be clearly identified and distinguished from each other and from other structures like end-feet (n = 8 sessions). A watershed algorithm custom-written in MATLAB was used to simultaneously and iteratively expand the domains of all seed points using binary dilation until the domains encountered either the boundary of another domain or reached a distance of ≥35 μm from the seed point.

## Statistics

Unless otherwise indicated, non-parametric, two-sided tests were used (Mann-Whitney rank sum test and Wilcoxon signed-rank test for unpaired and paired conditions, respectively), and no corrections for multiple testing were performed. Since results were often hierarchically organized (e.g., 22 imaging sessions distributed across 4 animals), we computed variability within and across animals and report the values in supplementary table 1. Importantly, within-animal variability was similar to variability across all measurements and in general higher than across-animal variability, suggesting that the single measurements (i.e., from a single imaging session) can be considered to be not strongly influenced by the batch (i.e., animal identity) as quantified by intra-class-correlation^101^ (supplementary table 1). For statistical tests on results based on hierarchically organized data, we used a recently published dedicated toolbox^102^. Box plots used standard settings in MATLAB, with the central line at the median of the distribution, the box at the 25^th^ and 75^th^ percentiles and the whiskers at extreme values excluding outliers.

## Supplementary figures

**Figure S1.**
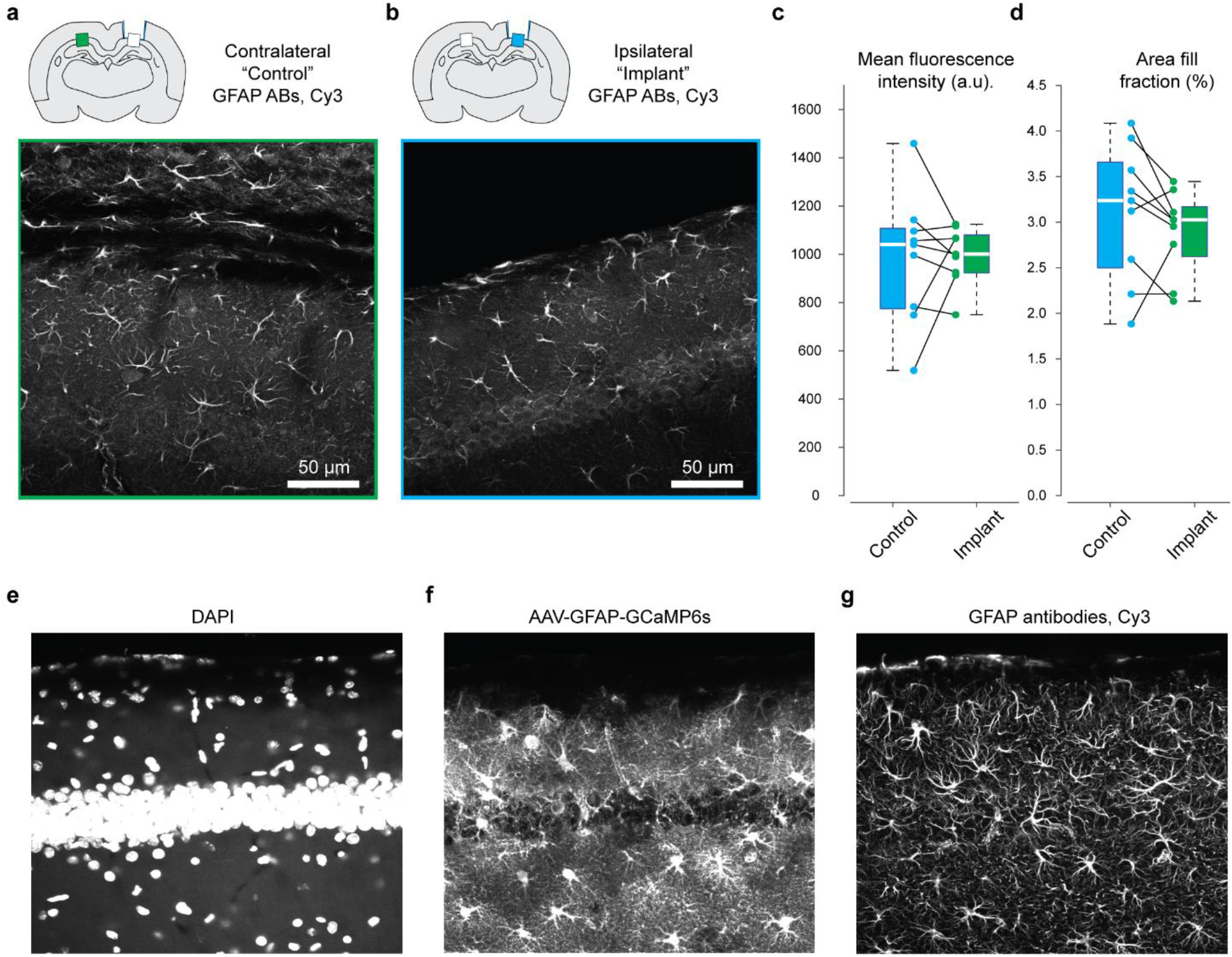
Histology of GFAP expression in astrocytes. **a-b**, Examples of GFAP expression in CA1 *stratum oriens* from the same animal in the location contralateral (a) and ipsilateral (b) to the injection and cannula implantation site. **c**, Quantification of mean fluorescence intensity in regions of interest drawn across the imaging site (*stratum oriens*), compared within the same slice. 9 slices from 3 animals. No statistically significant difference observed (Wilcoxon signed rank test). **d**, Quantification of area covered by astrocytes (thresholding; see Methods) within selected regions of interest. The same threshold was applied to paired images from the same slice. **e-g**, Histology channels of Fig. 1d, shown separately in grayscale. **e**, nuclear stain DAPI. **f**, Virus-induced GCaMP6s expression in hippocampal astrocytes. **f**, GFAP-antibody staining of astrocytes. Maximum intensity projection across 12 μm. While antibodies tended to more heavily stain the distal processes of astrocytes, all astrocytes labeled with the GFAP antibody also exhibited expression with AAV-induced GCaMP6s.

**Figure S2.**
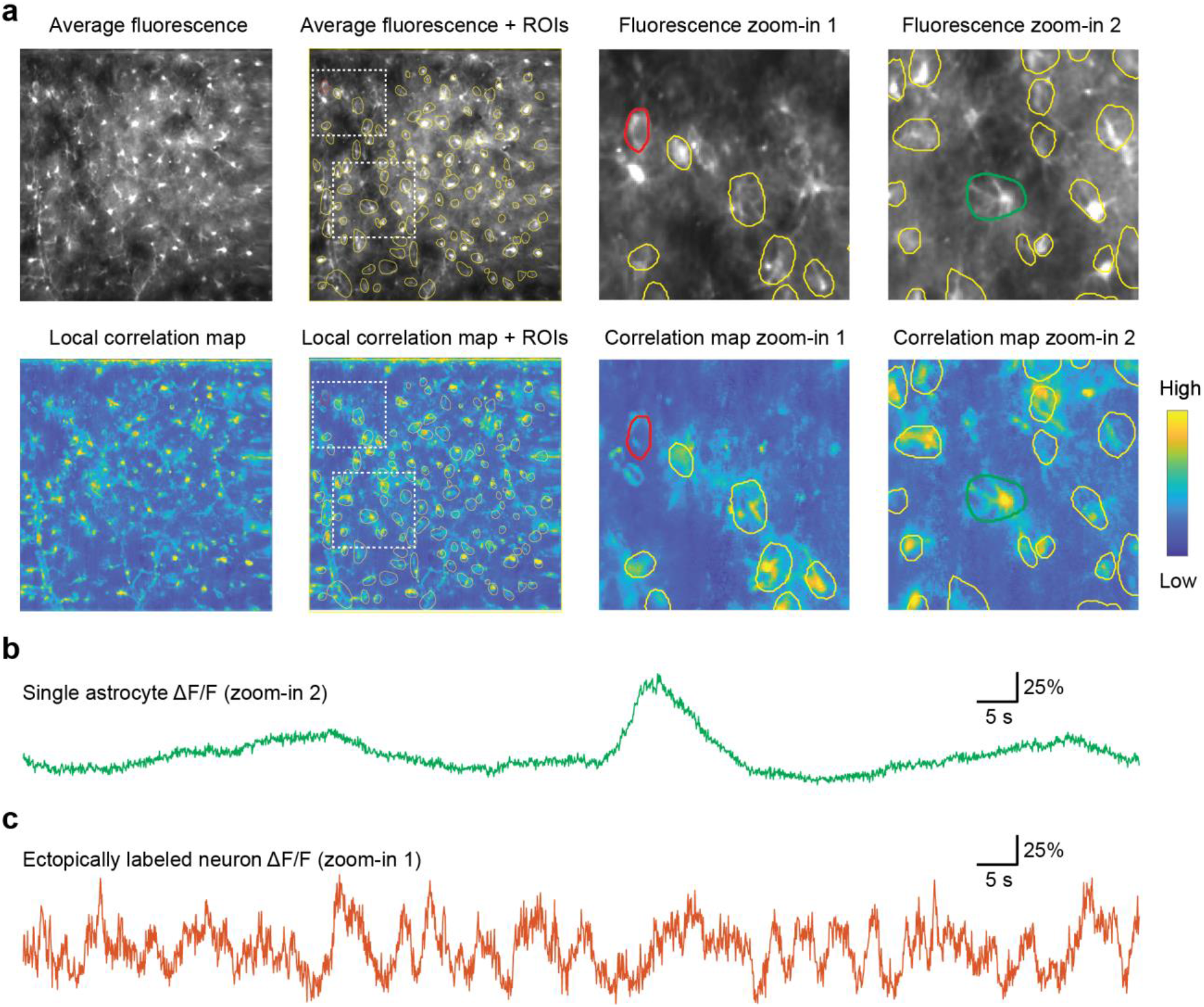
Manual selection of astrocytic active ROIs (related to Fig. 1). **a**, ROIs were selected based on mean fluorescence (structural label, top row) and a map of local correlations (functional label, bottom row). High values in the map of local correlations indicate structural components that are activated in a coherent manner. During manual selection of ROIs, temporal components for manually selected ROIs were displayed (see panels b and c), allowing, *e*.*g*., to quickly recognize and discard ROIs of ectopically labeled interneurons. **b**, Example of a temporal component extracted from an astrocytic ROI centered around a soma (zoom-in 2 from (a), green ROI). **c**, Example of a temporal component extracted from an interneuron ROI (zoom-in 1 from (a), red ROI), displaying the fast-varying nature of neuronal as opposed to astrocytic signals.

**Figure S3.**
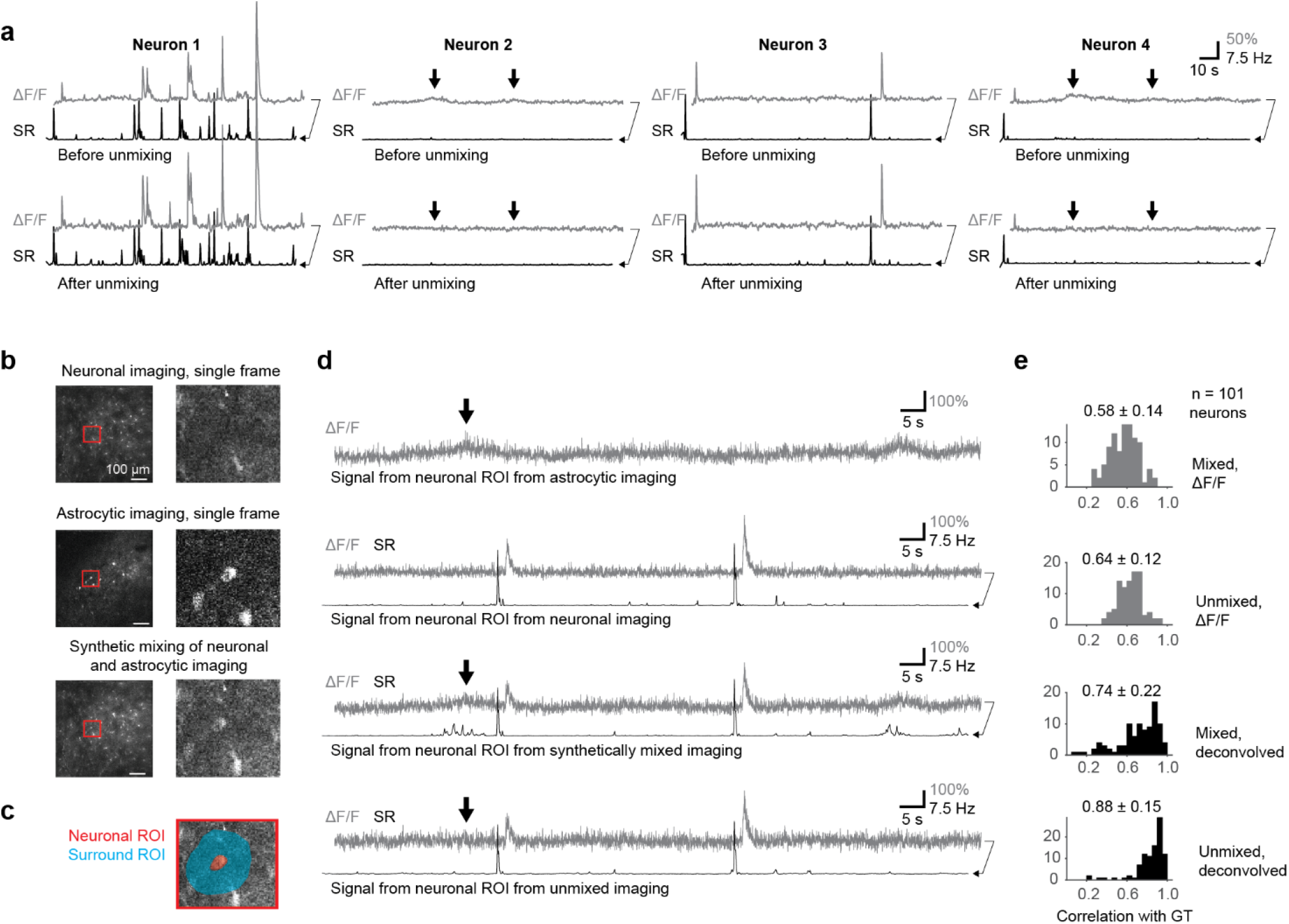
Unmixing of astrocytic signals from neuronal calcium recordings. As astrocytes and neurons were recorded simultaneously using indicators with overlapping non-distinguishable spectra (GCaMP6s and GCaMP6f), we took advantage of the fact that the two cell types were spatially separated (astrocytes in the *stratum oriens*, neurons in the pyramidal layer) and that typical time courses of mean activity patterns were clearly distinct. Astrocytic signals were not visibly contaminated by neuronal signals, as evidenced by the lack of a 0 s-peak in the correlation function (Fig. 4a). Neuronal signals were, however, contaminated by surrounding gliapil. We performed unmixing of astrocytic signals using linear regression of the surrounding gliapil (Methods). To visualize and validate our approach, we show here examples of unmixed neuronal imaging data **(panel a)** and synthetically mixed and afterwards unmixed signals **(panels b-e). a**, Typical examples for neuronal traces (∆F/F and deconvolved spike rate SR) before and after unmixing. For many neurons, unmixing has no clearly visible effect (neurons 1 and 3) since there is only sparsely distributed gliapil in the pyramidal cell layer. For some neurons, contamination can be observed as a slow transient that is clearly distinguishable from calcium transients that originate from neuronal spikes (neuron 2 and 4). In this case, deconvolution, which is trained to detect fast onset transients, removes the slow transients (neuron 2). Slow transients are also removed by unmixing and further suppressed by deconvolution (neurons 2 and 4). Therefore, both unmixing and deconvolution remove astrocytic contamination when present. Deconvolved traces are shifted to the left for visualization purposes. **b**, Single frames of separate neuronal imaging data (top), astrocytic imaging data (middle) and the synthetically mixed data (bottom). **c**, Illustration of neuronal soma ROI (red) and surrounding contamination ROI (blue). **d**, Time traces derived – from top to bottom – from astrocytic imaging, from neuronal imaging, from synthetically mixed imaging data and from the unmixed imaging data, with the associated deconvolved traces in black (if applicable). Deconvolved traces are shifted to the left for visualization purposes. **e**, Distributions of correlation values for ∆F/F traces before and after unmixing (correlation to the ∆F/F traces of the original neuronal imaging data before synthetic mixing) and of correlation values for deconvolved traces obtained from data before and after unmixing (correlation to the original deconvolved traces). Each data point of this distribution corresponds to a neuron and the correlation of its extracted time trace with the reference extracted from the original neuronal imaging data. Together, both unmixing and deconvolution contribute to recover ground truth. All differences between the four distributions (mixed vs. unmixed ∆F/F; mixed vs. unmixed deconvolved; mixed ∆F/F vs. mixed deconvolved; unmixed ∆F/F vs. unmixed deconvolved) in panels (e) were statistically significant (paired Wilcoxon test, p<10^-10^ after Bonferroni correction for multiple comparisons).

**Figure S4.**
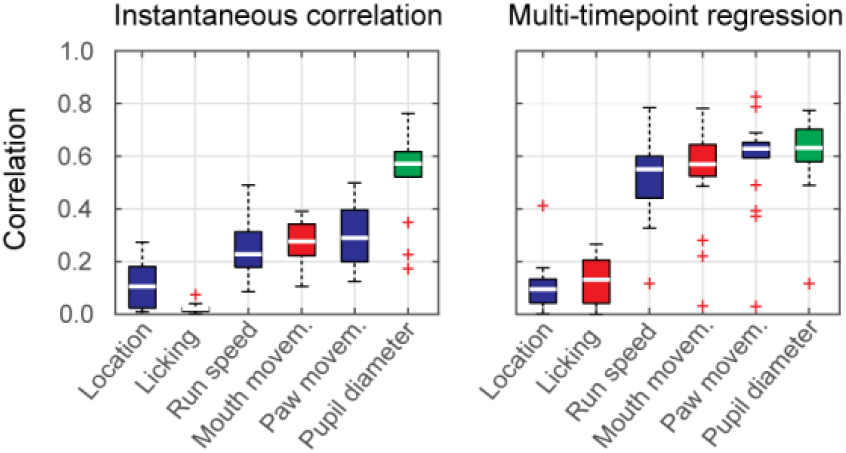
Global astrocytic activity can be well explained by past behavior, mean neuronal spike rate or pupil diameter. Same analysis as in Fig. 3c,d but for a different set of animals, in which calcium imaging was performed in astrocytes only (thus lacking the predictions by neuronal ∆F/F and SR; 19 imaging sessions from 3 animals). Results are similar to the results for the 4 animals shown in Fig. 3c,d.

**Figure S5.**
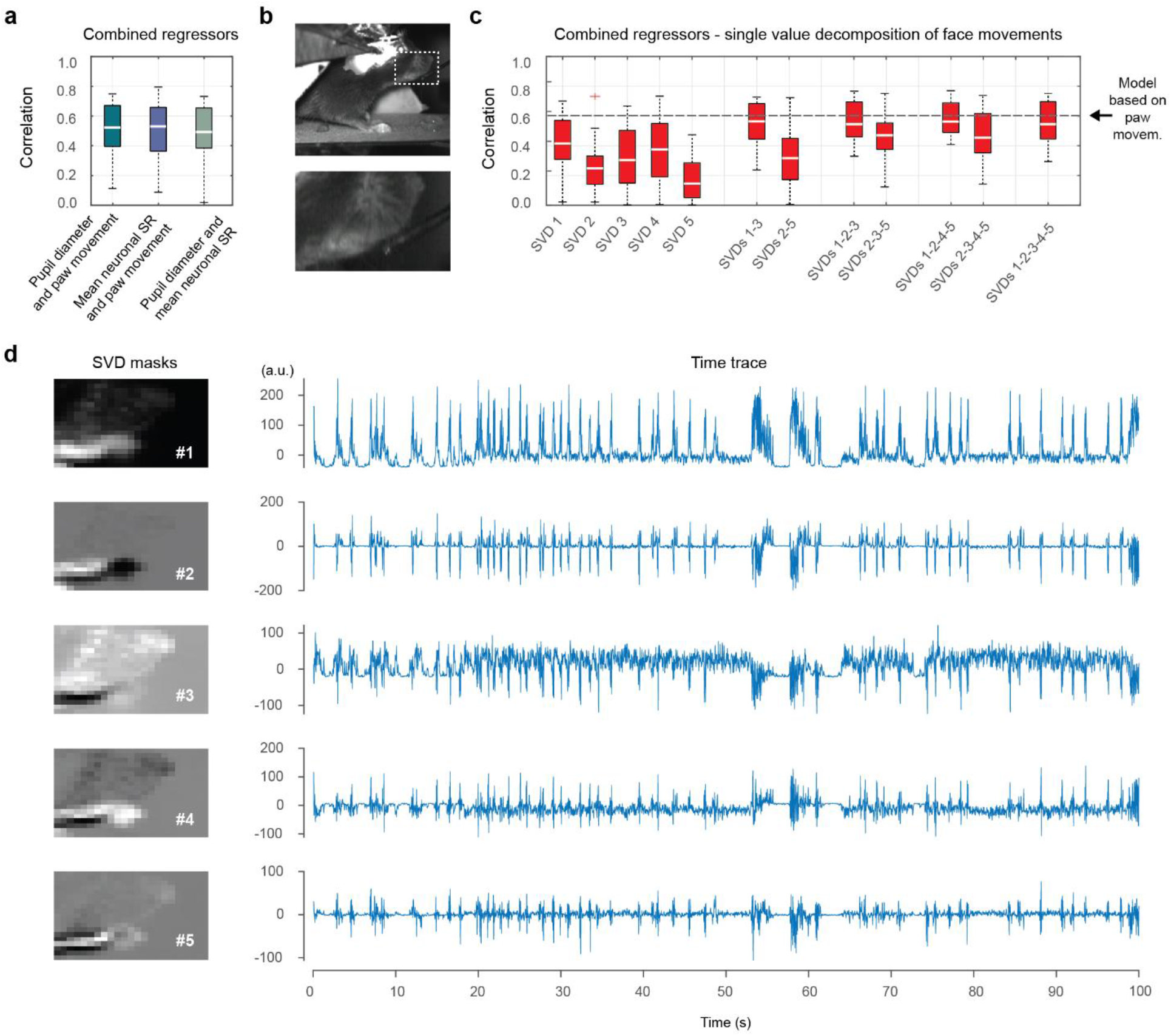
Predictions of global astrocytic activity based on combined and higher-dimensional regressors. **a**, Using combined regressors (*e*.*g*., pupil diameter and paw movement) instead of a single regressor (*e*.*g*., only pupil diameter) did not improve predictions, as measured by correlation values (y-axis). **b**, Using the face recording and its movement components (SVD) to explain global astrocytic activity. **c**, Predictions when combining single or multiple regressors (SVDs) extracted from face movement using Facemap^15^. Example spatial and temporal components are shown in panel (d). No combination of multiple face movement components resulted in predictions better than from simple paw movement regression (single-instead of multi-dimensional regressor). All results measured across 22 sessions for 4 animals. **c**, Visualization of spatial and temporal components for mouth movement analysis for an excerpt of a single imaging session. The lack of improvement when using multiple or higher-dimensional regressors (panels (a) and (c)) is likely due to two effects. First, the auto-correlated astrocytic signal yields relatively few independent data points that result in overfitting when increasing the number of regressors and deterioration of cross-validated predictions. Second, distinct regressors lead to highly similar predictions due to redundant signals (Fig. 3e,f), resulting in lack of strong improvements when combining regressors.

**Figure S6.**
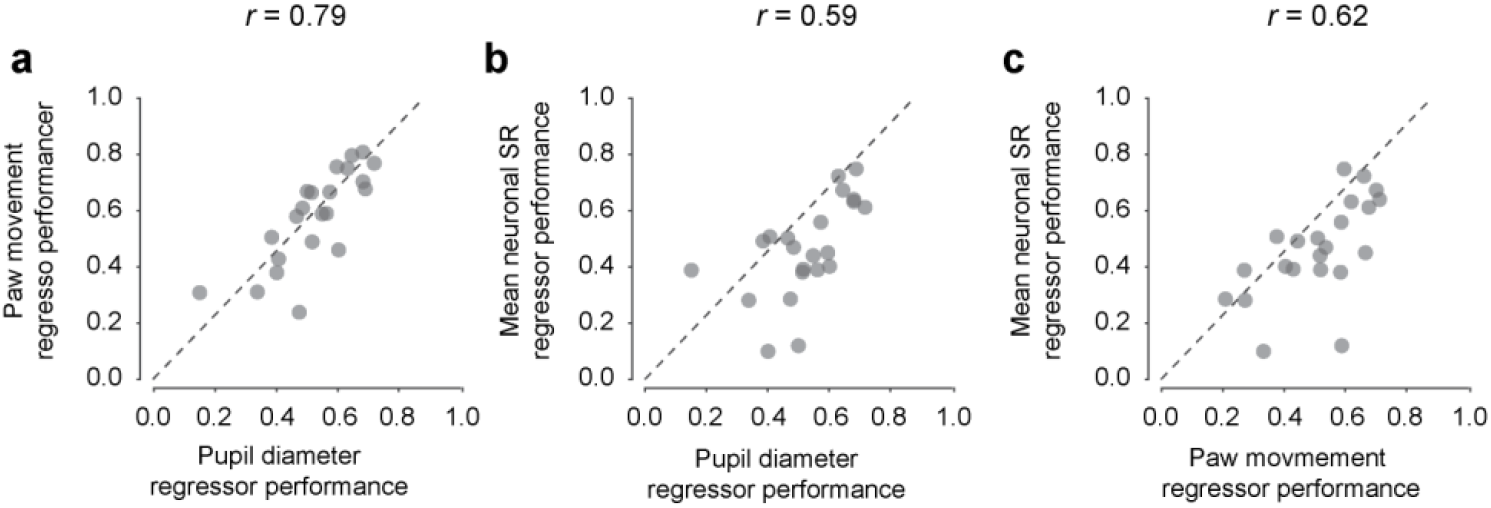
Performance for different regressors across sessions is highly correlated. Extension of Fig. 3g. **a**, Performance of pupil diameter vs. paw movement as regressors. **b**, Performance of pupil diameter vs. mean neuronal spike rate (SR) as regressors. **c**, Performance of paw movement vs. mean neuronal spike rate as regressors. For all panels, a single data point represents the average performance of the cross-validated dilated linear regression averaged for an imaging session. Values (*r*) indicate the correlation between the two regressors’ performances. The dashed lines represent the identity relationship.

**Figure S7.**
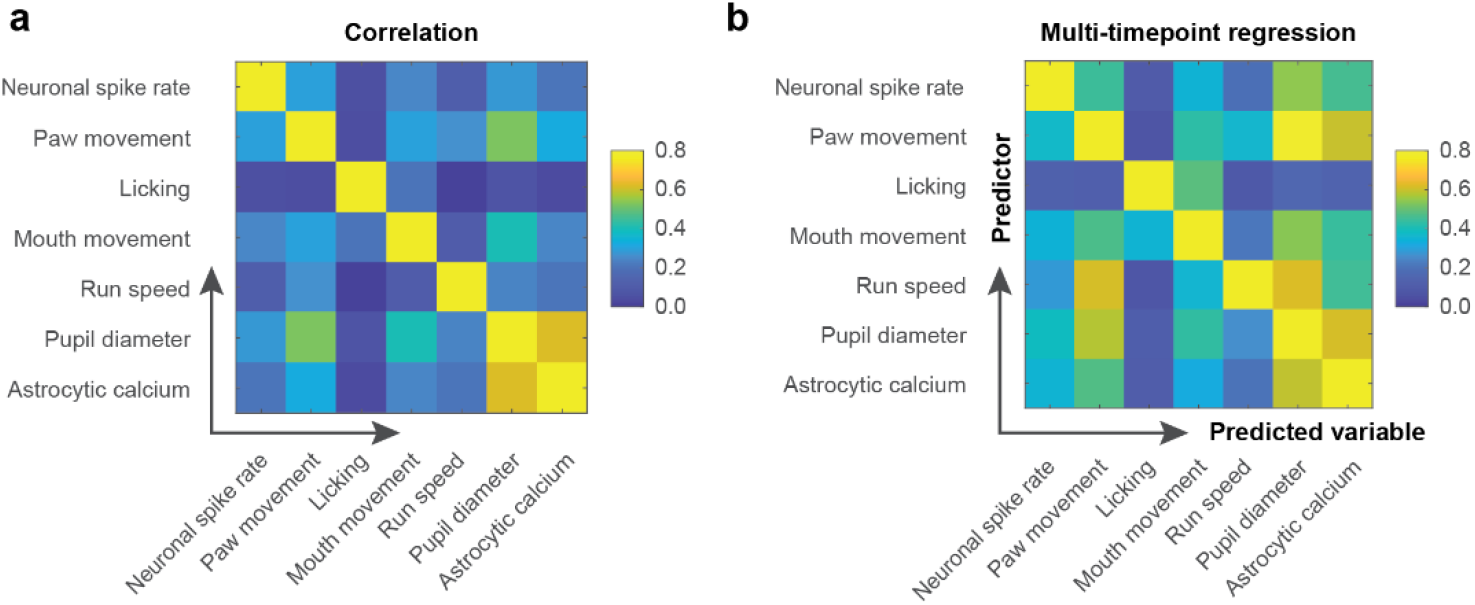
Instantaneous and time-dependent co-variation of behavioral, neuronal and astrocytic signals. Extension of Fig. 3c-d. **a**, Correlation of each observed variable with each of the other observed variables. Most pairs of variables (with few exceptions: pupil diameter vs. astrocytic calcium and paw movement vs. pupil diameter) are only weakly correlated. The right-most column corresponds to the median values in Fig. 3a. **b**, Multi-timepoint regression as described in Fig. 3 was used to predict a variable (column) from each of the other variables (row). The right-most column corresponds to the median values in Fig. 3d. By the definition of correlation and correlation functions (concepts that are directly related to variance explained), slowly-varying signals (*e*.*g*., astrocytic mean activity or pupil diameter) are easier to predict than quickly varying signals (*e*.*g*., mean neuronal activity). It is therefore natural, and should be considered for any interpretation, that a quickly varying signal like mean neuronal activity explains a large fraction of the variance of astrocytic calcium but not vice versa. All values were computed as medians across 22 sessions from 4 animals.

**Figure S8.**
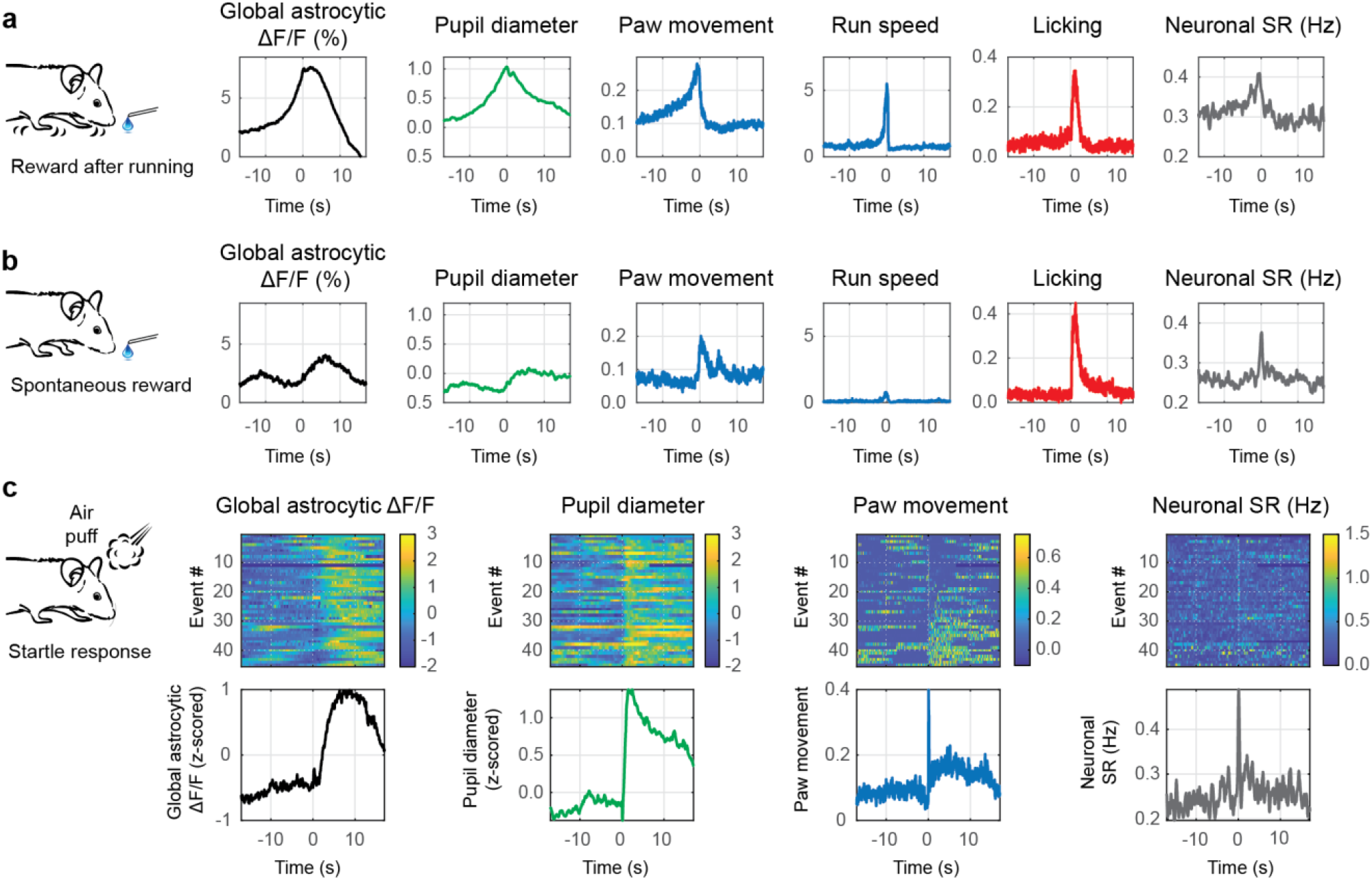
Pupil diameter but not body movement is reliably coupled to global astrocytic events. **a**, Event-triggered average traces for rewards that are obtained at a reward location after running, aligned to reward consumption (average across n = 260 events). All variables with arbitrary scaling (a.u.) unless otherwise indicated. **b**, Event-triggered average traces for spontaneous rewards, aligned to reward consumption. (n = 168). All variables with arbitrary scaling (a.u.) unless otherwise indicated. **c**, Event-triggered traces for air-puff induced startle responses (top: heatmap of individual traces for 45 events; bottom: average across events). Single traces for ∆F/F and pupil diameter are z-scored. Traces in heatmaps are sorted by increasing paw movement during the first 3 s after the air puff.

**Figure S9.**
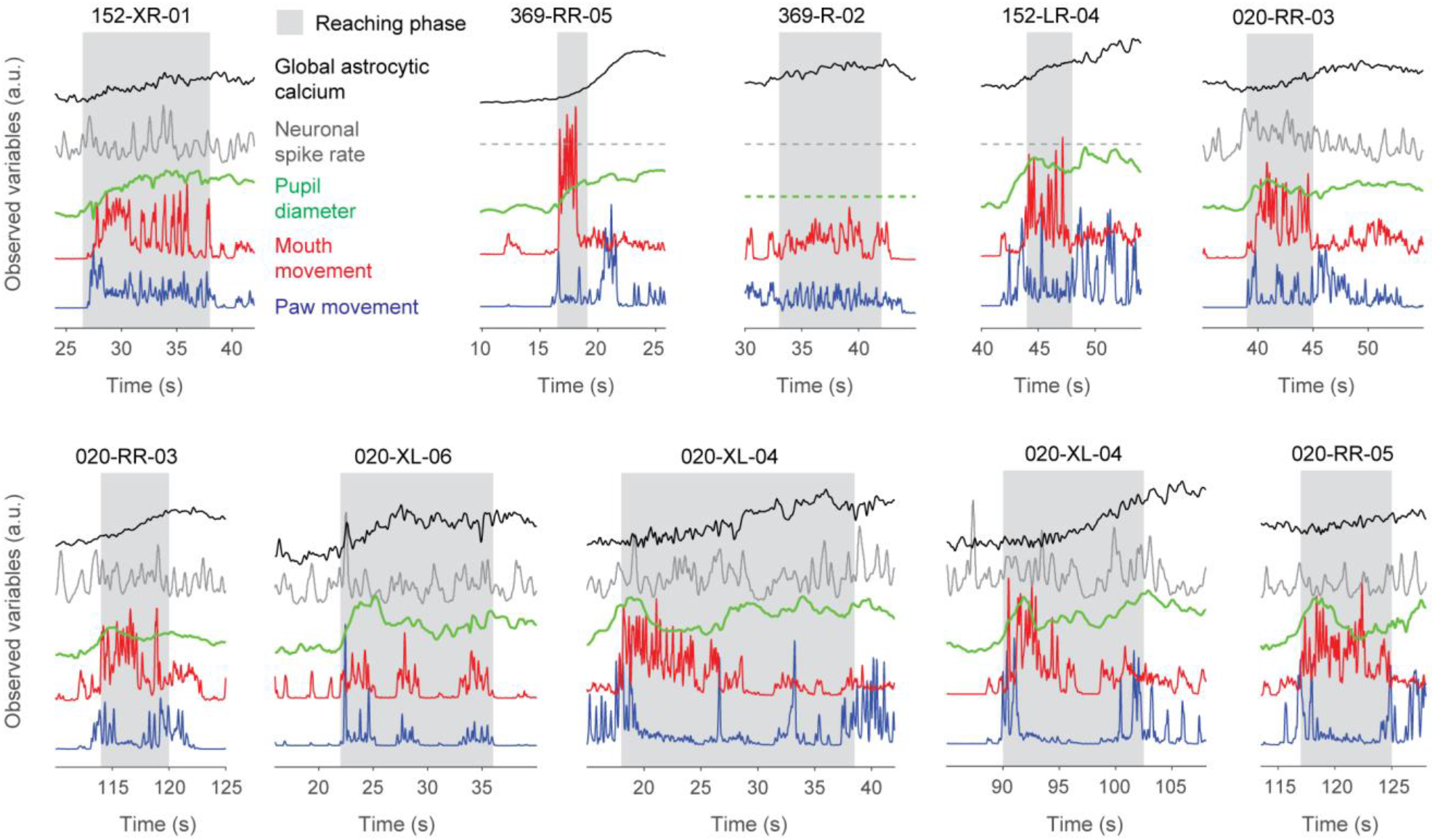
Global astrocytic calcium increases during non-locomotor movements (reaching). Simultaneously recorded variables are color-coded and represented with dashed flat lines if no recordings were available. Panel titles indicate mouse ID (*e*.*g*., 152-XR for the first panel) and the respective imaging session (*e*.*g*., 01). The (manually defined) grasping phase is highlighted as gray background. Reaching, which resulted in paw movement close to the lick spout in proximity of the mouth, is usually visible as deflections of the “mouth movement” (red). All events in this figure are shown, together with the recorded variables and the behavioral video in Movie 5.

**Figure S10.**
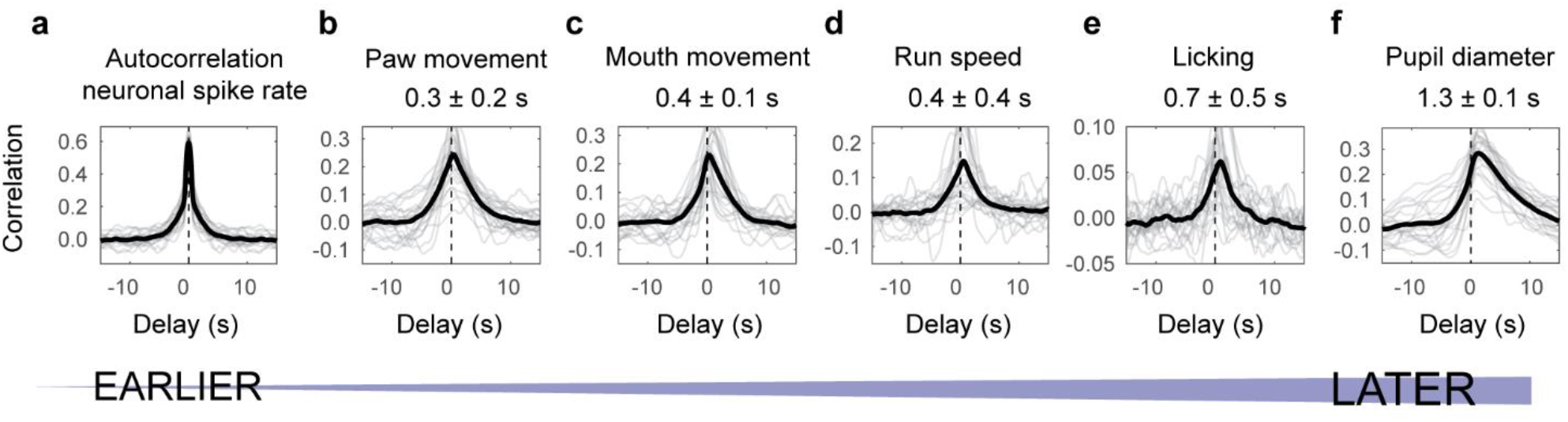
Temporal sequence of behavioral variables with respect to mean neuronal spike rate. Analogous to Fig. 4 but using deconvolved neuronal spike rate instead of global astrocytic activity as the reference signal. A peak of the correlation function with positive lag indicates that the neuronal spike rate peaked on average earlier than the inspected variable. Grey traces are correlation functions extracted from single sessions, black traces are averages across sessions. The delays indicated are median values ± standard error across sessions (n = 22 sessions across 4 animals).

**Figure S11.**
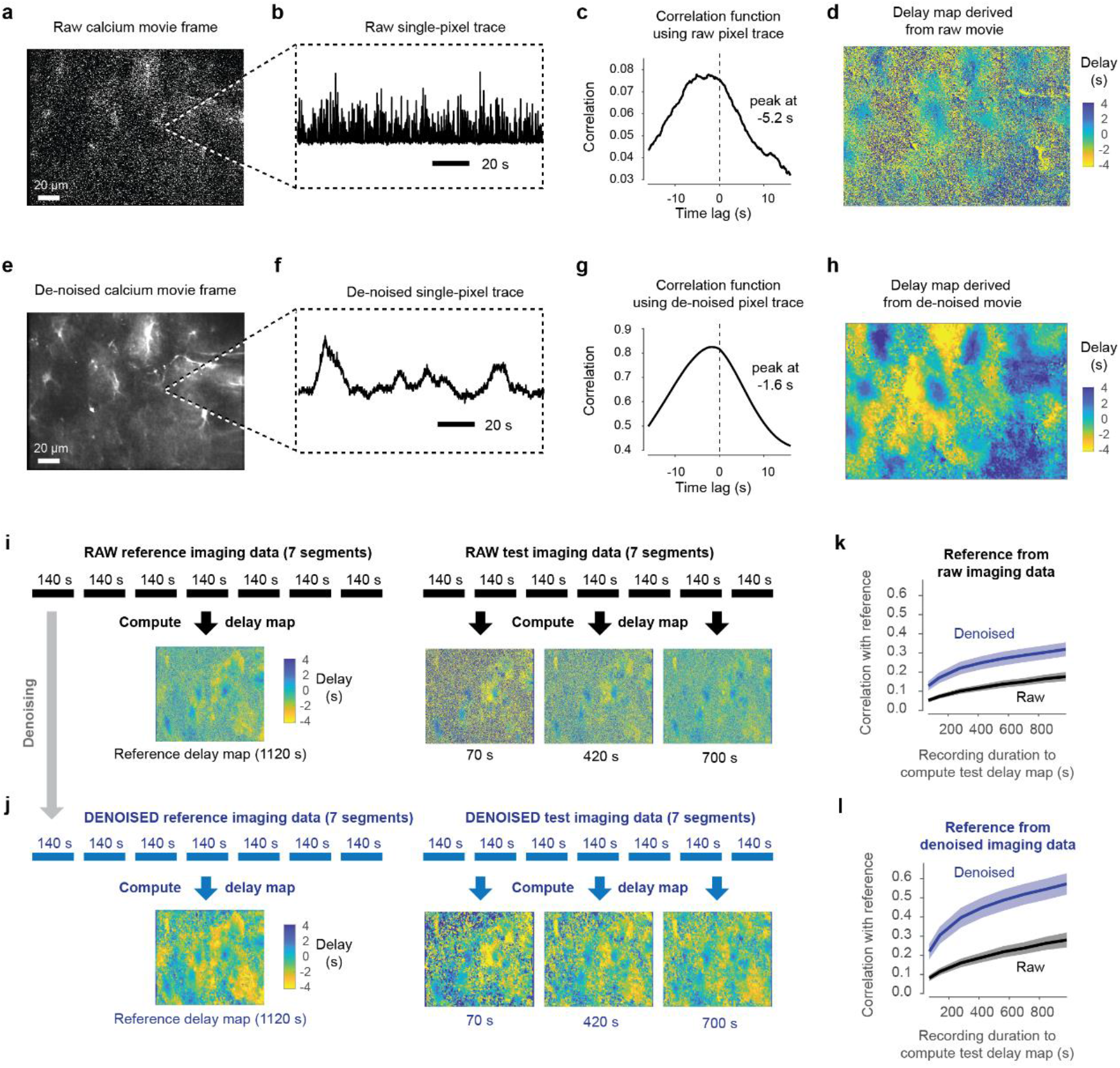
Pixel-wise computation of delays is improved by self-supervised de-noising. Illustration of the workflow (panels a-h). The top row (a-d) shows the results of the processing pipeline as described in Fig. 5e but without de-noising of the raw movie. The bottom row (e-h) shows the results using the de-noised movie **a**, Single frame of the raw movie. **b**, Single-pixel trace extracted from the raw movie in (a). **c**, Correlation function computed from the pixel trace in (b) with the global astrocytic activity across the entire FOV. The peak of the correlation function is, in this case, detected at -5.2 s. **d**, Delays for all pixel traces, mapped onto the imaging FOV. **e-h**, Same as (a-d) but based on the de-noised movie. The resulting delay map (h) is less noisy than in (d). **Validation of delay maps based on raw vs. denoised data (panels i-l)**. A recording from the same FOV comprising 14 segments of each 140 s duration was acquired and split into two equal halves. The first half was used to compute a reference delay map. A second delay map (test delay map) was computed based on a varying number of segments (0.5 - 7 segments) and then compared to the reference map. **i**, Reference map and example test delay maps computed from the raw recording are shown. **j**, Reference map and example test delay maps computed from the denoised recording are shown. **k**, Similarity of test delay maps to the reference map computed from the *raw* recordings. The similarity increases with the number of segments used to compute the test delay map but is always higher when computed from denoised recordings. Shaded corridors indicate the standard deviation across 7 imaging FOVs. **l**, Similarity of test delay maps to the reference map computed from the *denoised* recordings. Shaded corridors indicate the standard deviation across 7 imaging FOVs. Again, the similarity increases with the number of segments used to compute the test delay map but is always higher when computed from denoised recordings. These analyses show that delay maps converge (1) when more frames are used for their computation and (2) when the underlying recordings are denoised.

**Figure S12.**
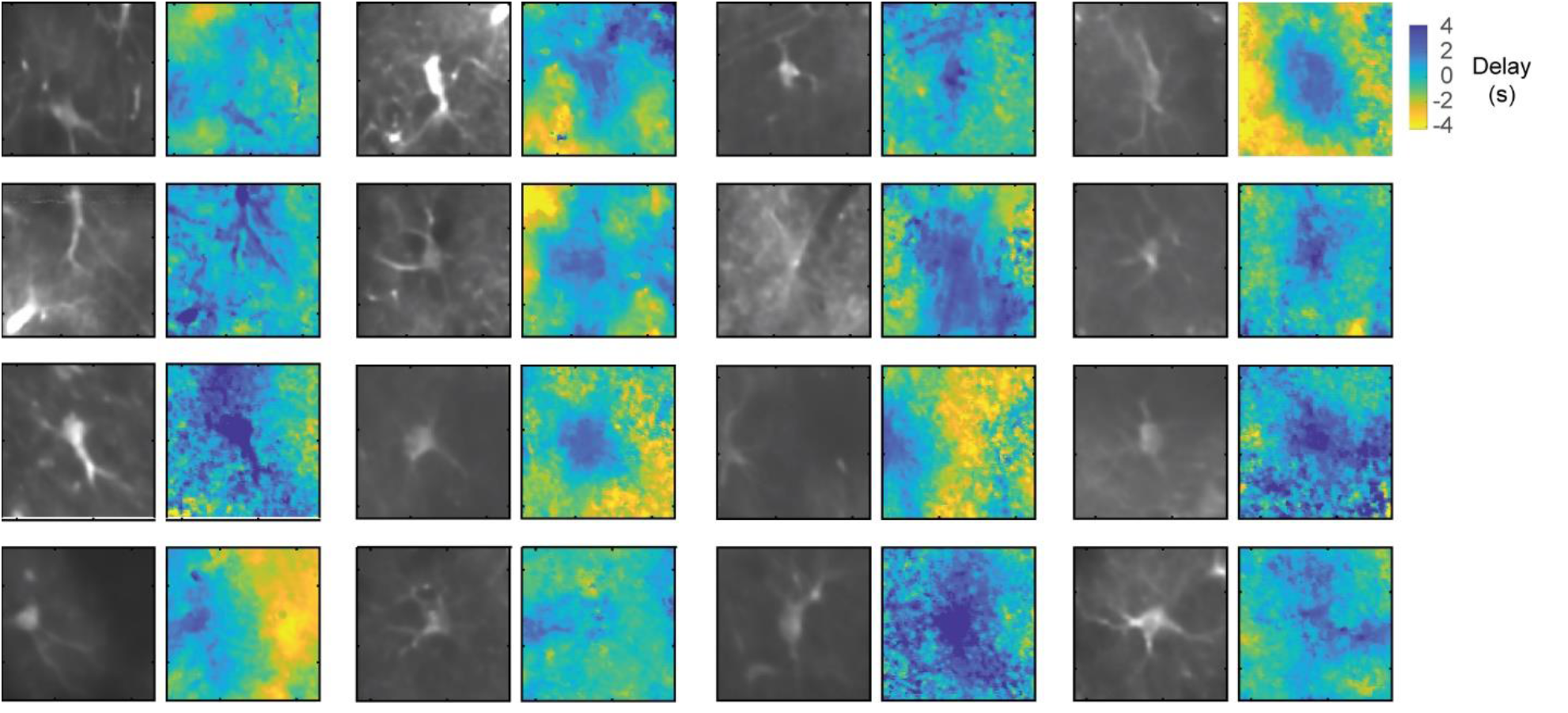
Further examples of isolated astrocytes (fluorescence average, left) together with the local delay maps (right). Extension of Fig. 5h. The color code (as in Fig. 5) indicates propagation of activity from distal to somatic compartments on a timescale of seconds. The side length of each FOV excerpt is approximately 55 μm.

**Figure S13.**
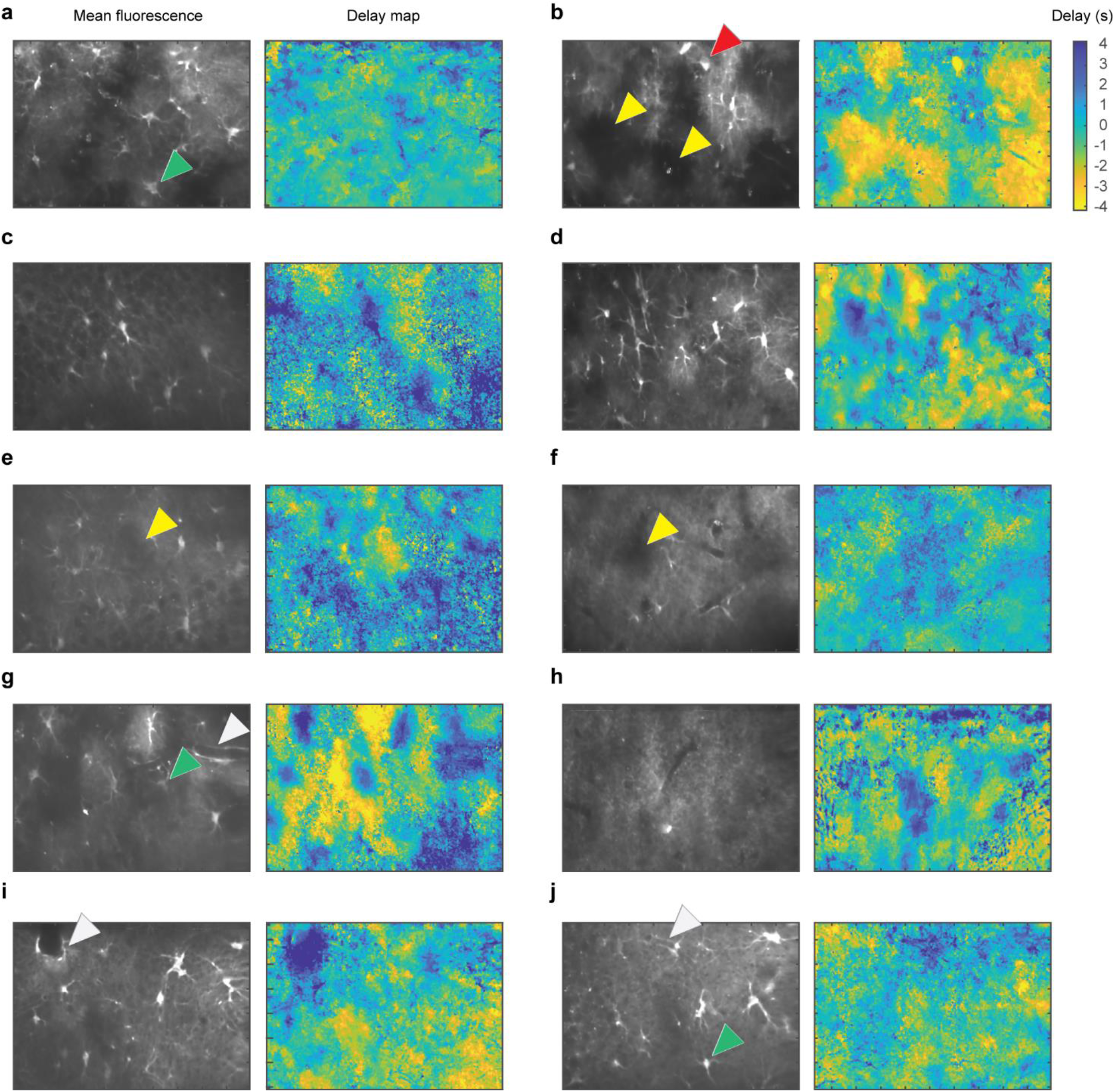
Examples of delay maps. **a-j**, Maps of delays of astrocytic signal with respect to the global mean fluorescence, computed as described in Fig. 5 and Fig. S11. Each delay map represents an entire imaging FOV (40x objective, 200 μm side length in x-direction). Yellow arrow heads highlight regions that are devoid of somata and thick processes, therefore mostly containing fine gliapil processes. Green arrow heads highlight astrocytic somata that are, unlike other soma examples shown in Fig. 5h and Fig. S11, not activated in a delayed manner with respect to the global mean activation. White arrow heads highlight astrocyte processes around blood vessels, exhibiting a delayed activation with respect to the global mean activity. The red arrow head highlights an ectopically labeled interneuron; such interneuron pixels were blanked for the analysis in Fig. 6.

**Figure S14.**
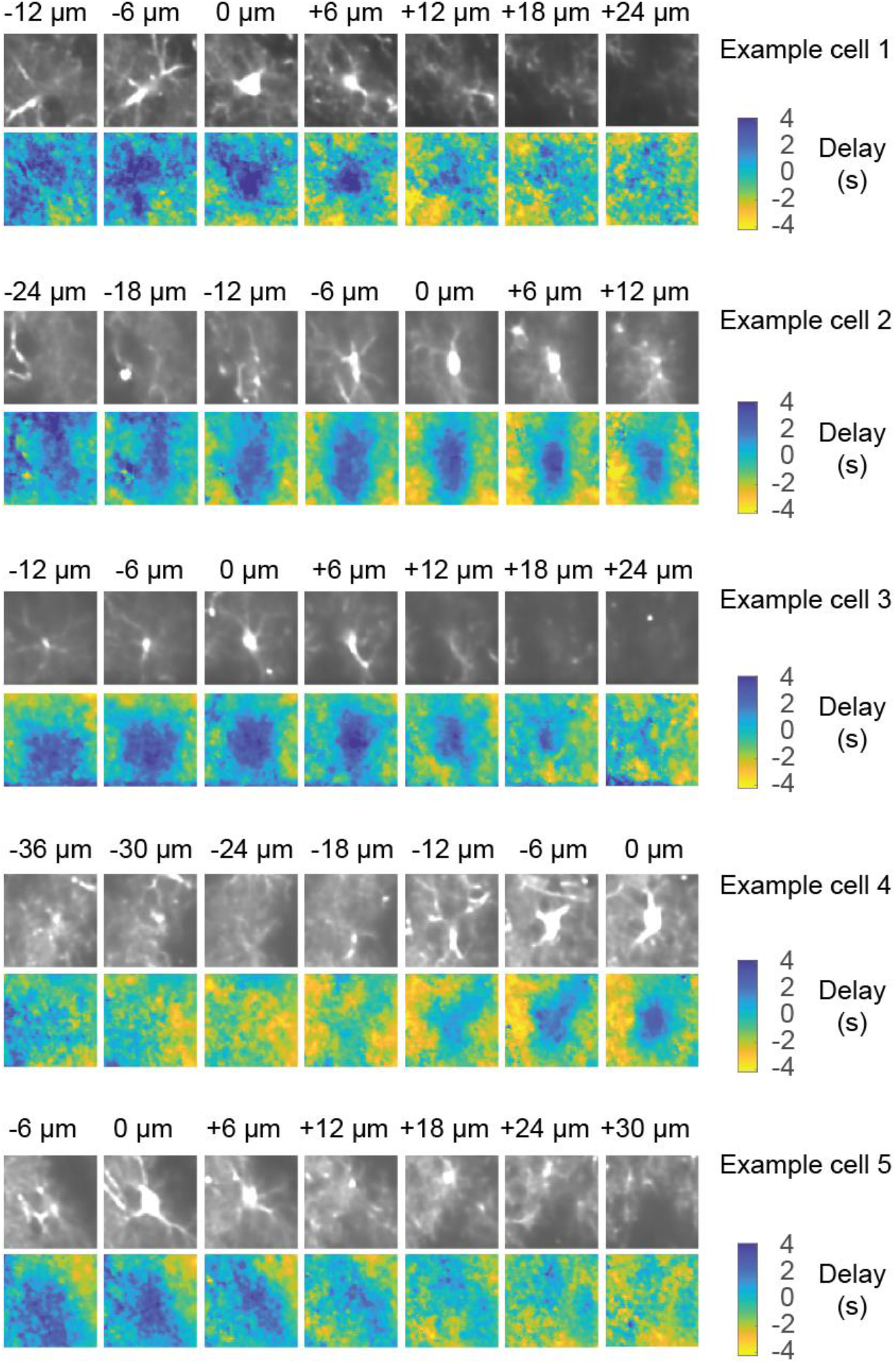
Further examples of 3D delay maps based on dense multi-plane calcium imaging. Extension of Fig. 5f. The 3D delay map exhibits the longest positive delay in the imaging plane with the center of the soma. The putative center of the soma is labeled as the “0 μm” imaging plane. The side length of a single tile is approx. 55 μm. Example cells are taken from two imaging sessions in two mice.

**Figure S15.**
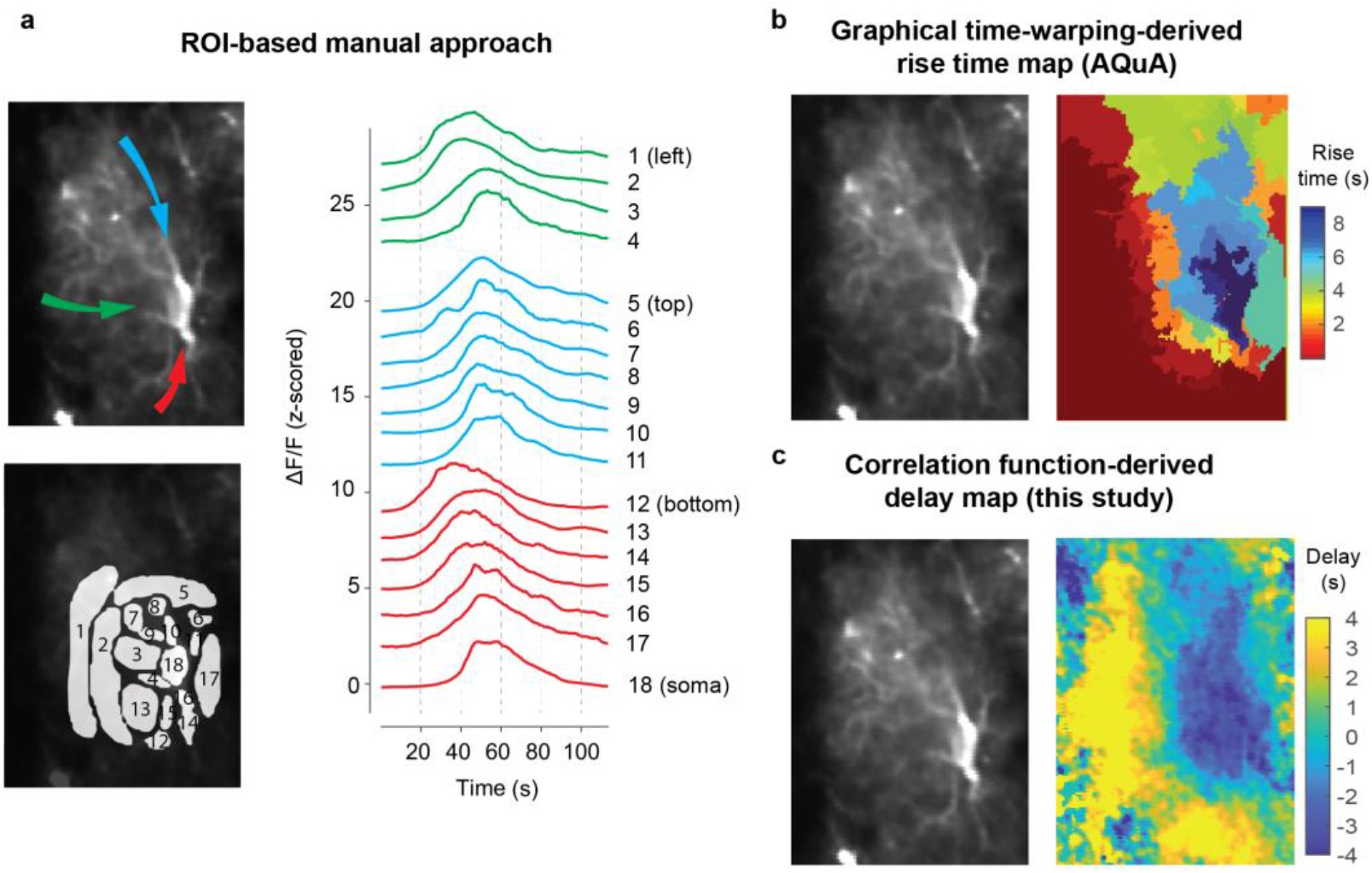
Centripetal propagation replicated using diverse methods. This event in this specific astrocyte was chosen for a comparative analysis since the spatio-temporal dynamics were clearly distinct by eye. Only in this case, the alternative methods (ROI-based approach, AQuA) yielded useable results. The event is therefore not representative but showcases that under conditions of excellent SNR and clearly discernible spatio-temporal events, our approach based on correlation function-derived delay maps is consistent with other analysis approaches. **a**, ROI-based approach. ROIs were manually drawn with an interactive ROI-tool (Rupprecht et al., Neuron, 2018) based on spatially distinct time traces. The ROIs were manually grouped to reveal three distinct pathways of calcium signal propagation towards the soma (along the blue, green and red arrows). Earliest activation is seen from the red processes (ROIs 12-17) and the green processes (ROIs 1-4), while the blue processes (ROIs 5-11) are activated only slightly before the soma. **b**, Rise time map derived using the graphical time-warping method implemented in AQuA (Wang et al., 2019). Briefly, parameters in AQuA were adjusted to label only one single event that comprises the entire astrocyte and its processes. To make this work with AQuA, which is not designed to detect such global events, the denoised movie was fed into the toolbox in temporally reversed order. From the event, toolbox parameters were adjusted until sub-events were reasonably large in their spatial extent and not noisy. Then, the rise time map based on graphical time warping was extracted. NaN values in the resulting rise time map were filled by valid values of nearest neighboring pixels. **c**, Delay map based on the maximum of pixel-wise correlation functions computed between the respective pixel trace and the global mean activity, as described in Fig. 5

**Figure S16.**
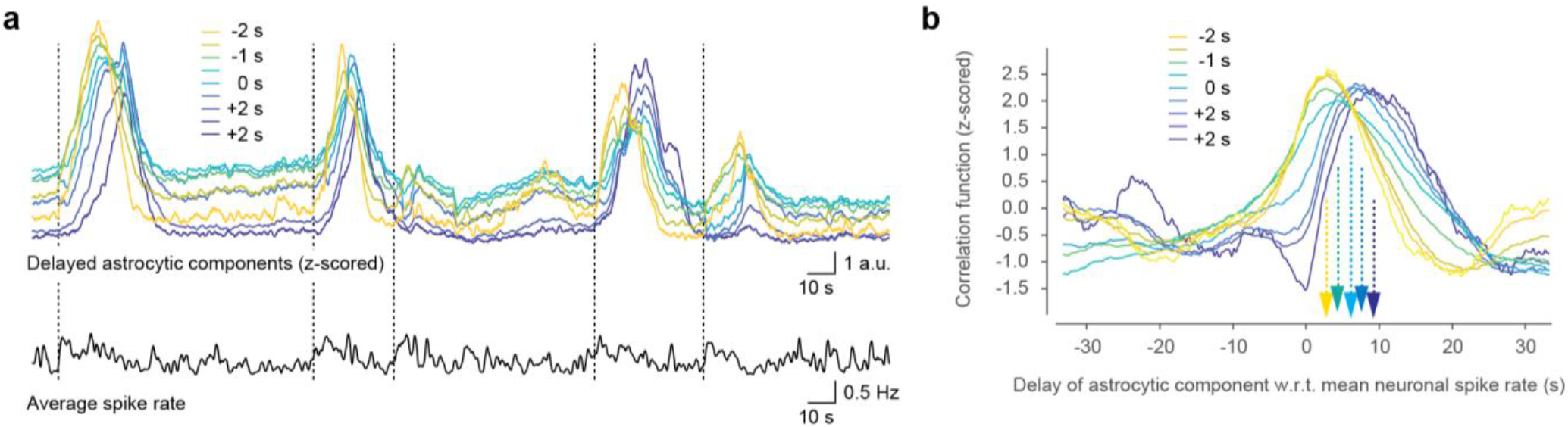
Early and late astrocytic components and neuronal activity, related to Fig. 6c. **a**, Example of delayed components extracted from a FOV as defined in Fig. 6 (colored traces) together with the simultaneously recorded average deconvolved spike rate (black, below). Prior to onset of visually detectable astrocytic events (dashed vertical lines), neuronal spike rates tend to be transiently increased. Also the earliest astrocytic component (yellow) is increased only after this transient increase of neuronal activity. Astrocytic traces were z-scored for better visualization of dynamics. **b**, Separate correlation functions of early and late astrocytic components with simultaneously recorded mean spike rates (average across 12 sessions from 4 mice; averaging was performed in a weighted manner according to the number of pixels associated with a respective delayed component). The correlation functions between 2.5 s (earliest component) and 9.0 s (latest astrocytic component) after neuronal activity. Due to the long auto-correlation time of astrocytic and also neuronal activity (Fig. 4), only the peak of the correlation functions can be meaningfully interpreted.

**Figure S17.**
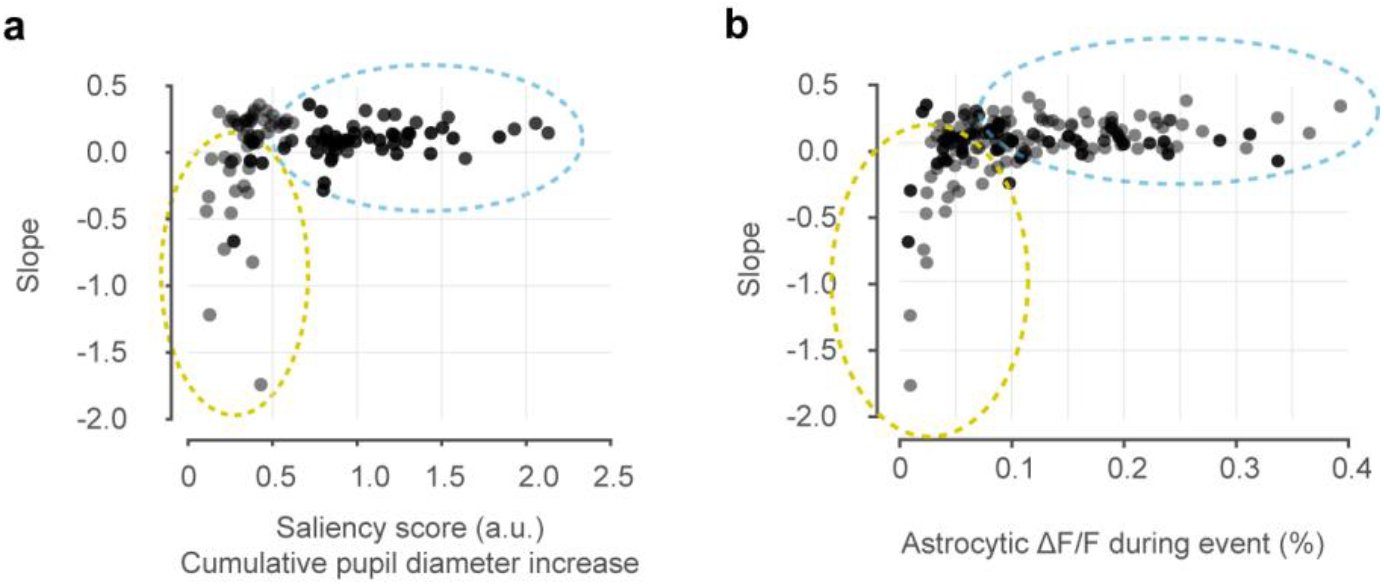
Slope of astrocytic global events as a function of calcium event magnitude and of a ‘saliency score’. Panels are analogous to panels Fig. 6d, but use the saliency score (a) or the astrocytic mean ΔF/F value (b) instead of pupil diameter as the dependent variable (x-axis). The saliency score is computed from the sum of the rectified derivative of the measured pupil diameter in the time window during the event, but shifted to the past by 2 seconds. The score therefore measures the amount of (positive) pupil change, instead of pupil diameter itself.

**Figure S18.**
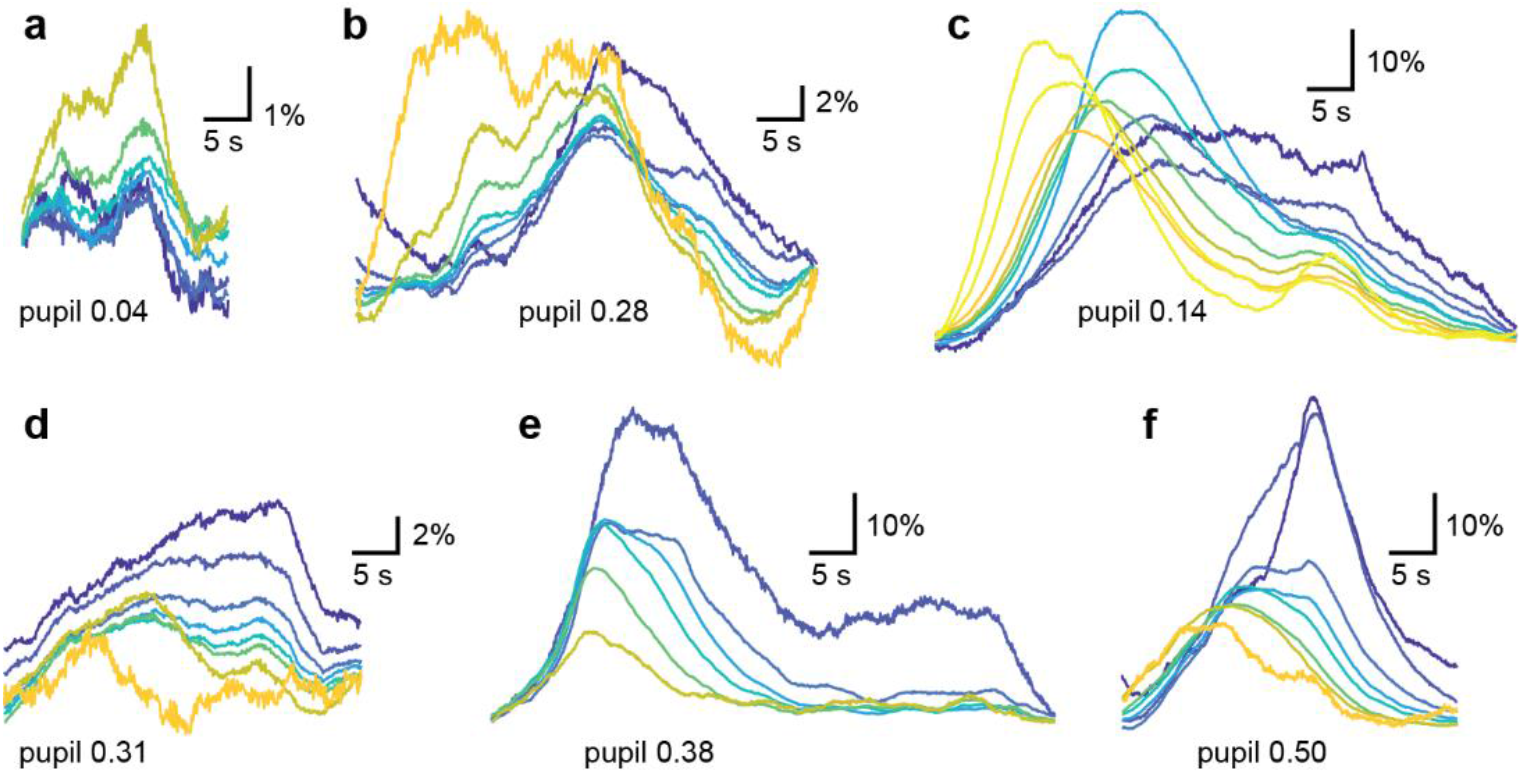
Example events, extension of Fig. 6e. These examples highlight the diversity of astrocytic events that cannot always be well captured as single-phase events. Note the different y-axis scale bars. The x-axis scale bars are identical across events. **a**, Example of a fading event (as defined in Fig. 6) that is barely visible except in the gliapil (yellow). **b**, Biphasic gliapil event. The first gliapil activation (yellow) does not result in centripetal propagation of calcium activity, but a second peak of gliapil activity manages to do so. **c**, Another example of biphasic gliapil activation. The first peak results in centripetal propagation, while the second gliapil peak is only weakly reflected by somatic calcium. **d**, Another example of biphasic gliapil activity. Slow somatic integration barely reflects the faster gliapil fluctuations. **e**, Example of very prominent centripetal propagation, resulting in persistent activation of the somatic region for 10s of seconds. Note the relatively high pupil value. **f**, Similarly prominent centripetal propagation, resulting in a striking display of longer-lasting somatic calcium activity. Note the relatively high pupil value.

**Figure S19.**
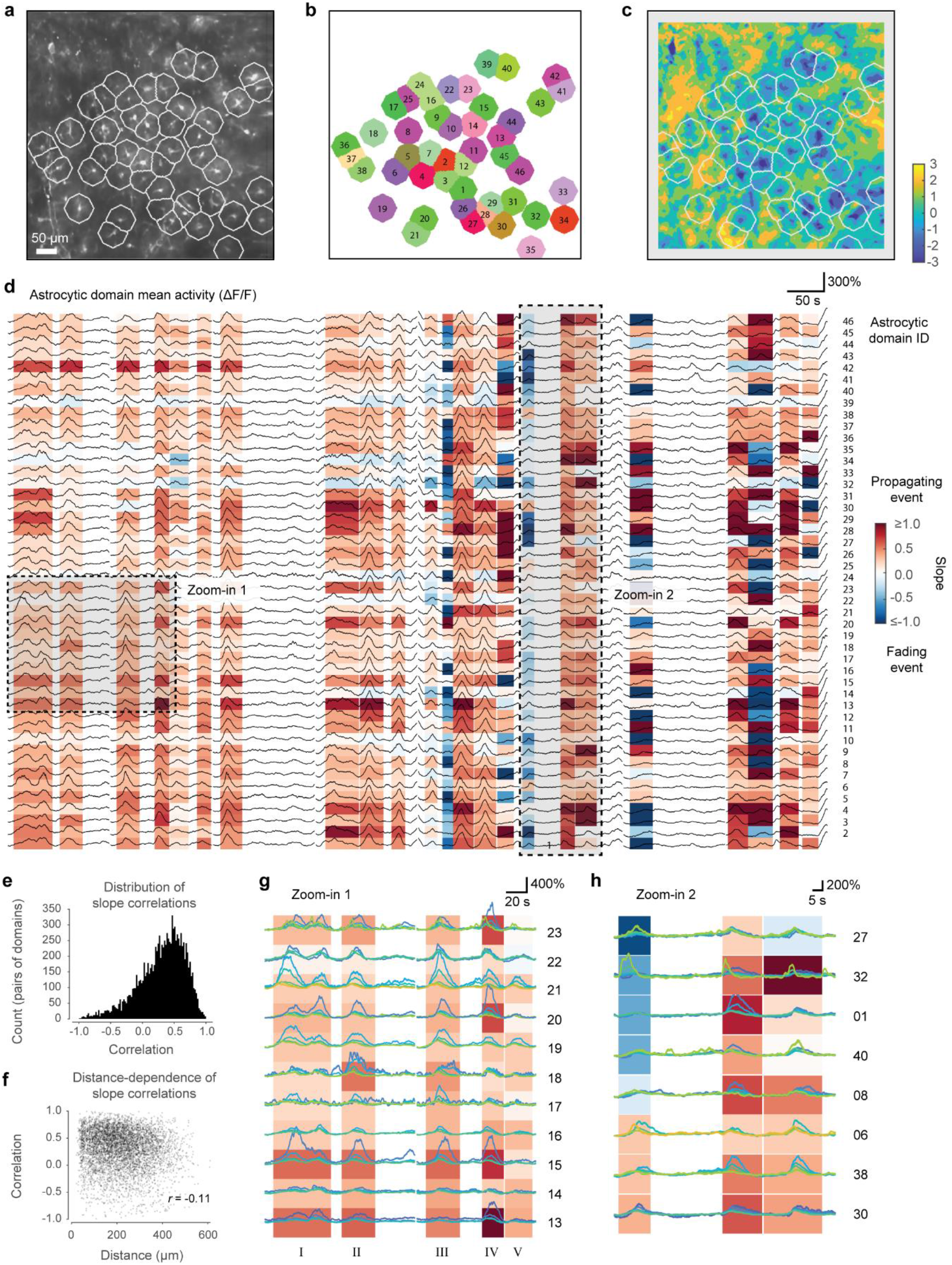
Centripetal propagation can be variable across different putative single astrocytic domains. Manual seeding of the astrocyte cell body centroid with subsequent watershed segmentation was used to create putative astrocytic domains (cell bodies identified based on average fluorescence in **(a)**, segmented domains with ID numbers in **(b)**). For each domain, the delay map **(c)** was used to extract delayed traces as shown in Fig. 6a-c, and a slope was fitted to the delayed traces for each detected event and each astrocytic domain, as described in Fig. 6c. Positive slope (blue) indicates an event propagating into the somatic region of the respective domain, a negative slope (red) indicates a fading event **(d)**. Overlaid traces display the average activity in the respective domain. **e**, Similarity of slopes across events for pairs of astrocytic domains quantified as correlation. The distribution is centered on positive values, showing that most astrocytic domains follow the propagation/fading of the majority of other domains. **f**, Distance-dependence of the pair-wise correlations of propagation slopes from (e), indicating a weak decay of the correlation with distance. However, low correlations occur also at neighboring domains (distance approx. 20-50 μm), and high correlations at large distances. **g**, Centripetal propagation appears more prominent in some astrocytic domains for some events. Zoom-in #1 from (d) into a subset of events (event numbers in roman numerals below) and astrocytic domains (numbers at the right side). In addition, delayed traces are shown (color-coding as in Fig. 6; from yellow = distal processes, to blue = astrocytic soma). In some domains, the somatic trace is activated much more than for other domains (*e*.*g*., domain 15) or for other events in other domains (*e*.*g*., domain 18 for event II, domain 20 for event IV, and domain 23 for event IV). This observation indicates that the strength of somatic activation by centripetal propagation is variable across astrocytes. **h**, Zoom-in #2 from (d), manually re-ordered selection of traces (numbers to the right indicate the putative astrocytic domain). The first event is the event of interest for the purpose of this panel. While the event is globally dominated by fading calcium signals as seen in (d), calcium activity in some domains clearly reaches the somatic regions (astrocytic domain IDs 30, 38 and 6). This observation suggests that centripetal propagation is a cell-autonomous process that can occur in a subset of astrocytes.

**Figure S20.**
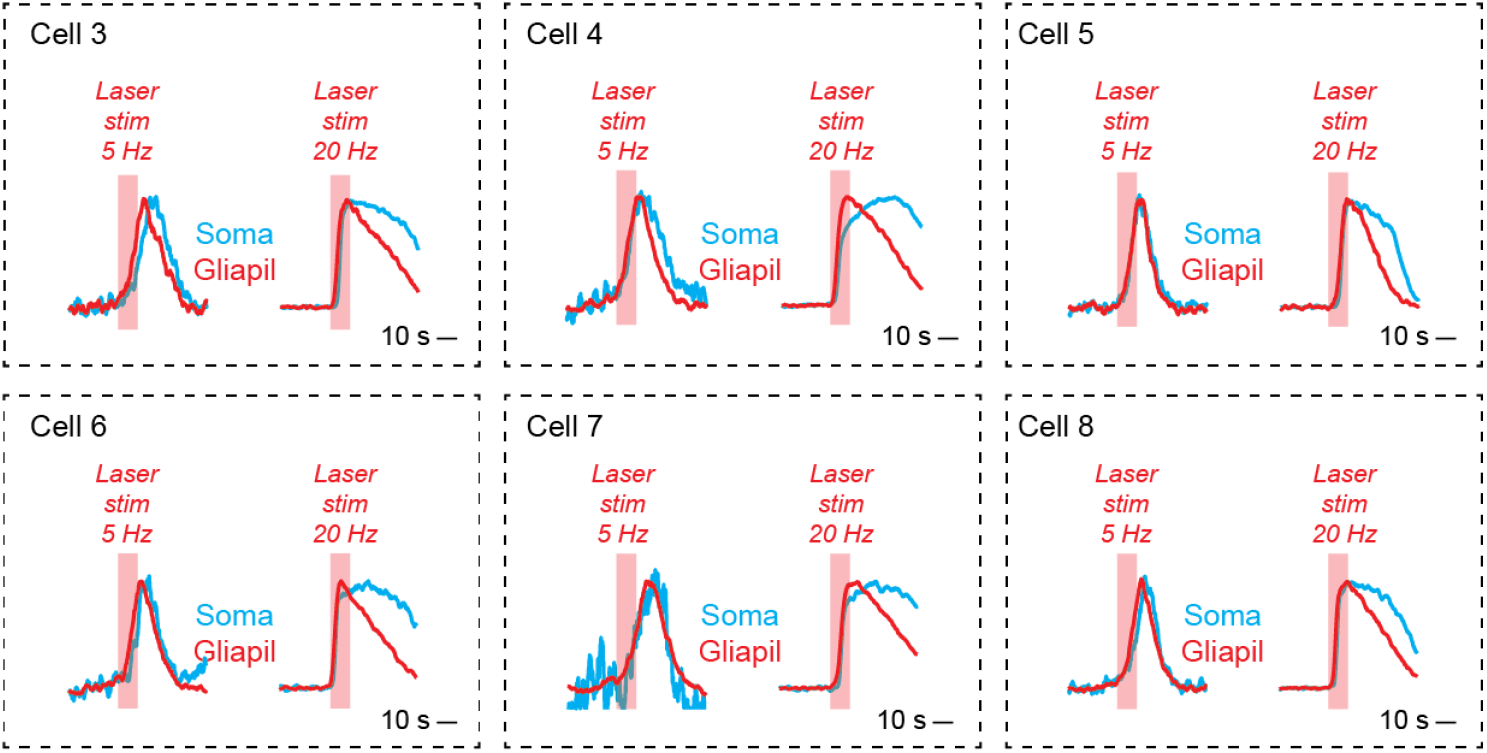
Persistent activation of astrocytic somata but not surrounding gliapil for strong stimulation of locus coeruleus, extension of Fig. 7n. For these astrocytes, somata (blue) were activated shortly after gliapil (red) for weaker stimuli (10 s stimulation, 5 Hz, see Methods for details). For stronger stimulation (10 s stimulation, 20 Hz), somatic calcium signals were not simply a delayed version of gliapil signals but showed a persistent component. Such persistent activation was also observed without optogenetic stimulation during spontaneous behavior (Fig. S18e,f).

**Figure S21.**
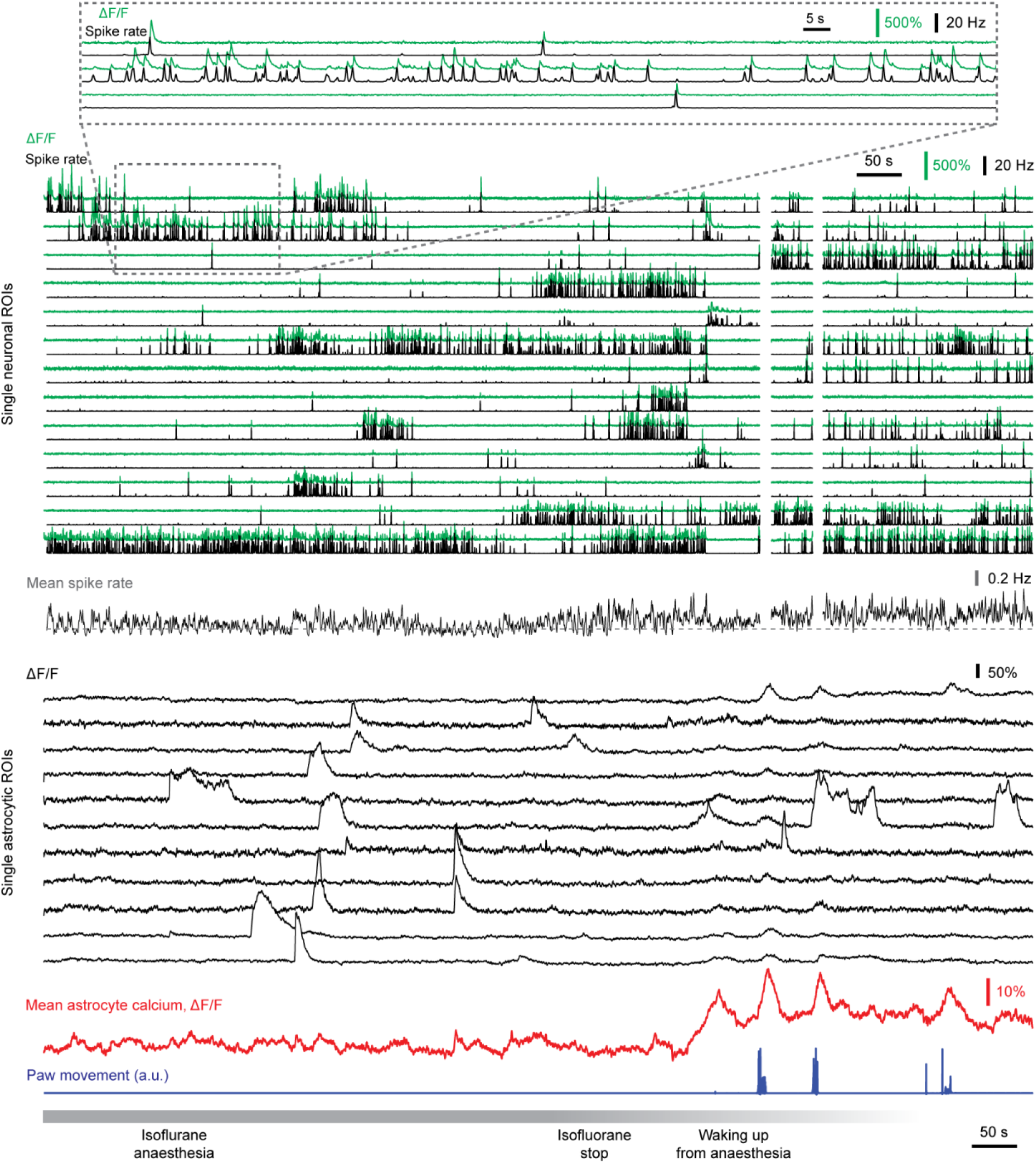
Calcium imaging of astrocytes and pyramidal neurons in CA1 during isoflurane anesthesia. Examples of extracted neuronal ΔF/F traces (green) and associated deconvolved spike rates (black) are shown (see also zoom-in at the top). White blanked time points were discarded due to excessive movement of the brain during the waking-up. Example astrocytic ΔF/F traces (black) are shown below, highlighting uncoordinated local but no global events before the waking up. Waking up (bottom; 1.5% in the beginning, set to 0% at “isoflurane stop”) is reflected by small and large paw movements (blue) and resulted in global astrocytic calcium signals (mean astrocytic calcium, red).

**Figure S22.**
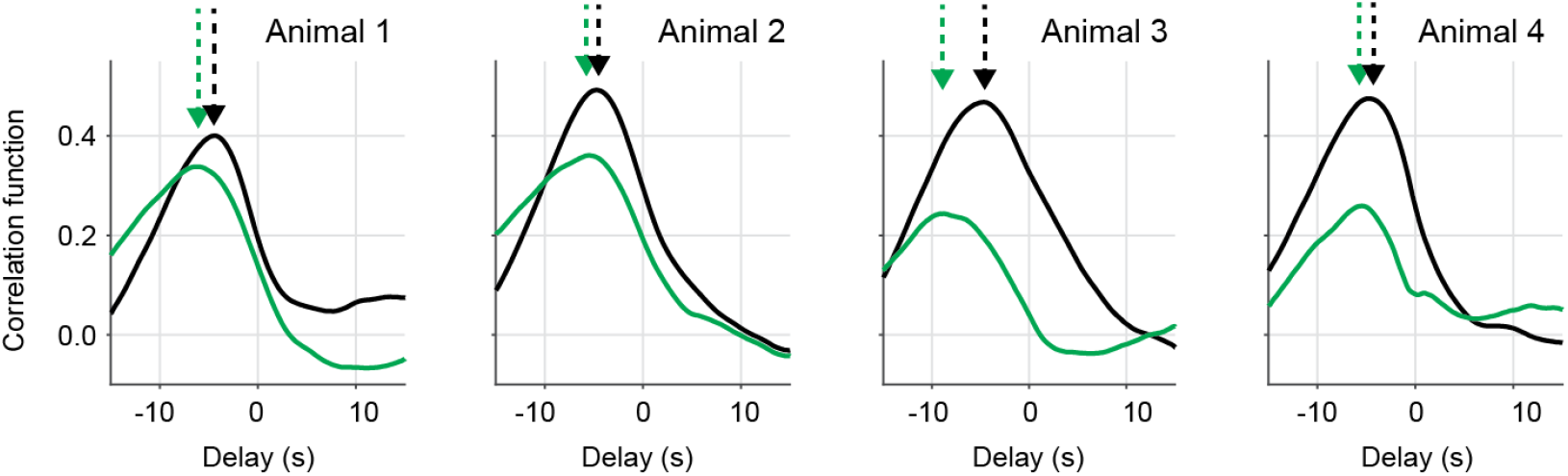
Correlation functions between global astrocytic activity and paw movement, extension of Fig. 8h. The peak of the correlation function indicates how much later global astrocytic activity occurred after paw movement. A longer delay indicates that the signaling pathways for global astrocytic signals and centripetal propagation were slower. Delays computed from correlation functions with prazosin injected-animals (green) were longer than delays from control animals (black) for all 4 animals. Control recordings were performed with injection of a vehicle, prazosin injections with injection of the same amount of solution with prazosin dissolved on the subsequent day. Each correlation function was computed from a recording session of 7 segment of 140 s each (980 s total). Extracted delays computed from all paired correlation functions (maximum value) are plotted in the box plot in Fig. 8h.

**Supplementary table 1.** Within- and across-animal variance for hierarchically acquired data. Details are described in the Statistics subsection of the Methods section. Intra-class-correlation (ICC) is defined as the across-animal variance divided by across-animal variance plus within-animal variance. “S.D.” indicates the standard deviation as a measure for variability.

|  | Variable | Median<br>across all<br>data points | S.D. across<br>all pairs | S.D. across<br>animals | S.D.<br>within<br>animals | ICC |
| --- | --- | --- | --- | --- | --- | --- |
| **Figure 1i:** Pairwise correlation | Correlation | 0.72 | 0.20 | 0.09 | 0.21 | 0.155172 |
| **Figure 1l:** Pairwise correlation | SO-SO | 0.5912 | 0.1642 | 0.0583 | 0.1339 | 0.159362 |
|  | SP-SP | 0.6099 | 0.1947 | 0.0793 | 0.0789 | 0.502528 |
|  | SR-SR | 0.6583 | 0.1333 | 0.0499 | 0.1564 | 0.09239 |
|  | SO-SP | 0.5731 | 0.1823 | 0.0172 | 0.0878 | 0.036958 |
|  | SO-SR | 0.5381 | 0.1640 | 0.1519 | 0.0972 | 0.709489 |
|  | SP-SR | 0.5735 | 0.1896 | 0.0020 | 0.1102 | 0.000329 |
| **Figure 3c:** Correlation | Pupil | 0.6147 | 0.1214 | 0.1223 | 0.1133 | 0.538145 |
|  | Paws | 0.3227 | 0.1255 | 0.0917 | 0.1268 | 0.34340 |
|  | Run | 0.2080 | 0.1069 | 0.0857 | 0.0851 | 0.503513 |
|  | SR | 0.2075 | 0.0967 | 0.0774 | 0.0770 | 0.502591 |
|  | Mouth | 0.2459 | 0.0962 | 0.0660 | 0.0750 | 0.436429 |
| | $\Delta F/F$ | 0.1693 | 0.1299 | 0.1259 | 0.0847 | 0.688420 |
|  | Licking | 0.0355 | 0.0580 | 0.0373 | 0.0469 | 0.387449 |
|  | Location | 0.0136 | 0.1114 | 0.0206 | 0.0587 | 0.109652 |
| **Figure 3d:** Multi-timepoint regression | Pupil | 0.6069 | 0.1544 | 0.0974 | 0.1557 | 0.281262 |
|  | Paws | 0.5997 | 0.1687 | 0.0764 | 0.1711 | 0.166238 |
|  | Running | 0.4596 | 0.1734 | 0.1070 | 0.1666 | 0.292032 |
|  | SR | 0.4498 | 0.2011 | 0.0858 | 0.2017 | 0.153225 |
|  | Mouth | 0.4415 | 0.1741 | 0.1435 | 0.1504 | 0.476536 |
| | $\Delta F/F$ | 0.2915 | 0.1856 | 0.1379 | 0.1693 | 0.398843 |
|  | Licking | 0.1381 | 0.2195 | 0.1416 | 0.1661 | 0.420879 |
|  | Location | 0.2067 | 0.1296 | 0.0717 | 0.0656 | 0.544341 |
| **Figure 3h:** Future and past | Pupil past | 0.5983 | 0.1507 | 0.1052 | 0.1505 | 0.32823 |
|  | Paws past | 0.6051 | 0.1729 | 0.0876 | 0.1670 | 0.215781 |
|  | SR past | 0.4682 | 0.1789 | 0.1068 | 0.1684 | 0.286843 |
|  | Pupil future | 0.5669 | 0.1670 | 0.1353 | 0.1638 | 0.405571 |
|  | Paws future | 0.3243 | 0.1922 | 0.1517 | 0.1893 | 0.391061 |
|  | SR future | 0.1739 | 0.1777 | 0.1451 | 0.1455 | 0.498624 |
| **Figure 3j:** DGL | Pupil DGL | 0.5809 | 0.1416 | 0.1249 | 0.1203 | 0.518754 |
|  | Paws DGL | 0.6480 | 0.1488 | 0.0999 | 0.1422 | 0.330455 |
|  | SR DGL | 0.5360 | 0.1627 | 0.0856 | 0.1517 | 0.241506 |
| **Figure 4:** Delays | Neurons spike rate | 4.2167 | 3.3913 | 1.22240 | 2.8253 | 0.157679 |
|  | Paw movement | 3.6667 | 1.2760 | 0.76277 | 0.94236 | 0.395832 |
|  | Licking | 3.6667 | 3.2866 | 0.99943 | 1.2708 | 0.382150 |
|  | Mouth movement | 3.6333 | 1.5115 | 1.66980 | 1.3221 | 0.614665 |
|  | Run speed | 3.3667 | 1.5704 | 1.25090 | 1.4083 | 0.441016 |
|  | Pupil diameter | 1.5000 | 1.0573 | 0.78489 | 0.74105 | 0.528706 |
| **Figure 8:** Spike rates | Awake (Hz) | 0.14081 | 0.031215 | 0.020572 | 0.026125 | 0.382743 |
|  | Anesthetized (Hz) | 0.0930 | 0.0110 | 0.0110 | - | 0 |

## Movie captions

**Movie 1** | **Single-plane calcium imaging of hippocampal astrocytes *in vivo***. Calcium imaging of the calcium reporter GCaMP6s in the *stratum oriens* of hippocampal CA1 in a behaving mouse through an implanted cannula. The raw movie was denoised using a self-consistent denoising algorithm (DeepInterpolation, see Methods). For improved display of dim astrocytic processes, some of the brighter areas in the field of view are saturated. The field of view is 600 x 600 μm^2^, the original frame rate 30.88 Hz (displayed at 10x speed).

**Movie 2** | **Triple-layer calcium imaging of hippocampal astrocytes *in vivo***. Calcium imaging of the calcium indicator GcaMP6s in three layers of hippocampal CA1 (left: *stratum oriens*, middle: *stratum pyramidale*, right: *stratum radiatum)* in a behaving mouse through an implanted cannula using a tunable lens for fast z-scanning. The raw movie was denoised using a self-consistent denoising algorithm (DeepInterpolation, see Methods). The original field of view for each plane is 200 x 200 μm^2^, the original frame rate 10.29 Hz (displayed at 12x speed).

**Movie 3** | **Simultaneous monitoring of astrocytic and neuronal activity, pupil changes and body movement**. The extracted traces are smoothed with a 5-point moving window and z-scored for clarity. The x-axis indicates time in seconds. The movie is displayed at real-time speed. Related to Figure 2.

**Movie 4** | **Examples of mouth movement without detectable effect on astrocytes**. The extracted traces are smoothed with a 5-point moving window and z-scored for clarity. The x-axis indicates time in seconds. The movie is displayed at 2x speed. Related to Figure 3.

**Movie 5** | **Examples of front limb movements (reaching and grooming) and its effect on astrocytic activity**. The extracted traces are smoothed with a 5-point moving window and z-scored for readability. The x-axis indicates time in seconds. The movie is displayed at 2x speed. Related to Figure S9.

**Movie 6** | **Self-consistent denoising of astrocytic calcium movies using DeepInterpolation**. In the first part of the movie, the left version shows 140 s of a raw astrocytic calcium recording from hippocampal CA1 (*stratum oriens*). The right version shows the movie denoised using DeepInterpolation (see Methods). In the second part of the movie, the raw calcium movie is again shown together with the denoised version, but with the raw movie smoothed using a Gaussian filter (standard deviation 10 time points), resulting in smoothed time courses but visibly less efficient denoising compared to the self-consistent denoising method. Related to Figures 5 and S11.

**Movie 7** | **Optogenetic stimulation of locus coeruleus during two-photon calcium imaging in hippocampal astrocytes**. Related to Figure 7. Calcium imaging of the calcium reporter GCaMP6s in the *stratum oriens* of hippocampal CA1 in a behaving mouse. The yellow trace indicates the mean fluorescence across the FOV, the dot indicates the current time. The recording consists of multiple segments with 5-20 s between segments, with segment boundaries indicated by interrupted yellow traces. The raw movie was denoised using a self-consistent denoising algorithm (DeepInterpolation, see Methods). The field of view is 650 x 650 μm^2^, the original frame rate 30.88 Hz (displayed at increased speed).

**Movie 8** | **Stimulus-triggered average for optogenetic activation of locus coeruleus during two-photon calcium imaging in hippocampal astrocytes**. Stimulus-triggered averages from the data underlying Movie 7, related to Figure 7. First part: raw fluorescence. Second part: Average fluorescence, overlaid with ∆F/F in yellow. Third part: Zoom-in to three selected astrocytes. Before stimulus-triggered averaging, the raw movie was denoised using a self-consistent denoising algorithm (DeepInterpolation, see Methods). The field of view is 650 x 650 μm^2^, the original frame rate 30.88 Hz (displayed at increased speed).

## Notes

https://github.com/HelmchenLabSoftware/Centripetal_propagation_astrocytes

### Competing Interest Statement

The authors have declared no competing interest.

### Summary of Updates

New experiments and analyses have been added. In particular, figures and text describing optogenetic perturbations of the locus coeruleus during two-photon calcium imaging or fiber photometry (new Fig. 7) and pharamacological perturbation of noradrenergic signaling (new Fig. 8). Eight additional supplementary figures have been addded, and further supplementary figures have been substantially enhanced. New authors, who contributed to the new experiments, were added to the author list (Sian Duss, Denise Becker, Johannes Bohacek).

